# Divergent evolution of early terrestrial fungi reveals the evolution of Mucormycosis pathogenicity factors

**DOI:** 10.1101/2022.06.24.497490

**Authors:** Yan Wang, Ying Chang, Jericho Ortanez, Jesús F. Peña, Derreck Carter-House, Nicole K Reynolds, Matthew E Smith, Gerald Benny, Stephen J Mondo, Asaf Salamov, Anna Lipzen, Jasmyn Pangilinan, Jie Guo, Kurt LaButti, William Andreopolous, Andrew Tritt, Keykhosrow Keymanesh, Mi Yan, Kerrie Barry, Igor V Grigoriev, Joseph W Spatafora, Jason E Stajich

## Abstract

Fungi have evolved over millions of years and their species diversity is predicted to be the second largest on the earth. Fungi have cross-kingdom interactions with many organisms which have mutually shaped their evolutionary trajectories. Zygomycete fungi hold a pivotal position in the fungal tree of life and provide important perspectives on the early evolution of fungi from aquatic to terrestrial environments. Phylogenomic analyses have found that zygomycete fungi diversified into two separate clades, the Mucoromycota which are frequently associated with plants and Zoopagomycota that are commonly animal-associated fungi. Genetic elements that contributed to the fitness and divergence of these lineages may have been shaped by the varied interactions these fungi have had with plants, animals, bacteria and other microbes. To investigate this, we performed comparative genomic analyses of the two clades in the context of Kingdom Fungi, benefiting from our generation of a new collection of zygomycete genomes. We identified lineage-specific genomic content which may contribute to the disparate biology observed in these zygomycetes. Our findings include the discovery of undescribed diversity in CotH, a Mucormycosis pathogenicity factor, which was found in a broad set of zygomycetes. Reconciliation analysis identified multiple duplication events and an expansion of CotH copies throughout Mucoromycotina, Mortierellomycotina, Neocallimastigomycota, and *Basidiobolus* lineages. A kingdom-level phylogenomic analysis also identified new evolutionary relationships within the sub-phyla of Mucoromycota and Zoopagomycota.

## INTRODUCTION

Fungi play diverse ecological roles and interact with various organisms in both terrestrial and aquatic environments (James, Kauff, et al. 2006; Stajich et al. 2009; Spatafora et al. 2017; Fisher et al. 2020). Since their divergence from a common ancestor with animals over 1 billion years ago, fungi have evolved complex relationships with other organisms, including animals, bacteria, plants, protists, and other fungi (Currie et al. 2003; Frey-Klett et al. 2011; Parfrey et al. 2011; Gruninger et al. 2014; Uehling et al. 2017; Wang et al. 2018; Chambouvet et al. 2019; Malar et al. 2021). As a distinct eukaryotic kingdom, Fungi are characterized by chitinous cell walls and osmotrophic feeding style, although neither of these characters is diagnostic for the kingdom (Richards et al. 2017; James et al. 2020). The versatile enzymes secreted by fungi facilitate their success in utilization of diverse polysaccharides and are key members of ecosystems supporting nutrient cycling processes (Hori et al. 2013; Chang et al. 2015; Solomon et al. 2016; Richards and Talbot 2018; Chang et al. 2022). Zygomycete fungi are a historically enigmatic group as their diversity and phylogenetic placement on the fungal tree of life remained somewhat cryptic based on morphological characters alone. The lineages emergence coincide with major transition of fungi from aquatic environment to terrestrial ecologies, which was characterized by the evolutionary loss of the flagellum (James, Letcher, et al. 2006; James, Kauff, et al. 2006; Chang et al. 2021). The zygomycete fungi are recognized by their gametangial conjugation, production of zygospore, and coenocytic aseptate or septate hyphae (White et al. 2006; Hibbett et al. 2007; Spatafora et al. 2017; Naranjo-Ortiz and Gabaldón 2020). Nevertheless, zygospore structures have not been observed for most members of zygomycete fungi due to their cryptic sexual stage or lack of appropriate culture approaches. Zygomycete fungi were found to be paraphyletic based on genome-scale evidence, as a result, two new phyla (Mucoromycota and Zoopagomycota) were established to accommodate the current members (Spatafora et al. 2016). However, incomplete sampling of zygomycete lineages has made resolution of the origin of terrestrial fungi difficult to resolve with standard phylogenetic approaches (Chang et al. 2021; Li et al. 2021).

Mycological and fungal cell biology research has been historically biased in favor of members of the Dikarya. Several established research model organisms have advanced fields of cell biology including the brewer’s yeast *Saccharomyces cerevisiae*, the fission yeast *Schizosaccharomyces pombe*, the red bread mold *Neurospora crassa*, and the filamentous mold *Aspergillus nidulans*. These model organisms contributed to an expansion in the understanding of eukaryotes. Fungi were among the some of the first sequenced eukaryotic genomes (Goffeau et al. 1996; Wood et al. 2002; Galagan et al. 2003; Galagan et al. 2005). However, genomic research on zygomycete fungi had to wait for the first Mucoromycotina genome to be sequenced in 2009 (Ma et al. 2009). The majority of our existing knowledge of zygomycetes has come from studies of arbuscular mycorrhizae (Glomeromycotina) or saprophytes classified in Mucoromycota, such as the black bread mold *Rhizopus stolonifer*. Studies on the other zygomycetes phylum, Zoopagomycota, are still rare, and the biodiversity of Zoopagomycota fungi is likely greatly underestimated and the research progress is largely hindered by the lack of axenic cultures. Culture independent studies have identified multiple zygomycetes as amplicon-based operational taxonomic units (OTUs) in unexplored ecological sites (Metcalf et al. 2016; Picard 2017; Pombubpa et al. 2020; Reynolds et al. 2021) and many “unknown” fungal OTUs will likely to be identified with the help of increasing fungal genomes, especially more representatives in the sparsely sequenced zygomycete lineages.

To fill this gap, our recent emphasis on sequencing zygomycete genomes through the ZyGoLife project (Spatafora et al. 2016; https://zygolife.org) have produced over a hundred genomes. The output has become the largest collection of genomic information for this fungal clade. Various techniques were also developed and employed to obtain genome sequences of the uncultured zygomycete species. The breakthroughs include the single-cell genomics as well as fungus-host co-culture techniques (Ahrendt et al. 2018) and sequencing of metagenomes of sporocarps (Chang et al. 2019). Progress on genomics and related multi-omics have greatly expanded our knowledge on zygomycetes. This includes the identification of a mosquito-like polyubiquitin gene in a zygomycete fungus inhabiting the gut of mosquitoes (*Zancudomyces culisetae*, Zoopagomycota) (Wang et al. 2016), the discovery of a photosynthetic mycelium using algal symbionts (*Linnemannia elongata*, Mortierellomycotina) (Du et al. 2019; Vandepol et al. 2020), the isolation of cicada behavior modifying alkaloids from *Massospora* (Entomophthoromycotina) (Boyce et al. 2019), and the expansion of secondary metabolite genes of amphibian gut fungi (*Basidiobolus*, Entomophthoromycotina) via Horizontal Gene Transfer from bacteria co-existing in the gastrointestinal tract (Tabima et al. 2020). However, a conundrum remains as to the evolutionary history of the zygomycete fungi. What evolutionary processes were associated with the divergence of the ancestors of Mucoromycota and Zoopagomycota into species which primarily associate with plants and plant material or animal and fungal hosts, respectively. We hypothesize that comparisons of gene content will enable identification of genetic elements that have contributed to their success in these ecologies and their reproductive strategies and may be reflected in lineage-specific genes, those with expanded copy number or enrichment in specific pathways or processes that underpin adaptations to these hosts and environments. In addition, the construction of a well-resolved phylogenetic tree incorporating the expanded collection of zygomycete genomes is an important framework to consider the complex natural history and relationships among these diverse fungi. Our work has contributed to the generation of 131 recent zygomycete genomes (Table 1), which were used to investigate the evolution and cryptic genetics behind the biology of these early-diverging fungi.

**Table 1.**
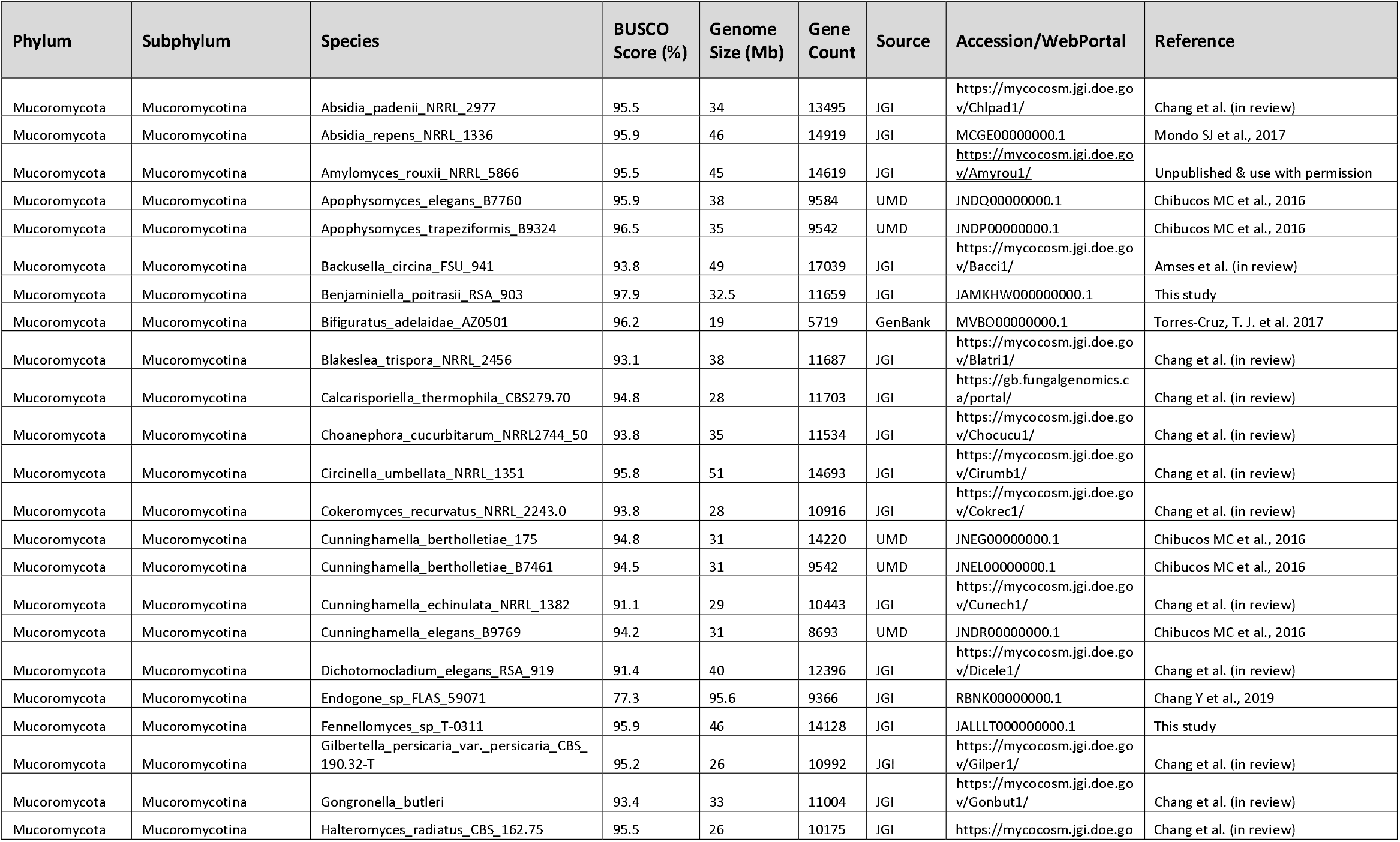
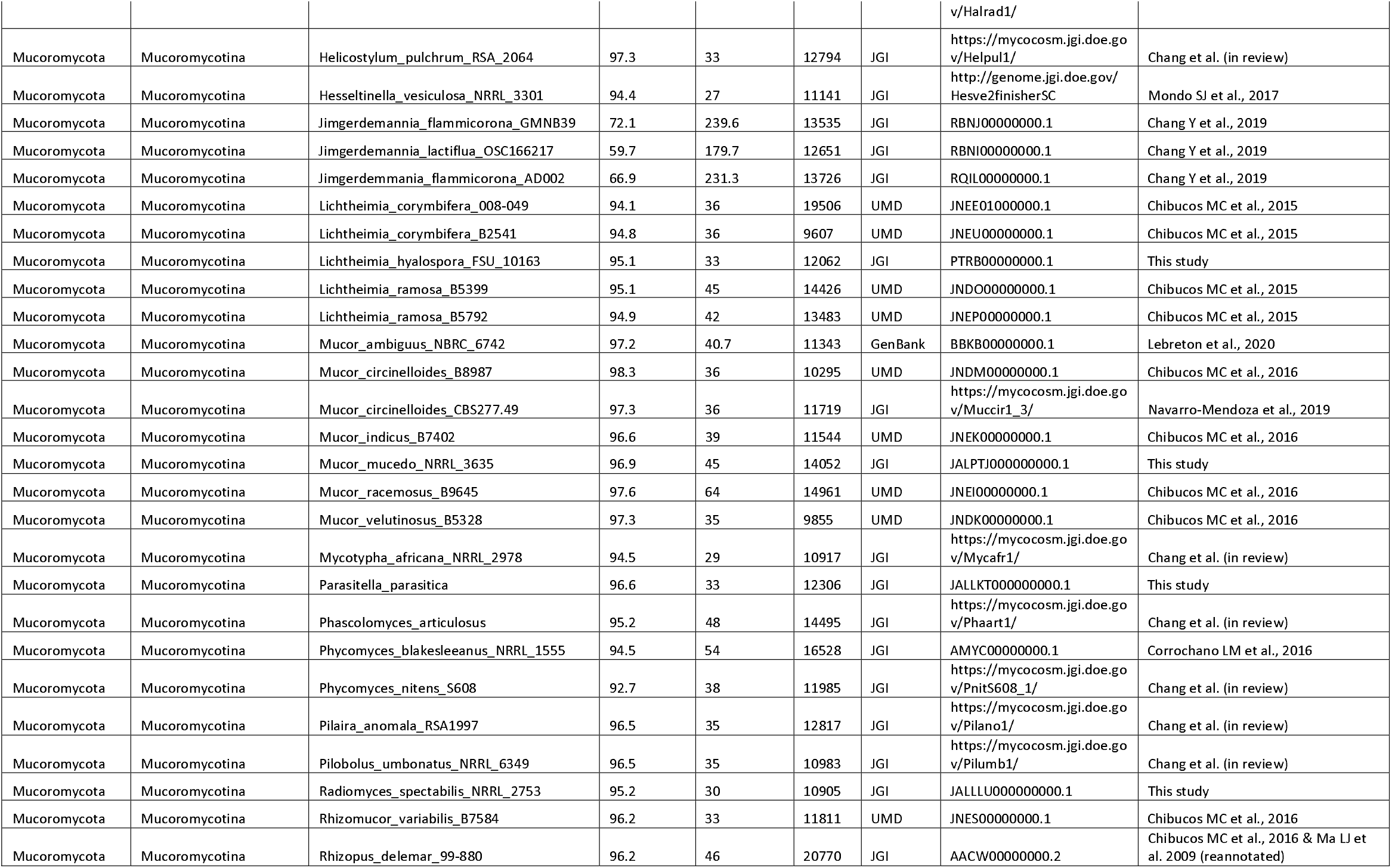
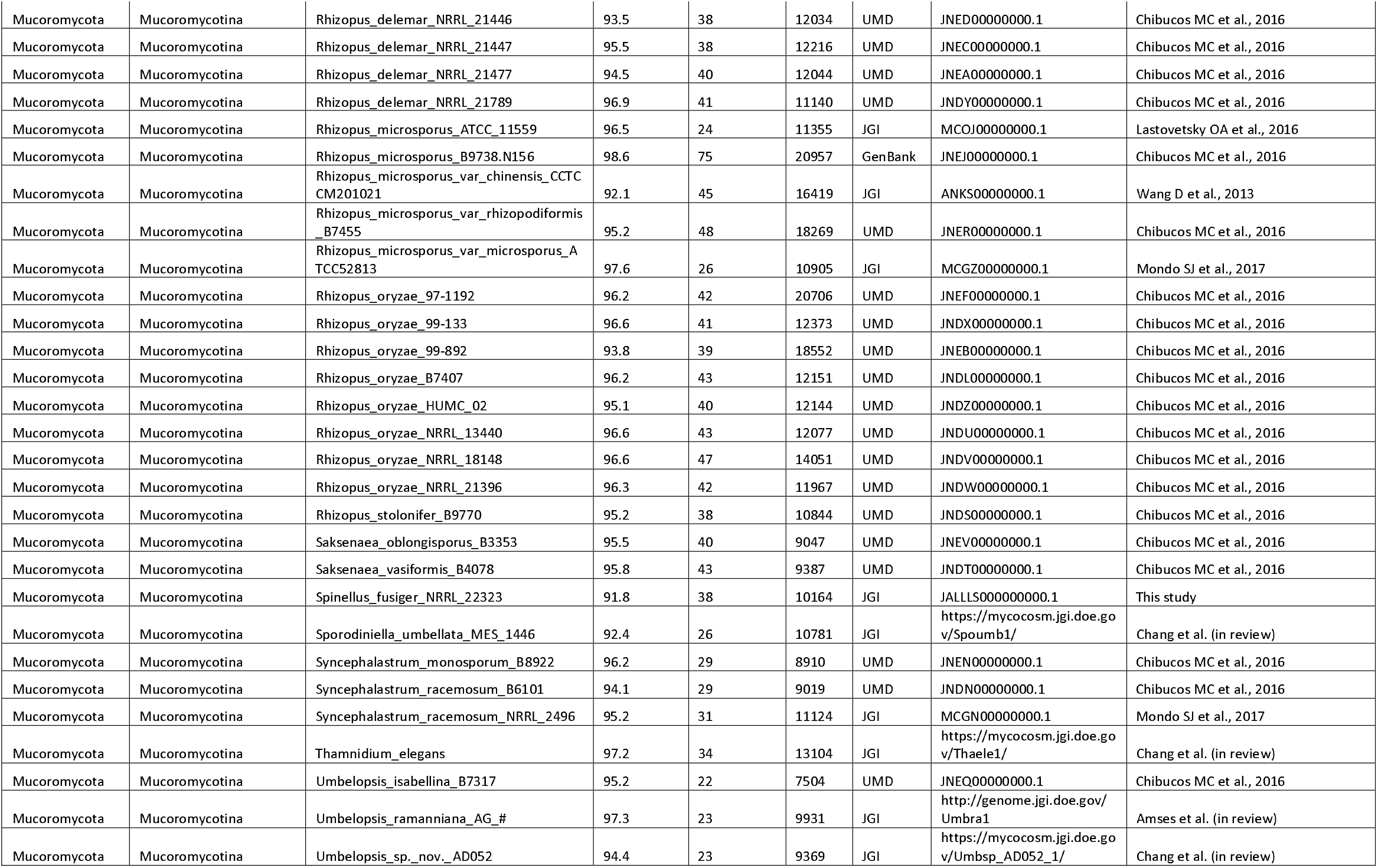
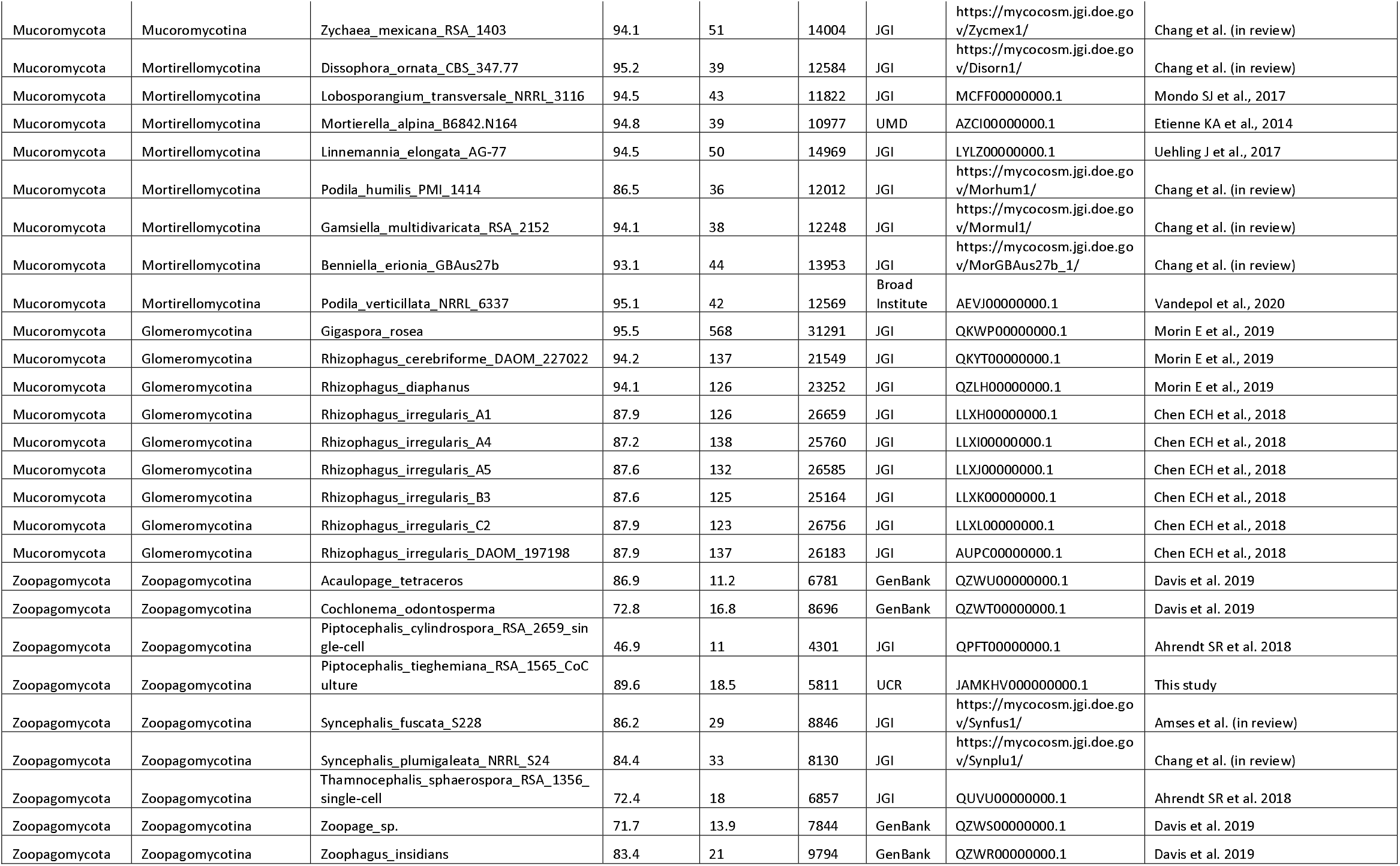
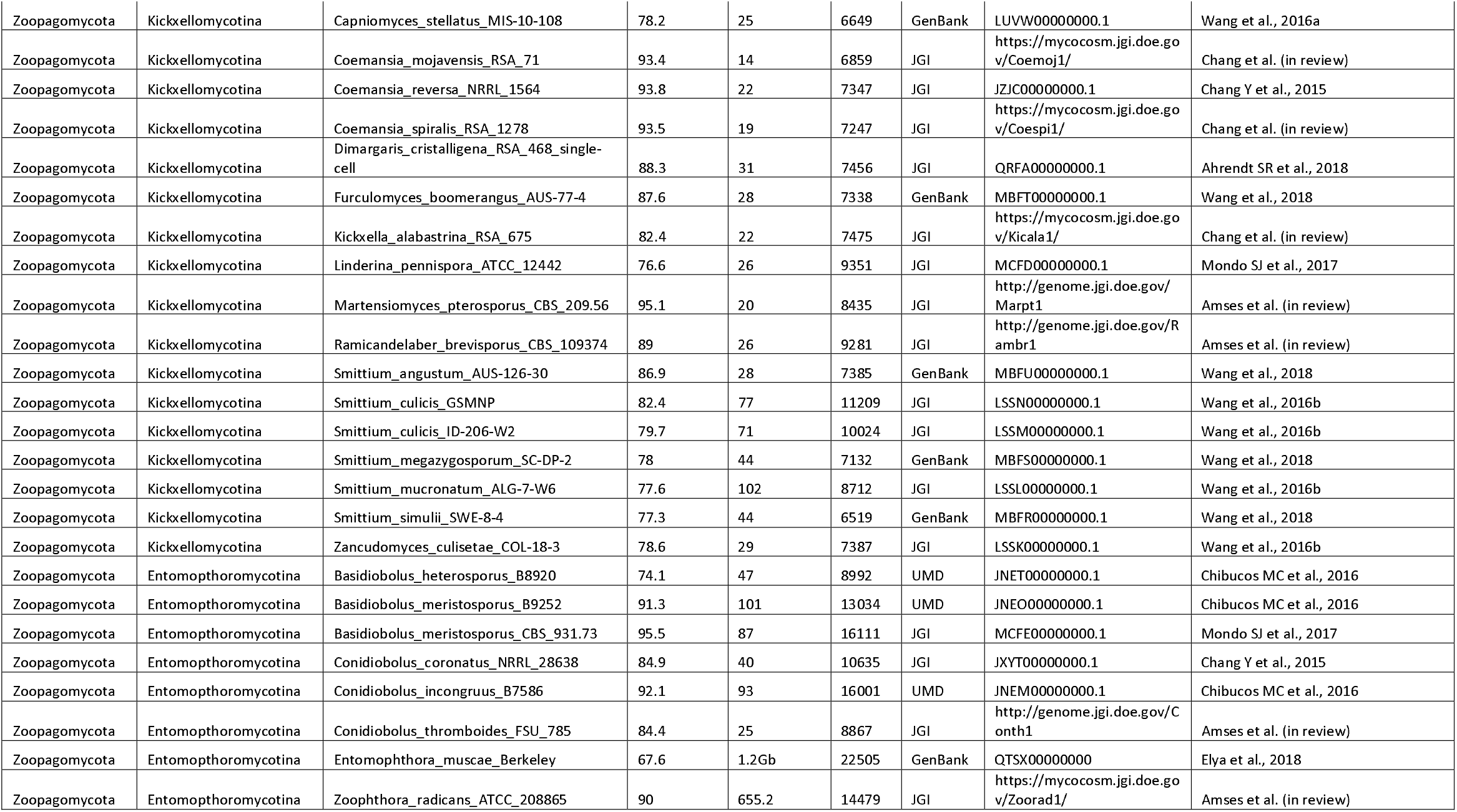
List of 131 zygomycete fungi included in the current phylogenetic and comparative genomics study

The focus on these phyla is motivated by not only understanding their ecological roles and history, but also in the context of the increase in Mucormycosis, a deadly human-infectious disease, that has risen in prevalence and public attention due to high infection rates and co-morbidity during the COVID-19 pandemic (Garg et al. 2021; Revannavar et al. 2021). Mucormycosis is caused by members of Mucoromycotina, in particular many genera of the Mucorales fungi (Soare et al. 2020). We cataloged the prevalence of Mucormycosis pathogenicity factors across Mucorales genomes and profiled their evolutionary conservation among members of the Fungal Kingdom. We identified the genes for the Mucormycosis invasin factor in three Mortierellomycotina species as well (*Dissophora ornate, Lobosporangium transversle*, and *Mortierella* species) which all share a highly similar protein motif associated with the disease in Mucorales fungi indicating these fungi may have additional potential for mammalian infection and the more ancient nature of this factor within these fungi. Our study highlights the importance of research on zygomycetes to characterize the unique and shared molecular components of their biology that can be examined as more genome sequences become available. Our improved resolution phylogeny will enhance the study of the evolutionary relationships for both organismal and molecular genetics of these important fungi.

## MATERIALS AND METHODS

### Fungal taxa and genome sampling

In total, 181 fungal genome sequences were analyzed in this study. Nine genomes were generated in this study and 172 were obtained from GenBank or the Joint Genome Institute MycoCosm portal (Grigoriev et al. 2014; https://mycocosm.jgi.doe.gov), with 136 produced by the ongoing 1000 Fungal Genome Project (1KFG: http://1000.fungalgenomes.org/) and Zygomycetes Genealogy of Life Project (ZyGoLife: http://zygolife.org/). The dataset includes 131 zygomycetes genomes (Table 1), with 97 sampled from Mucoromycota clade and 34 from Zoopagomycota. In addition, we included 43 Dikarya genomes and seven representatives (Supplementary Table 1) from other early-diverging fungal lineages to enable kingdom-wide comparative analyses. The following nine genomes were produced for this study: *Amylomyces rouxii* NRRL 5866, *Benjaminiella poitrasii* RSA 903, *Fennellomyces* sp. ATCC 46495, *Lichtheimia hyalospora* FSU 10163, *Mucor mucedo* NRRL 3635, *Parasitella parasitica* NRRL 2501, *Radiomyces spectabilis* NRRL 2753, *Spinellus fusiger* NRRL 22323, *Piptocephalis tieghemiana* RSA 1565.

### Genome sequencing and assembly

The genome sequencing of *Spinellus fusiger* NRRL 22323, *Radiomyces spectabilis* NRRL 2753, *Mucor mucedo* NRRL 3636, *Benjaminiella poitrasii* RSA 903 and *Fennellomyces* sp. ATCC 46495, was performed from 5 ug of genomic DNA was sheared to >10kb using Covaris g-Tubes. The sheared DNA was treated with exonuclease to remove single-stranded ends and DNA damage repair mix followed by end repair and ligation of blunt adapters using SMRTbell Template Prep Kit 1.0 (Pacific Biosciences). The library was purified with AMPure PB beads. PacBio Sequencing primer was then annealed to the SMRTbell template library and Version P6 sequencing polymerase was bound to them for *S. fusiger, R. spectabilis* and *Fennellomyces* sp. ATCC 46495. The prepared SMRTbell template libraries were then sequenced on a Pacific Biosciences RSII sequencer using Version C4 chemistry and 1×240 sequencing movie run times. For *B. poitrasii* and *M. mucedo*, sequencing polymerase was bound to them using the Sequel Binding kit 2.1 and then the prepared SMRTbell template libraries were sequenced on a Pacific Biosystems’ Sequel sequencer using v3 sequencing primer, 1M v2 SMRT cells, and Version 2.1 sequencing chemistry with 1×360 sequencing movie run times. Filtered subread data was then used to assemble all lineages using Falcon (version 0.4.2 for *S. fusiger* and *R. spectabilis*, version 1.8.8 for *M. mucedo* and *B. poitrasii*, and version 0.7.3 for *Fennellomyces* sp. ATCC 46495). *S. fusiger and R. spectabilis* were then further improved using finisherSC version 2.0 (Lam et al. 2015). All assemblies were then polished using either Quiver version smrtanalysis_2.3.0.140936.p5 (*S. fusiger, R. spectabilis* and *Fennellomyces* sp. ATCC 46495) or Arrow version SMRTLink v5.1.0.26412 (*M. mucedo* and *B. poitrasii*).

*Parasitella parasitica* NRRL 2501, *Piptocephalis tieghemiana* and *Lichtheimia hyalospora* were sequenced using the Illumina platform. For *P. parasitica* and *P. tieghemania*, 100 ng of DNA was sheared to 300 bp using the Covaris LE220 and size selected using SPRI beads (Beckman Coulter). The fragments were treated with end-repair, A-tailing, and ligation of Illumina compatible adapters (IDT, Inc) using the KAPA-Illumina library creation kit (KAPA biosystems). Additionally, a 4kb mate pair library was constructed for *P. parasitica*. For this, 5-10 ug of DNA was sheared using the Covaris g-TUBE(TM) and gel size selected for 4 kb. The sheared DNA was treated with end repair and ligated with biotinylated adapters containing loxP. The adapter ligated DNA fragments were circularized via recombination by a Cre excision reaction (NEB). The circularized DNA templates were then randomly sheared using the Covaris LE220 (Covaris). The sheared fragments were treated with end repair and A-tailing using the KAPA-Illumina library creation kit (KAPA biosystems) followed by immobilization of mate pair fragments on strepavidin beads (Invitrogen). Illumina compatible adapters (IDT, Inc) were ligated to the mate pair fragments and 8 cycles of PCR was used to enrich for the final library (KAPA Biosystems). The prepared libraries were quantified using KAPA Biosystems’ next-generation sequencing library qPCR kit and run on a Roche LightCycler 480 real-time PCR instrument. The quantified libraries were then prepared for sequencing on the Illumina HiSeq sequencing platform utilizing a TruSeq paired-end cluster kit, v4. Sequencing of the flowcell was performed on the Illumina HiSeq2500 sequencer using HiSeq TruSeq SBS sequencing kits, v4, following a 2×150 indexed run recipe. Each fastq file was QC filtered for artifact/process contamination and subsequently assembled together with AllPathsLG version R49403 (Gnerre et al. 2011).

Since *P. tieghemania* is an obligate mycoparasite, it was maintained as co-culture with *Umbelopsis* sp. nov. AD052. The *P. tieghemania* contigs required further processing to separate these two assemblies. First, metagenomic scaffold sequences were binned into two groups using metabat (v2.12.1). The filtered reads were mapped to the sequences of the two bins and split into two separate datasets corresponding to each bin using bbsplit.sh in bbtools(ambiguous=all). The two datasets were then re-assembled separately. Scaffolds with length less than 2kb were excluded. Then, four closely related genomes were used for reference genome to classify and filter re-assembled scaffolds based on BLASTN similarity (evalue < 1e-30). One included *Piptocephalis* related genome, *Piptocephalis cylindrospora*, and the others were *Umbelopsis* related genomes, *Umbelopsis* sp. AD052, *Umbelopsis isabellina* AD026 and *Umbelopsis* sp. PMI 123. If the scaffolds were covered more by *Piptocephalis* main genome than *Umbelopsis* main genomes, it would be classified to *Piptocephalis tieghemiana*, and vice versa. The scaffolds without any similarity to the four genomes were discarded.

For *L. hyalospora*, 500 ng of DNA was sheared to 270 bp using the Covaris E210 (Covaris, Woburn, MA) and size selected using SPRI beads (Beckman Coulter, Brea, CA). The fragments were treated with end-repair, A-tailing, and ligation of Illumina adapters using the TruSeq Sample Prep Kit (Illumina, San Diego, CA), followed by quantification of libraries using KAPA Biosystem’s next generation sequencing library qPCR kit and run on a Roche LightCycler 480 real-time PCR instrument. The quantified libraries were multiplexed and the pools were then prepared for sequencing on the Illumina HiSeq sequencing platform utilizing a TruSeq paired-end cluster kit, v3, and Illumina’s cBot instrument to generate a clustered flowcell for sequencing. Sequencing of the flowcell was performed on the Illumina HiSeq2000 sequencer using a TruSeq SBS sequencing kit 200 cycles, v3, following a 2×150 indexed run recipe. Genomic reads were QC filtered for artifact/process contamination and subsequently assembled with Velvet. The resulting assembly was used to create a simulated 3 Kbp insert long mate-pair library, which was then assembled together with the original Illumina library with AllPathsLG release version R42328.

### Transcriptome sequencing and assembly

For all lineages except *L. hyalospora*, Stranded cDNA libraries were generated using the Illumina Truseq Stranded RNA LT kit. mRNA was purified from 1 ug of total RNA using magnetic beads containing poly-T oligos. mRNA was fragmented and reversed transcribed using random hexamers and SSII (Invitrogen) followed by second strand synthesis. The fragmented cDNA was treated with end-pair, A-tailing, adapter ligation, and 8 cycles of PCR. For *L. hyalospora*, Plate-based RNA sample prep was performed on the PerkinElmer Sciclone NGS robotic liquid handling system using Illumina’s TruSeq Stranded mRNA HT sample prep kit utilizing poly-A selection of mRNA following the protocol outlined by Illumina in their user guide: https://support.illumina.com/sequencing/sequencing_kits/truseq-stranded-mrna.html, and with the following conditions: total RNA starting material was 1 ug per sample and 8 cycles of PCR was used for library amplification. The prepared libraries were then quantified using KAPA Biosystems’ next-generation sequencing library qPCR kit and run on a Roche LightCycler 480 real-time PCR instrument. The quantified libraries were then prepared for sequencing on the Illumina HiSeq sequencing platform utilizing a TruSeq paired-end cluster kit, v4. Sequencing of the flowcell was performed on the Illumina HiSeq2500 sequencer using HiSeq TruSeq SBS sequencing kits, v4, following a 2×150 indexed run recipe (2×100 for *L. hyalospora*). Filtered fastq files were used as input for de novo assembly of RNA contigs. For all lineages except *L. hyalospora* and *P. parasitica*, reads were assembled into consensus sequences using Trinity version 2.1.1. Trinity was run with the --normalize_reads (In-silico normalization routine) and --jaccard_clip (Minimizing fusion transcripts derived from gene dense genomes) options. For *L. hyalospora* and *P. parasitica*, Rnnotator version 2.5.6 or later was used. *P. parasitica* was further improved using eight runs of velveth (v. 1.2.07) performed in parallel, once for each hash length for the De Bruijn graph. Minimum contig length was set at 100. The read depth minimum was set to 3 reads. Redundant contigs were removed using Vmatch (v. 2.2.4) and contigs with significant overlap were further assembled using Minimus2 with a minimum overlap of 40. Contig postprocessing included splitting misassembled contigs, contig extension and polishing using the strand information of the reads. Single base errors were corrected by aligning the reads back to each contig with BWA to generate a consensus nucleotide sequence. All nine new genomes in this study were annotated using the JGI Annotation pipeline (Grigoriev et al. 2014).

### Phylogenomic analyses

A set of 758 phylogenetic markers, “fungi_odb10”, from the Benchmarking Universal Single-Copy Orthologs (BUSCO) v4.0.5 was employed for the kingdom-wide phylogenomic analyses (Seppey et al. 2019). We used the PHYling pipeline (DOI: 10.5281/zenodo.1257002) to extract best hit copies using hmmsearch v3.3.2 (cutoff=1E^−10^) from the genes predicted in each species against the marker set. A total of 617 (out of 758) well-conserved markers were identified as the best hit from the 181 fungal genomes. A backbone tree including 80 genomes, subsampled based on BUSCO scores and phylogenetic placement (except for the outgroup *Drosophila melanogaster*), recovered 604 orthologs. All orthologs were aligned separately using hmmalign v3.3.2 to the marker profile-HMM and then concatenated into a super-alignment with partitions defined by each marker. The best phylogenomic tree was searched and identified using the super-alignment file and partition scheme as the input with the best-fit model option for maximum likelihood analyses implemented in IQ-TREE v.1.5.5 (Nguyen et al. 2015; Kalyaanamoorthy et al. 2017). Branch supports were evaluated using 1000 ultrafast bootstrap replicates (Hoang et al. 2017).

### Identification of lineage-specific genes and Pfam domains in zygomycete fungi

All orthologous groups of the 80 genomes included in the backbone tree were identified using a comparative genomic pipeline that utilized all-vs-all BLASTp search v2.6.0 (cutoff=1E^−5^) (DOI: 10.5281/zenodo.1447224) (Altschul et al. 1990). Orthagogue v1.0.3 was used to infer putative orthologs and Markov-Clustering Algorithm v14-137 (MCL, inflation value of 1.5) was utilized to generate disjoint clusters (Van Dongen 2000; Ekseth et al. 2014). Shared genome components were counted using a permissive strategy that a gene family shared by at least 10 of the 80 included taxa was retained. Zygomycetes-specific genes are the ones that only exist in zygomycete fungi (Mucoromycota and Zoopagomycota) and are absent in all other lineages. The absence-presence pattern of gene families across the Kingdom Fungi was plotted using the “aheatmap” function in R package “NMF” (Gaujoux and Seoighe 2010). Protein domains coded by the 80 taxa were examined in a similar way. Each Protein Family (Pfam) entry in the Pfam database v31.0 was searched against the predicted proteomes of all included 80 taxa (using the threshold of 1E-3 with >50% overlap percentage). The Pfam domains dominated in either Mucoromycota or Zoopagomycota were inferred by the ratios of their copy numbers in Zoopagomycota and Mucoromycota. The disproportion was visualized by plotting the binary logarithm of the ratio for each Pfam entry so that dominated Pfam domains in each phylum will be isolated on the edge. The figure was plotted using R package “ggplot2” (Wickham 2016). Subphylum-level distribution of each discussed Pfam domain was plotted using the “radarchart” function implemented in R package “fmsb”. All lineage-specific genome content was summarized in Table 2 (with detailed Pfam names listed in Supplementary Table 2). Gene Ontology (GO) terms of Zoopagomycota “unique” genes were inferred and annotated using InterProScan v5.54 and WEGO v2.0 respectively (Jones et al. 2014; Ye et al. 2018).

**Table 2.**
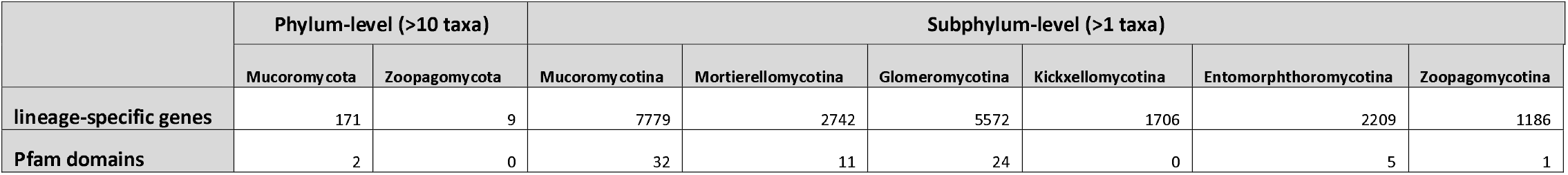
Summary of phylum-level and subphylum-level lineage-specific genes and Pfam domains in zygomycete fungi

### Phylogenetic analysis of the spore coating protein (CotH) in fungi

A total of 846 protein sequences that contain at least one CotH domain were identified in the 80 genomes included in the backbone tree. Absent in all Dikarya species, CotH genes were largely found in zygomycetes (all included six Mortierellomycotina members, 27 Mucoromycotina taxa, and one *Basidiobolus*) and in Neocallimastigomycota (including 2 taxa). Previously classified CotH families 1-5 (CotH 1-5) from *Rhizopus oryzae* were included in our phylogenetic analyses to categorize the newly identified CotH copies. Highly similar CotH sequences (>90%) were removed using CD-HIT v4.6.4 and poor-quality ones were manually excluded from the multiple sequence alignment using MUSCLE v3.8.31 (Edgar 2004; Fu et al. 2012). We employed IQ-TREE v1.5.5 to identify the most appropriate substitutional model and to reconstruct the phylogenetic tree of all fungal CotH copies with ultrafast bootstraps (1000 replicates) (Nguyen et al. 2015; Hoang et al. 2017; Kalyaanamoorthy et al. 2017). The final input includes 754 sequences with 230 distinct patterns for CotH classification. Species-gene tree reconciliation analysis was conducted with Notung v3.0 BETA using the 80-taxa backbone tree as the species tree (Stolzer et al. 2012). The --*phylogenomics* command line option was used to generate a summary report of gain and loss events of CotH families in Kingdom Fungi.

## RESULTS

### Phylogenetic relationships and genome statistics of zygomycete fungi

Collaborative efforts to sequence fungi have generated the 131 zygomycetes genomes presented in this study and the relationships among these species has remained an open research question. Most of the assembled zygomycete genomes were assessed to have BUSCO scores higher than 80% (Fig. 1a, Table 1). The phylogenetic analysis using all available zygomycetes genomes and 50 additional representatives from other fungal clades (Fig. 1a and Supplementary Figure 1) provided an updated tree representing the placement of these fungi in the kingdom. At the phylum level, the reconstructed phylogeny exhibits the same topology as presented in Spatafora et al. (2016). That is, Zoopagomycota forms a sister group to the clade comprising Mucoromycota and Dikarya, and the traditional zygomycete fungi (Mucoromycota and Zoopagomycota) remain paraphyletic. The increased sampling size and new set of protein-coding gene phylogenetic markers provide additional confidence in these arrangements. This is in contrast to a kingdom-wide study that also uses protein-coding genes from BUSCO datasets suggests that zygomycetes could still be monophyletic with a different sampling strategy (Li et al. 2021). It should be noted that the marker sets used in this study (fungi_odb10 with 758 markers) and Li et al. (fungi_odb9 with 290 markers) differ, as well as the strategies to extract the hits—protein searches against genome annotations in this study and BUSCO predicted gene models in Li et al. Regardless of whether zygomycetes are paraphyletic or monophyletic, it is not controversial that Mucoromycota and Zoopagomycota are monophyletic phyla. At the subphylum level, however, the tree topology recovered new phylogenetic relationships with consistency in both the comprehensive tree (Fig. 1a) and the backbone tree (Fig. 2a). For example, Glomeromycotina grouped with Mortierellomycotina instead of being the earliest branch within Mucoromycota (Spatafora et al. 2016). *Basidiobolus* members were not grouped within Entomophthoromycotina, instead, they were found as the earliest diverging lineage in Zoopagomycota (Figs. 1a, 2a, and Supplementary Figure 1). The present subphylum-level classification received full bootstrap supports in the comprehensive tree (Fig. 1a and Supplementary Figure 1). This tree topology is identical in the backbone tree with strong supports, only two nodes within Zoopagomycota clade receiving relatively low values (82/100, Fig. 2a).

**Figure 1:**
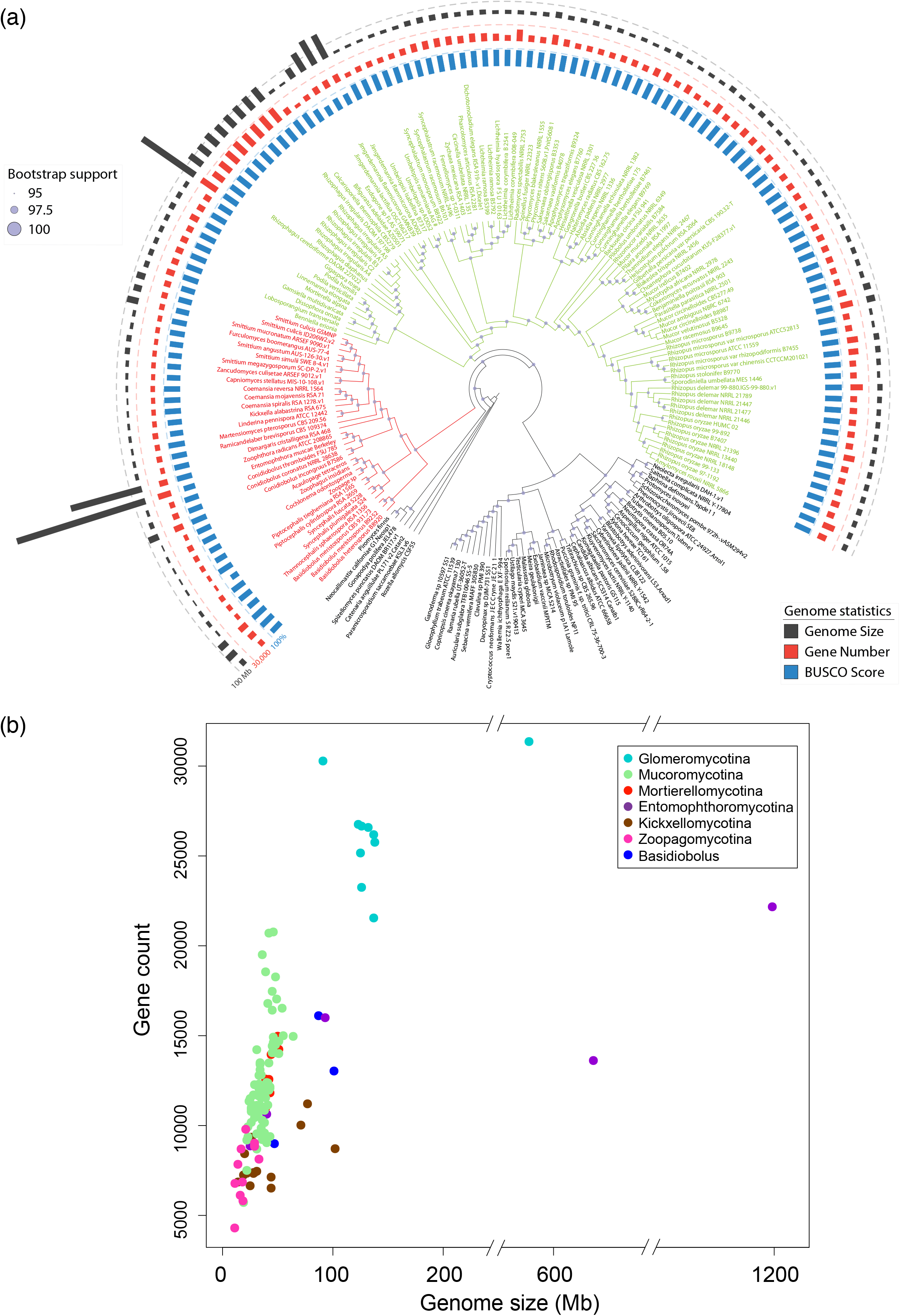
Phylogenetic relationships and genome statistics of zygomycete fungi. (a) The maximum-likelihood tree was inferred from a phylogenomic dataset of 617 protein sequences identified in the included 181 genomes. Branches and tip labels of Mucoromycota and Zoopagomycota were colored in green and red separately. The bootstrap supports are indicated on each node relatively. Tracks from the inside to outside are mapped based on the BUSCO scores, protein-coding gene numbers, and genome size of included zygomycete fungi (detailed bootstrap values and branch lengths are shown in Supplementary Figure 1). (b) The density of protein-coding genes in each genome was plotted using genome sizes on the x-axis against the gene counts on the y-axis. Each dot was colored based on their phylogenetic placement shown in the legend.

**Figure 2:**
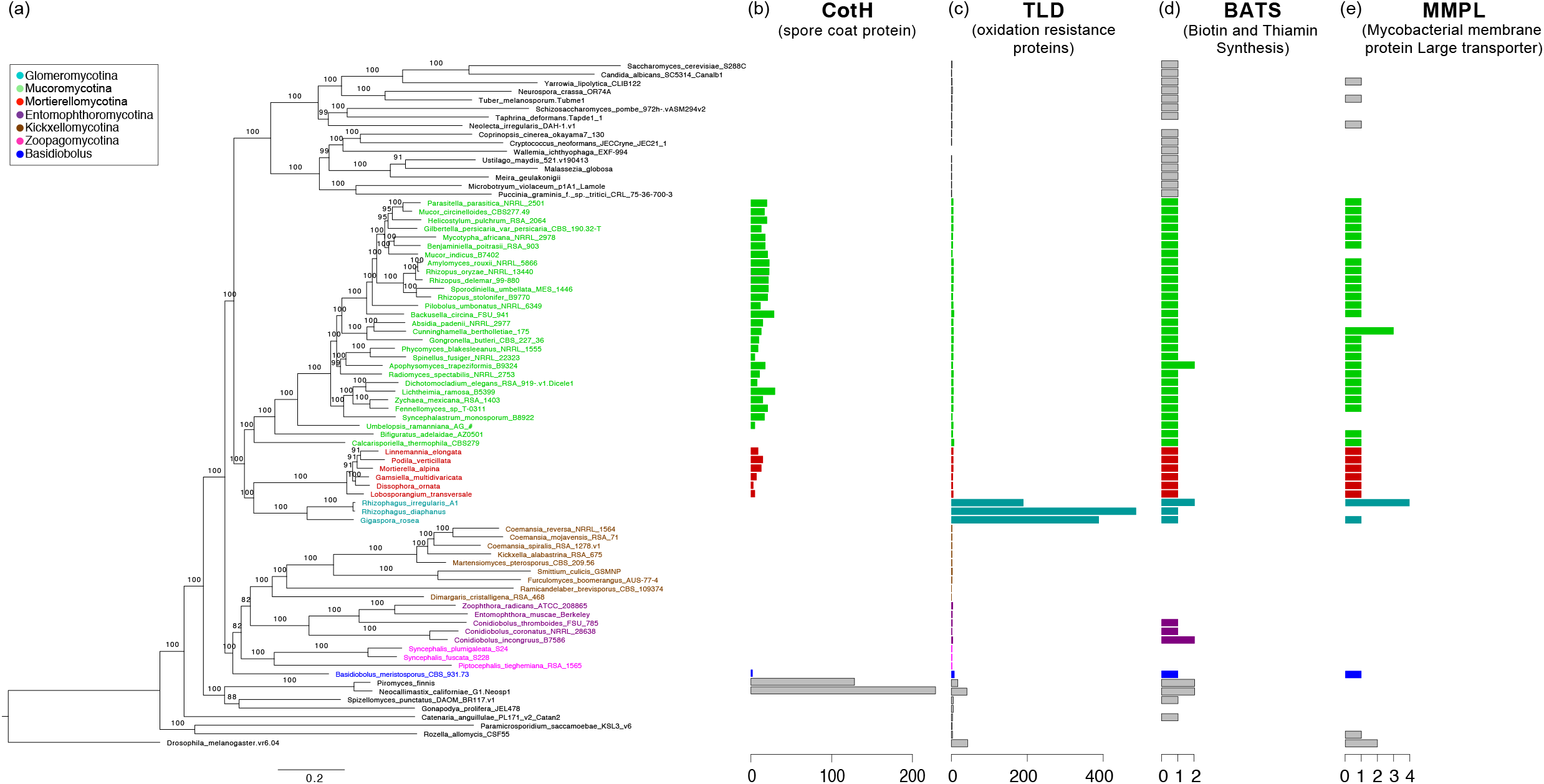
Phylogenetic backbone and highlighted genome content in zygomycete fungi. Zygomycetes genomes that are well assembled (BUSCO score above 80%) and represent unique phylogenetic positions were selected to reconstruct the backbone phylogenomic tree. (a) The backbone phylogenomic tree of zygomycetes includes 80 taxa (rooted with *Drosophila melanogaster*). All bootstrap values (out of 100) were labeled on the branches. (b-e) Protein family domains found with striking patterns in zygomycete fungi are plotted with the copy numbers individually.

Our results suggest that the saprobe, *Calcarisporiella thermophila*, is sister to the rest of the Mucoromycotina. Plant symbionts like *Bifiguratus, Endogone*, and *Jimgerdemannia* form a monophyletic clade which was placed between *C. thermophila* and Mucoromycotina (Fig. 1a). Members of saprobes, pathogens, and mycoparasites were joined in more derived groups of Mucoromycotina.

In the Kickxellomycotina clade, the mycoparasite, *Dimargaris cristalligena*, is sister to the other members. *Ramicandelaber brevisporus* follows and leads to two separate monophyletic clades composed of insect symbionts (*Capniomyces, Furculomyces, Smittium*, and *Zancudomyces*) and soil saprobes (*Coemansia, Kickxella, Linderina, Martnesiomyces*). Both clades (insect symbionts & soil saprobes) are on relatively long branches implying early divergent evolution and underexplored biodiversity (Fig. 2a and Supplementary Figure 1). Insect pathogens were grouped together on a separate lineage, Entomophthoromycotina, forming a sister clade to Kickxellomycotina (Figs. 1a, 2a). The three included *Conidiobolus* species support a paraphyletic genus with the *C. coronatus* monophyletic with *C. incongruus*, while *C. thromboides* was more closely related to *Zoophthora radicans* and *Entomophthora muscae*. Zoopagomycotina is monophyletic and sister to the joined group of Entomophthoromycotina (excluding *Basidiobolus*) and Kickxellomycotina (Figs. 1a, 2a).

The density of genes arranged in the genome of zygomycete fungi exhibited varying patterns among subphyla which was observed in plots of gene counts against genome sizes (Fig. 1b). Most zygomycete fungi have genome sizes ranging from 20 Mb to 100 Mb and gene counts range from 5k to 20k. The Mucoromycotina fungi have relatively similar genome sizes, but gene counts vary from 6k to 21k. The soil saprobes in Kickxellomycotina and small animal associates in Zoopagomycotina have small genome sizes (10-20 Mb) and gene counts (4-8k). On the other hand, Glomeromycotina fungi tend to have large genome sizes (>100 Mb) with the most abundant gene numbers (20-30k) in all zygomycete fungi, which are among the largest fungal genome sizes sequenced to date. As an extreme case, the genome sizes of Entomophthoromycotina members exhibit the widest range and can be as large as 1.2 Gb according to the existing genome assemblies (Stajich et al, in revision), however, their gene counts (9-23k) are more modest.

### Orthologous gene families and Pfam domains in zygomycete fungi

The 80 species used for the backbone tree were examined for orthologous gene families across the Kingdom Fungi. We identified 8,208 orthologous families which had genes from at least 11 of the 80 genomes. These gene families were subjected to more focused analyses to examine the presence/absence pattern of genome contents across the Kingdom Fungi, with a special attention on the divergent evolution between Mucoromycota and Zoopagomycota (Fig. 3). The Mucoromycota members harbor 171 phylum-specific gene families that are present in at least two of the three Mucoromycota subphyla and absent in all other fungal lineages, while Zoopagomycota only have nine such gene families (Table 2). At the subphylum level there were considerably more lineage-specific gene families, ranging from 1,186 (in Zoopagomycotina) to 7,779 (in Mucoromycotina) (Table 2).

**Figure 3:**
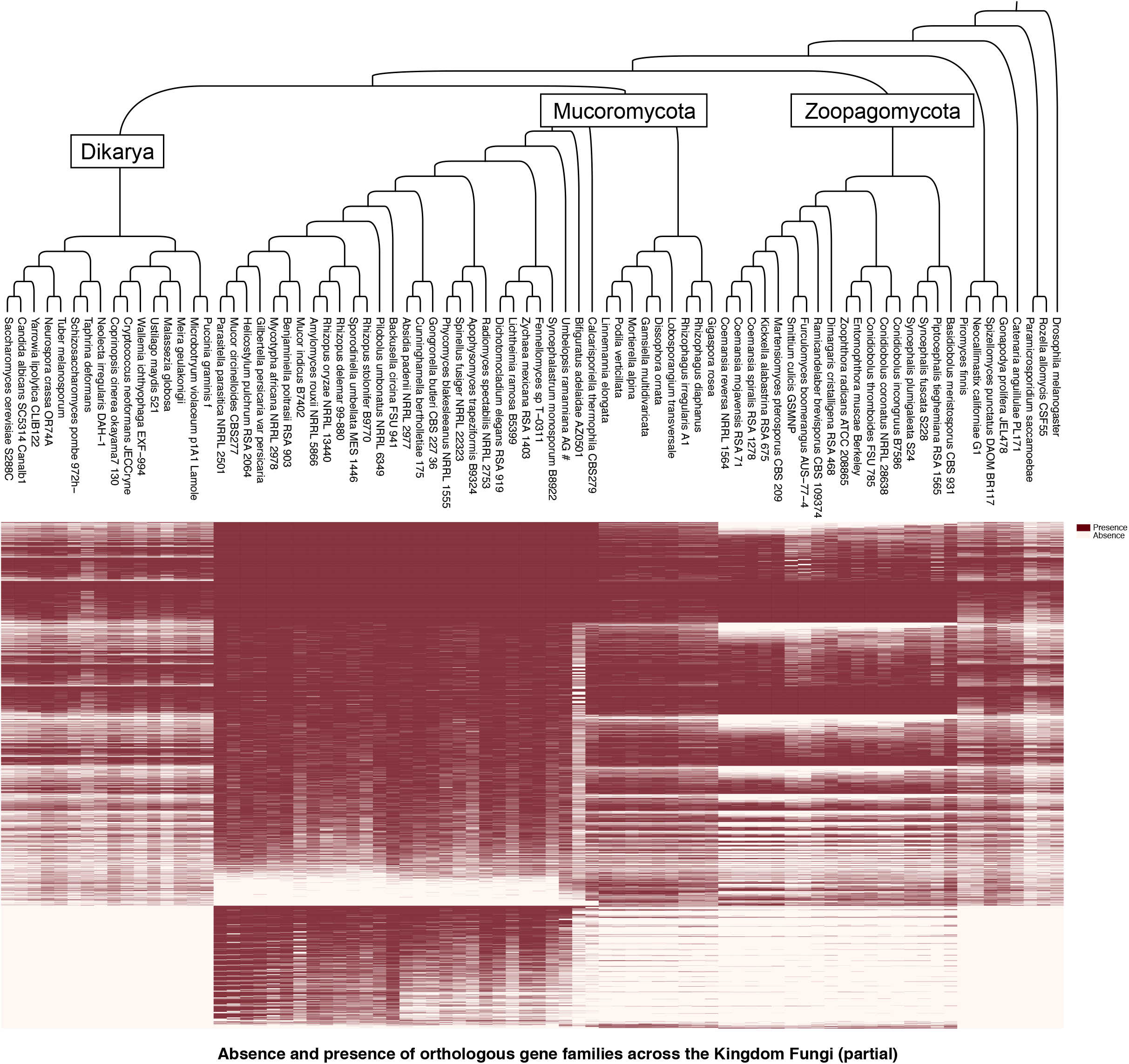
Absence and presence of orthologous gene families across the Kingdom Fungi. Orthologous gene families were examined in the genomes included in the backbone tree. The 8,208 gene families were found present in at least 10 of the 80 taxa and thus included to examine the absence/presence pattern of genome content among different fungal lineages (a complete map showing the unfiltered 62,689 gene families was included in Supplementary Figure 2).

We used protein domains cataloged in the Pfam database as an additional means to catalog unique and shared content. A total of 7,616 Pfam models had at least one similar sequence in the examined 80 genomes. Mucoromycota members possess two unique Pfam domains, with the CheR (PF01739) found in all three subphyla and the C9orf72-like (PF15019) in Mucoromycotina and Mortierellomycotina, while no phylum-specific Pfam domains were identified in the Zoopagomycota. At the subphylum level, a range of unique Pfam domains were observed, with 11 to 32 in the three subphyla of Mucoromycota and 0-5 in the ones in Zoopagomycota (Table 2 and Supplementary Table 2). Interestingly, the CotH domain (PF08757), a potential invasin factor of Mucormycosis, was found in Mortierellomycotina, *Basidiobolus*, and Neocallimastigomycota genomes (Fig. 2b), but had previously only been described in the Mucoromycotina (Chibucos et al. 2016). In addition, the oxidation resistance protein domain (TLD, PF07534) has greatly expanded in copy number in the Glomeromycotina with up to 400 copies (Fig. 2c). Kickxellomycotina and Zoopagomycotina members lacked Biotin and Thiamin synthesis associated domain (BATS, PF06968) and mycobacterial membrane protein large transporter domain (MMPL, PF03176) (Fig. 2d and 2e). Interestingly, *Basidiobolus meristosporus* is the only Zoopagomycota member that maintains at least one copy of every examined domain (Fig. 2b-e), including CotH and MMPL that are absent in all other Zoopagomycota members.

To identify all Pfam domains that are consistent with divergent evolution between Mucoromycota and Zoopagomycota, we calculated the relative abundance of each Pfam domain in their genomes. In total, 285 Pfam domains were present at least four-fold differences (i.e., absolute value of the binary logarithm >2) between the two phyla with 243 of them featured in Mucoromycota while 42 in Zoopagomycota (Fig. 4 and Supplementary Table 3). Without consideration of non-zygomycetes lineages, we found 70 Pfam domains in Mucoromycota that are completely missing in Zoopagomycota, whereas no such Pfam domains can be identified in Zoopagomycota. Zoopagomycota is a historically understudied fungal clade with few representative genomes until our recent studies. As a result, the lack of Zoopagomycota specific Pfam domains may be an artifact of insufficient sampling before domain curation in Pfam. To overcome this possibility, we examined the orthologous gene family dataset to calculate the relative abundance of gene families to test for differences between the two phyla. This revealed 22 gene families in Zoopagomycota that were absent in all Mucoromycota members (Supplementary Figure 3). Gene Ontology analysis shows that more than 50% of these genes are involved in binding, catalytic activity, cellular process, and metabolic process (Supplementary Figure 4). Finer scales of examination suggest they are closely related to nitrogen compound, organic substance, and primary metabolic process (Supplementary Figure 5). This provides a compact set of genes and processes that are distinct to the Zoopagomycota which could contribute to their unique lifestyles, reproductive biology, and ecological specializations.

**Figure 4:**
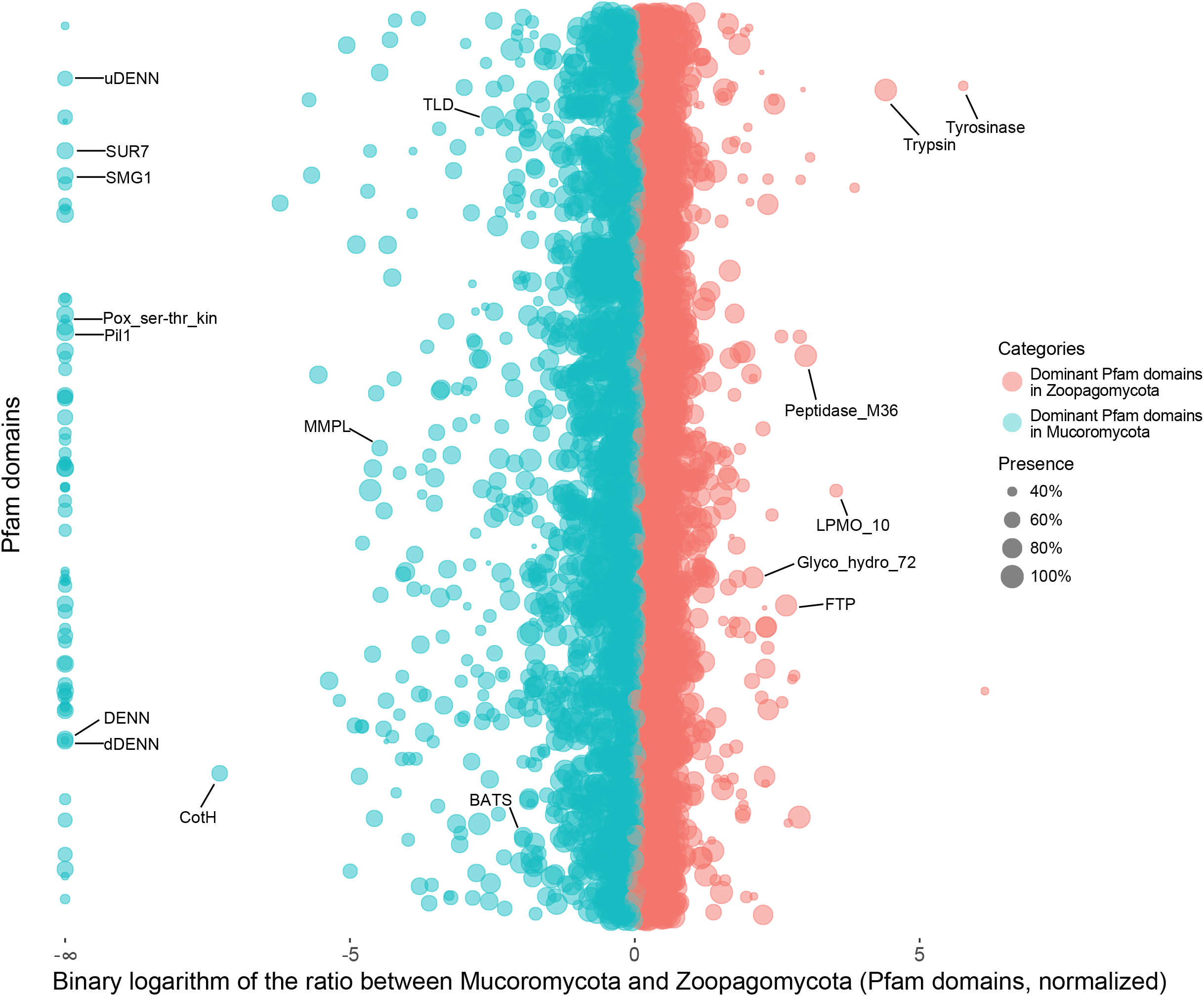
Protein family (Pfam) domains with differentiated enrichment in Mucoromycota or Zoopagomycota. Each dot represents a Pfam domain found in zygomycete fungi. The x-axis is the binary logarithm of the Pfam copy ratios between Zoopagomycota and Mucoromycota, and the y-axis is used to rank the Pfam domains in alphabetical order. The Pfam domains enriched in Mucoromycota are shown on the left side in cyan color, and the Zoopagomycota enriched ones are on the right side in red color. The bubbles (Pfam domains) with bigger sizes are shared by more zygomycetes members. The Pfam domains aligned on the left edge are domains only found in Mucoromycota and absent in Zoopagomycota. The domains discussed in the text were labeled with the Pfam name. A detailed chart including the names and ratios of all Pfam domains are also provided (Supplementary Table 3).

We found that many phylum-level distinct Pfam domains were favored unevenly in each subphylum group (Fig. 5). For example, both Pil1 (PF13805) and SUR7 (PF06687) domains are eisosome components and are involved in the process of endocytosis. They are missing entirely from the Zoopagomycota but are encoded in the genomes of all (Pil1) or a majority (SUR7, except for *Mortierella multidivaricata* and *Gigaspora rosea*) of Mucoromycota members (Figs. 4, 5a, & 5b). Interestingly, the Pil1 domain was enriched in copy number in the Mortierellomycotina (Fig. 5a), and SUR7 domain has the largest copy number in Mucoromycotina (Fig. 5b). The SMG1 domain (PF15785), a phosphatidylinositol kinase-related protein kinase, is a key regulator of growth. The Mucoromycota members maintain a single-copy SMG1 domain (except for *Cunninghamella bertholletiae* with 3 copies, and none in *Mucor circinelloides, Phycomyces blakesleeanus*, and *Syncephalastrum monosporum*), which is absent in Zoopagomycota species (Fig. 5c). There are 67 additional Pfam domains including DENN (PF02141), uDENN (PF03456), dDENN (PF03455), Pox_ser-thr_kin (PF05445) (Supplementary Table 3) with a similar presence/absence pattern and may be important components to better understand and characterize the Mucoromycota fungi.

**Figure 5:**
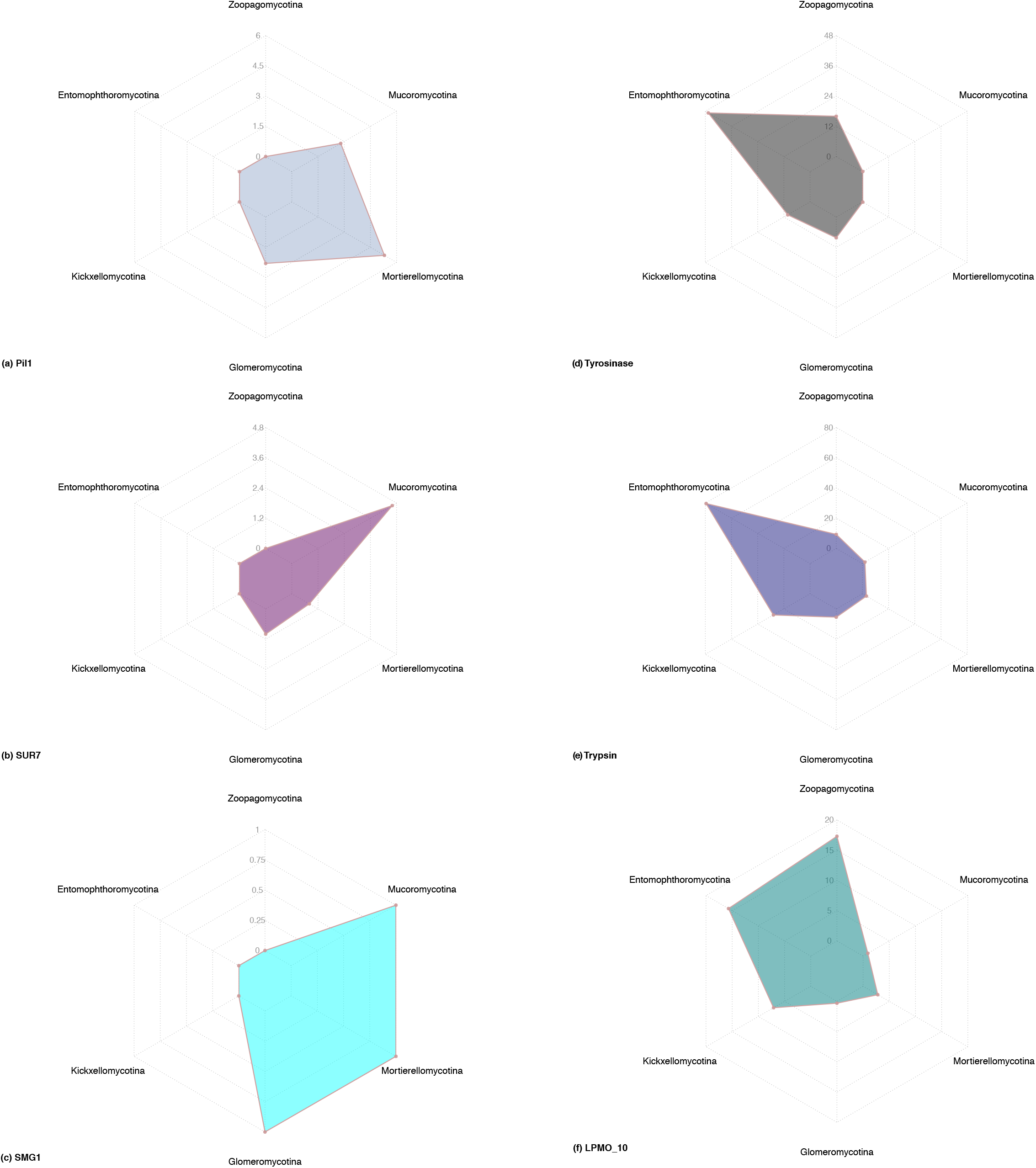
Subphylum-level distribution of six Pfam domains that may contribute to the divergent evolution of zygomycete fungi. The scales on each axis of the radar plots indicate the average copy number of the domain in each subphylum. (a-c) Pfam domains shared in all Mucoromycota subphyla and absent in the entire Zoopagomycota. (d-f) Distinct Pfam domains in Zoopagomycota subphyla and largely missing in Mucoromycota.

In contrast, while there are no Zoopagomycota-specific Pfam domains, there are some domains that exhibit copy number variance at the subphylum level. For example, the Tyrosinase domain (PF00264) is an important enzyme that controls the production of melanin and parasite encapsulation, especially in insects. It is also suggested that Tyrosinase is involved in the host-microbe defensive mechanism. The Tyrosinase domains are found on average with 48 copies in the Entomophthoromycotina members but absent in nearly all Mucoromycotina (except for *Calcarisporiella thermophila* with 7 copies) and Mortierellomycotina (except for *Mortierella verticillata* with 1 copy) (Fig. 5d). Similarly, Trypsin domain (PF00089), serine protease found in the digestive system of many vertebrates, was also enriched in copy number in the Entomophthoromycotina with 80 copies on average (Fig. 5e). The domain LPMO_10 (PF03067) is found in lytic polysaccharide monooxygenases which can cleave glycosidic bonds in chitin and cellulose and is significantly enriched in Zoopagomycota (Fig. 5f). All three examples (Trypsin, Tyrosinase, and LPMO_10) are related to animal-fungus interactions in the degradation of protein, chitin, and cellulose.

### Discovery of CotH in early-diverging fungi

The CotH domain as characterized in Mucorales fungi has positive correlations with the clinical pathogenesis of Mucormycosis (Chibucos et al. 2016). In our kingdom-wide study, we found additional copies of the CotH domain in a broader collection of fungi. Other than in Mucorales fungi, CotH was also found in *Basidiobolus*, Mortierellomycotina, and Neocallimastigomycota. The presence of this domain could indicate the potential of these fungi to support pathogenic interaction with animal hosts (Fig. 2b). A total of 846 CotH copies were identified in 34 zygomycetes genomes and two Neocallimastigomycota representatives (contributing 348 of the copies). Five CotH families (CotH 1-5) that were previously classified in *Rhizopus oryzae* were included in our phylogenetic analysis and helped us categorize the newly identified CotH copies (Fig. 6a). Zygomycetes CotH copies formed four distinct clades. ZyGo-A clade includes CotH families 1-3 that maintain true invasin motifs and can only be coded by Mucoromycotina and Mortierellomycotina members. ZyGo-B clade includes CotH families 4-5 and copies from Mucoromycotina. ZyGo-C clade is loosely joined with ZyGo-B (bootstrap support 34/100) and includes copies from Mortierellomycotina, and *Basidiobolus*. ZyGo-D is the largest clade among the four but only includes copies from Mucoromycotina. Both ZyGo-C and ZyGo-D clades represent new families of CotH. Interestingly, the distantly related anaerobic gut fungi (AGF, Neocallimastigomycota) can code CotH as well and form several distinct clades. In total, 531 duplications and 795 losses were identified along the evolution of CotH families in Kingdom Fungi. Nine nodes were associated with more than 10 duplication events (Fig. 6b). The absence of CotH in the most recent common ancestor of fungi was also inferred by Notung reconciliation analysis.

**Figure 6:**
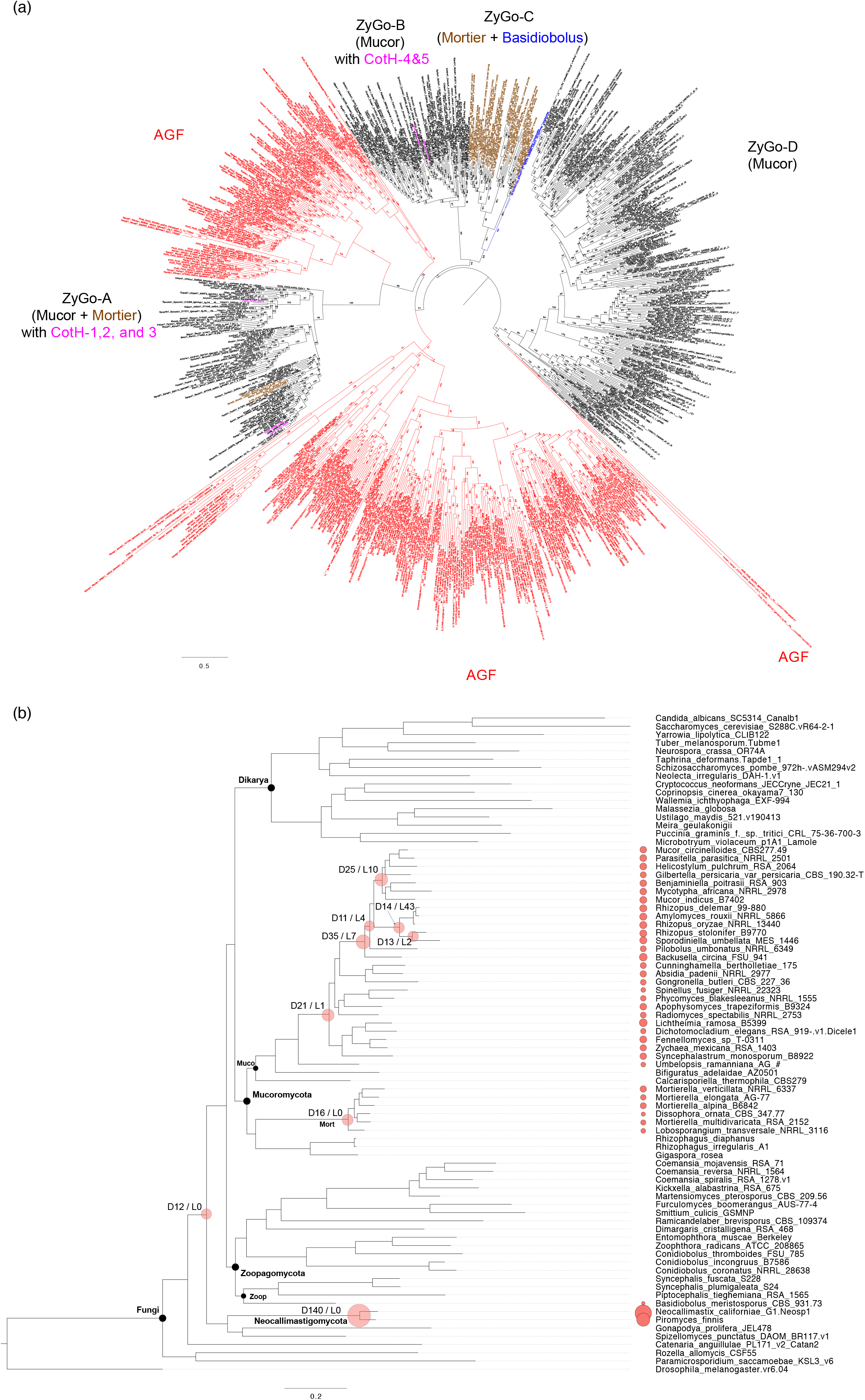
Phylogenetic analysis and evolution of CotH in Kingdom Fungi. (a) The 754 fungal CotH copies were identified from Mortierellomycotina (brown), Mucoromycotina (black), *Basidiobolus* (blue), and Neocallimastigomycota (red). The CotH phylogenetic tree was midpoint rooted and reconstructed using the maximum likelihood method with bootstrap supports (out of 100) labeled on each branch. The analysis included previously classified CotH families 1-5 (pink) to help categorize newly identified fungal CotH. (b) Reconstruction of CotH evolution in Kingdom Fungi with Notung. CotH copies identified in each genome were plotted at tree tips with proportional sizes. Nodes with more than 10 duplication events were highlighted with red bubbles and labelled with duplication (“D”) and loss (“L”) events. Node abbreviation: Muco, Mucoromycotina; Mort, Mortierellomycotina; Zoop, Zoopagomycotina.

## DISCUSSION

### Genome evolution of zygomycete fungi

Zygomycetes are important members of early-diverging fungi and studying their evolutionary history can help us better understand the terrestrialization of eukaryotes. Zygomycetes are ubiquitous and can live as arbuscular mycorrhizae, ectomycorrhizae, saprobes, or symbionts of various organisms, including animals, bacteria, plants, and fungi. During the evolutionary adaptation and diversification of zygomycetes, many associated organisms (hosts, symbionts, etc.) may have mutually shaped the structure and content of their genomes. Mucoromycotina members have served as exemplars to investigate various evolutionary events at the genome-scale. For example, whole-genome duplications have been identified repeatedly in Mucoromycotina (Ma et al. 2009; Corrochano et al. 2016), which contributed to the large expansion of gene counts (5-20k) among some Mucoromycotina members (Fig. 1b). Phylogenomic analyses suggest that an early split of Mucoromycotina involved the evolution of thermophily (i.e., *Calcarisporiella thermophila*) (Figs. 1a, 2a), which is followed by various lineages containing members of ectomycorrhizae, mycoparasites, plant and animal pathogens. In addition, transposable elements (TEs) have been suggested to be rich in some Mucoromycotina taxa, including *Rhizopus oryzae* (Ma et al. 2009) and *Endogone* sp. (Chang et al. 2019). The high proportion of TEs were also evident in other lineages of zygomycete fungi, like *Gigaspora* members (Morin et al. 2019) and *Basidiobolus meristosporus* (Muszewska et al. 2017). It has been suggested that TEs may have played a role in shaping transcriptional profiles, helped fungi adapt to different ecological niches, and contributed to the current fungal biodiversity (Castanera et al. 2016; Muszewska et al. 2017). It is still unclear what roles TE might have played in the evolution of Entomophthoromycotina members that exhibit the widest span of genome sizes (25-1200 Mb) in Kingdom Fungi and what resulted in the gigantic size of *Entomophthora muscae* and *Massospora cicadina*. More samples from this and related lineages may help us reconstruct the evolutionary history for the observed genome size modification in zygomycete fungi.

### *Phylogenomics of zygomycetes and* Basidiobolus

Zygomycete fungi hold important phylogenetic placement on the fungal tree of life. The former taxonomic unit, Zygomycota, has been recognized paraphyletic and thus been abandoned and replaced by Mucoromycota and Zoopagomycota to accommodate the six major lineages—Glomeromycotina, Mortierellomycotina, Mucoromycotina, Entomophthoromycotina, Kickxellomycotina, and Zoopagomycotina (James, Kauff, et al. 2006; White et al. 2006; Hibbett et al. 2007; Spatafora et al. 2016). Since the loss of flagella, the first evolutionary split of terrestrial fungi leads to Zoopagomycota and the clade of Mucoromycota and Dikarya (Chang et al. 2021). Mucoromycota is the sister clade of the subkingdom Dikarya clades (Ascomycota and Basidiomycota) (Figs. 1a & 2a), and analysis of zygomycete fungi is therefore essential for us to accurately reconstruct the evolutionary events that led to major lineages of terrestrial fungi. The arbuscular mycorrhizal fungi of Glomeromycotina with their distinct ecology formed a monophyletic clade with the soil saprobes and root endophytes of Mortierellomycotina (Figs. 1a & 2a). Mucoromycota members are mostly associated with plants or more commonly as decomposers of plant carbohydrates. Zoopagomycota members are mostly animal associated (either as commensals or pathogens) or mycoparasites. The Entomophthoromycotina clade presents several interesting patterns. For example, our phylogenomic results confirm the non-monophyly of *Conidiobolus* and encourage further work to reclassify this genus. One recent taxonomic effort based on a four-gene phylogeny proposed three new genera (*Capillidium, Microconidiobolus*, and *Neoconidiobolus*) to help delimitate *Conidiobolus* where *C. thromboides* was suggested in the *Neoconidiobolus* clade (Nie et al. 2020). In addition, our results suggest *Basidiobolus*, a traditional member of Entomophthoromycotina, as the earliest diverging lineage within Zoopagomycota (Figs. 1a, 2a & Supplementary Figure 1).

*Basidiobolus* has been characterized as a “rogue” taxon and is often found with conflicting phylogenetic placements. Using nuclear rRNA genes (18S+28S+5.8S genes), *Basidiobolus, Olpidium brassicae* (a plant pathogen), and *Schizangiella serpentis* (a snake pathogen) were grouped together and placed at the earliest diverging branch within Zoopagomycota (White et al. 2006). In a separate study using four genes (nuclear 18S and 28S rDNA, mitochondrial 16S, and RPB2), *Basidiobolus* was suggested as the earliest diverging member of Entomophthoromycotina (Gryganskyi et al. 2012). The recent genome-scale study based on 192 conserved orthologous proteins favored the *Basidiobolus* placement in Entomophthoromycotina as well (89/100 bootstrap support) (Spatafora et al. 2016). Interestingly, a recent genome-scale phylogenetic study examining the entire Kingdom Fungi found that *Basidiobolus* formed a sister clade to Mucoromycota instead of joining Zoopagomycota (Li et al. 2021). In the present study, we included the largest collection of zygomycete genomes to date and employed the newly released 758 “fungi_odblO” markers. The results suggested that *Basidiobolus* formed their own group within Zoopagomycota as the earliest diverging lineage (with 100/100 bootstrap, Supplementary Figure 1). The complexity of *Basidiobolus* is also evidenced by their enriched secondary metabolite genes, regionally duplicated genomes, and broad range of potential hosts including insects, amphibians, reptiles, and human beings (Henk and Fisher 2012; Tabima et al. 2020). This may explain the sources of phylogenetic conundrums that we have encountered in the last decades using different molecular markers. The natural history of *Basidiobolus* may not be easily resolved until an appropriate approach can be carried out to parse their complex genome composed of redundant genes from various sources, such as large-scale gene duplications or horizontal gene transfers. In addition, the kingdom-wide comparison helped discover many unique genome components in *Basidiobolus*, including the ones featured in Mucoromycota clades (e.g., CotH and MMPL), which will be discussed in the following sections.

### Divergent evolution of zygomycete fungi

We were able to identify different sets of gene content and Pfam domains favored by each of the zygomycete phyla, which well correspond to their disparate lifestyles (Figs. 3 & 4). As suggested by the presence of both Pil1 and SUR7 domains, eisosome-mediated endocytosis and related active transportation are important facilitators to saprotrophic Mucoromycota fungi (Walther et al. 2006). Among the 70 Mucoromycota-featured domains (Fig. 4 and Supplementary Table 3), DENN, uDENN, and dDENN also serve as regulators during eukaryotic membrane trafficking events (Zhang et al. 2012). This implies that Mucoromycota fungi are able to transport particles via membrane trafficking domains, while Zoopagomycota fungi, as animal-associated microbes, may use different mechanisms. Noteworthy, the Pox_ser-thr_kin, a poxvirus serine/threonine protein kinase, specifically identified in Mucoromycota genomes (Fig. 4 and Supplementary Table 3) indicates that large DNA viruses are embedded in Mucoromycota genomes (Jacob et al. 2011). Mycoviruses have been extensively studied in Dikarya fungi, especially for plant pathogens (Ghabrial et al. 2015; Marzano et al. 2016). The existence of mycoviruses among early-diverging fungi have not been examined until recently, which led to the discovery of a fungal–bacterial-viral system in the plant pathogenic *Rhizopus microsporus* (Espino-Vázquez et al. 2020) and RNA mycoviruses in roughly one fifth laboratory cultures of early diverging fungal lineages (Myers et al. 2020). Our preliminary analyses suggest that Mucoromycota members contain genomic hallmarks that interact with both bacteria (MMPL domain, Fig. 2e) and viruses (Pox_ser-thr_kin domain, Supplementary Table 3). The “mycobacterial membrane protein large transporter” domain is well represented in all three subphyla of Mucoromycota as well as *Basidiobolus* (Fig. 2e) with the fact that fungal-bacterial interactions have been well documented in these lineages (Uehling et al. 2017; Desirò et al. 2018; Chang et al. 2019; Bonfante and Venice 2020; Tabima et al. 2020). Although the TLD domain universally presents in almost all fungal lineages (except *Wallemia ichthyophaga*), the exceptionally large number of TLD domains identified in Glomeromycotina members are unusual (Fig. 2c). It implies that TLD and related oxidation resistance proteins serve important functions to protect these arbuscular mycorrhizal fungi from reactive oxygen species (Blaise et al. 2012).

Zoopagomycota, on the other hand, do not have exclusive Pfam domains, even though many domains are highly enriched suggesting important functions. Tyrosinase is a good example. Tyrosinase domains have well-recognized functions to synthesize melanin via the amino acid L-tyrosine in melanosomes. Melanin is important to protect organisms from diverse biotic and abiotic factors, including helping microbes counteract the attacks from host immune systems, specifically by neutralizing reactive oxygen species or other harmful molecules (Cordero and Casadevall 2020). As such, it is not surprising to find that Zoopagomycota fungi, especially the insect-associated ones, maintain a large number of melanin synthetic enzymes presumably helping them evade host immune responses. Trypsin is another Pfam domain featured in Zoopagomycota (Fig. 4). Trypsin domains catalyze the hydrolysis of peptide bonds and break proteins into smaller pieces, which is extremely active in digestive systems. We discovered up to 59 copies (in *Smittium culicis*) of Trypsin domain in the insect gut-dwelling fungi (Harpellales, Kickxellomycotina). Interestingly, insect pathogenic species in Entomophthoromycotina were found heavily relying on hydrolases with 204 copies of Trypsin domains in *Zoophthora radicans* alone (43-138 copies in other Entomophthoromycotina members), while other zygomycete lineages maintain 0-18 copies variously (Fig. 5e). Trypsin and Trypsin-like proteases have been studied in insects and entomopathogenic fungi for decades (Paterson et al. 1993; Dubovenko et al. 2010; Lazarević and Janković-Tomanić 2015). Results suggest that the Trypsin and Trypsin-like proteins are important for nutritional uptake and pathogenic processes of insect-associated fungi, which was also suggested with the potential to help develop new agents to control pest insects (Borges-Veloso et al. 2015; Lazarević and Janković-Tomanić 2015). The abundance of Trypsin domains identified in Zoopagomycota suggests that the expansion of Trypsin across fungal tree of life have occurred more than once (e.g., Ascomycota and Zoopagomycota) (Dubovenko et al. 2010). In addition, the emergence and detailed evolutionary patterns of Trypsin and Trypsin-like proteins in Ascomycota, Zoopagomycota, and insects deserve further examination. Many polysaccharides and protein degrading enzymes were also found expanded in Zoopagomycota, such as LPMO_10, Glyco_hydro_72 (PF03198), and Peptidase_M36 (PF02128) (Fig. 4), suggesting their important functions during the interactions of Zoopagomycota fungi with small animals or other fungi. The fungalysin metallopeptidase (Peptidase_M36) and the associated fungalysin/thermolysin propeptide motif (FTP, PF07504) were both found expanded in the obligate mycoparasite *Syncephalis* (Lazarus et al. 2017). Both domains may help mycoparasites inhibit peptidases produced by the hosts, but their exact function has not been clearly known (Markaryan et al. 1996; Finn et al. 2016). Interestingly, the BATS domain involved in the biotin and thiamin synthesis is found absent in Kickxellomycotina and Zoopagomycotina members (Fig. 2d). Both subphyla are short for available cultures, which is especially the case for the animal associated species. The inability to synthesize biotin and thiamin may be one of the culprits for the unsuccessful culture establishment in the lab. Supplementary biotin and thiamin could be suggested for future efforts on development of new cultures in these fungal lineages.

### Human infectious diseases caused by zygomycete fungi

Mucormycosis is a deadly human-infectious disease usually caused by *Rhizopus, Mucor*, and *Lichtheimia*. The current COVID-19 pandemic has triggered multiple cases of Mucormycosis in susceptible patients (Garg et al. 2021; Revannavar et al. 2021). The CotH was originally identified in bacteria as a spore-coat protein. It was later found in Mucorales fungi and identified as a potential invasin factor of the human-infectious Mucormycosis. The CotH was suggested to be directly involved in interactions between Mucorales pathogens and human endothelial cells (Chibucos et al. 2016). Our comparative genomic analyses provided a broader survey of CotH leading to discoveries of novel CotH families in Mucoromycotina strains and unexpected fungal lineages (*Basidiobolus*, Mortierellomycotina, and Neocallimastigomycota). CotH was maintained by almost every member of Mucoromycotina except the early-diverging taxa—*Calcarisporiella thermophila* and *Bifiguratus adelaidae*. Unexpectedly, all members of Mortierellomycotina were also able to code CotH domains with the same or highly similar pathogenic motif “MGQTNDGAYRDPTDNN”, which was proposed as a key factor for Mucormycosis. This implies that the included Mortierellomycotina taxa (*Dissophora ornate, Lobosporangium transversle*, and *Mortierella* species) may be facultative pathogens or have the potential to cause Mucormycosis or related human-infectious diseases if treated without caution. The results are informative to guide clinical practice as Mucormycosis may arise from many previously less documented situations, including the injuries during the natural disasters, unconscious contact, and triggered by other diseases like Novel Coronavirus Pneumonia (caused by COVID-19) (Neblett Fanfair et al. 2012; Revannavar et al. 2021). *Basidiobolus* is the only Zoopagomycota member that encodes CotH, albeit the copy number is low. On the other hand, Neocallimastigomycota members produce surprisingly high numbers of CotH domains with the largest duplication event (Fig. 6b). It is not clear why anaerobic gut fungi maintain so many CotH copies since they serve as primary plant polysaccharide degraders and do not pose any identifiable harm to their mammal hosts. Phylogenetic analyses suggest that CotH domains in fungi can be classified into at least seven major groups (ZyGo-A, B, C, D, and three AGF groups; Fig. 6a). The ZyGo-A is the only clade containing all known Mucormycosis invasin factors (i.e., CotH 2 and CotH 3) where Mortierellomycotina members are tightly clustered (Chibucos et al. 2016). The members in ZyGo-A, Mucoromycotina and Mortierellomycotina, should both have the potential to cause Mucormycosis.

There are additional emerging pathogens in Zoopagomycota. For example, members of the entomophthoralean fungi can cause infection in both insects and mammals, not only in immunocomprised patients, but also reported from immunocompetent individuals due to insect bites or other undetermined environmental contacts, especially in tropical and subtropical regions (Vilela and Mendoza 2018). *Basidiobolus* and *Conidiobolus* are two additional agents of human skin, subcutaneous, and gastrointestinal infections (Khan et al. 2001; Shaikh et al. 2016). *Basidiobolus* can be isolated from various types of environments, including soils or leaf litters, dung of frogs or lizards, and various insects (e.g., mosquitoes, mites, springtails) (Lyon et al. 2001; Garros et al. 2008; Manning and Callaghan 2008; Werner et al. 2012). Recently, people also found that *Basidiobolus* can infect human eyes (Tananuvat et al. 2018; Vilela and Mendoza 2018). The two CotH copies identified in *Basidiobolus* genomes may be involved in the pathogenic processes. *Conidiobolus*, however, do not maintain CotH copies, suggesting that *Conidiobolus* may take different strategies to infect mammalian hosts. Our comparative genomic analyses provided a broader view regarding the molecular mechanism of human-infectious zygomycete fungi. As the quick accumulation of genomic resources for this fungal lineage, a detailed natural history and complete pathogenic pathways should be revealed in the near future.

Our combination of phylogenomic and comparative genomic study of zygomycete fungi provided a fresh view of the phylogenetic relationships within the group. Multiple lineage-specific genome contents were revealed to help understand their cryptic ecology and relationships with other organisms in the environment. The unexpected findings of CotH in Mortierellomycotina, *Basidiobolus*, and Neocallimastigomycota proved the effectiveness of comparative genomics to predict biology of understudied organisms and the importance of fungal studies in the era of global climate change. The presented results may be applied to prevent further damage caused by the human-infectious Mucormycosis.

## DATA AVAILABILITY

Assembled genomes and annotation files are available at JGI MycoCosm website. All genome accession numbers are listed in Table 1.

## ACKNOWLEDGEMENTS

This material is based upon work supported by the National Science Foundation (DEB-1441604 to JWS and DEB-1441715 to JES), The authors thank Drs. M. Catherine Aime, William J. Davies, Gunther Doehlemann, Toni Gabaldón, Timothy Y. James for permission to use genomes ahead of publication. The work (proposals 10.46936/10.25585/60001019 and 10.46936/10.25585/60001062) conducted by the U.S. Department of Energy Joint Genome Institute (https://ror.org/04xm1d337), a DOE Office of Science User Facility, is supported by the Office of Science of the U.S. Department of Energy operated under Contract No. DE-AC02-05CH11231. JES is a paid consultant for Zymergen and Sincarne.

