## Supplementary Files for "Divergent evolution of early terrestrial fungi reveals the evolution of Mucormycosis pathogenicity factors"

**This supplementary document includes:**
Supplementary Figures 1-5
Supplementary Tables 1-3

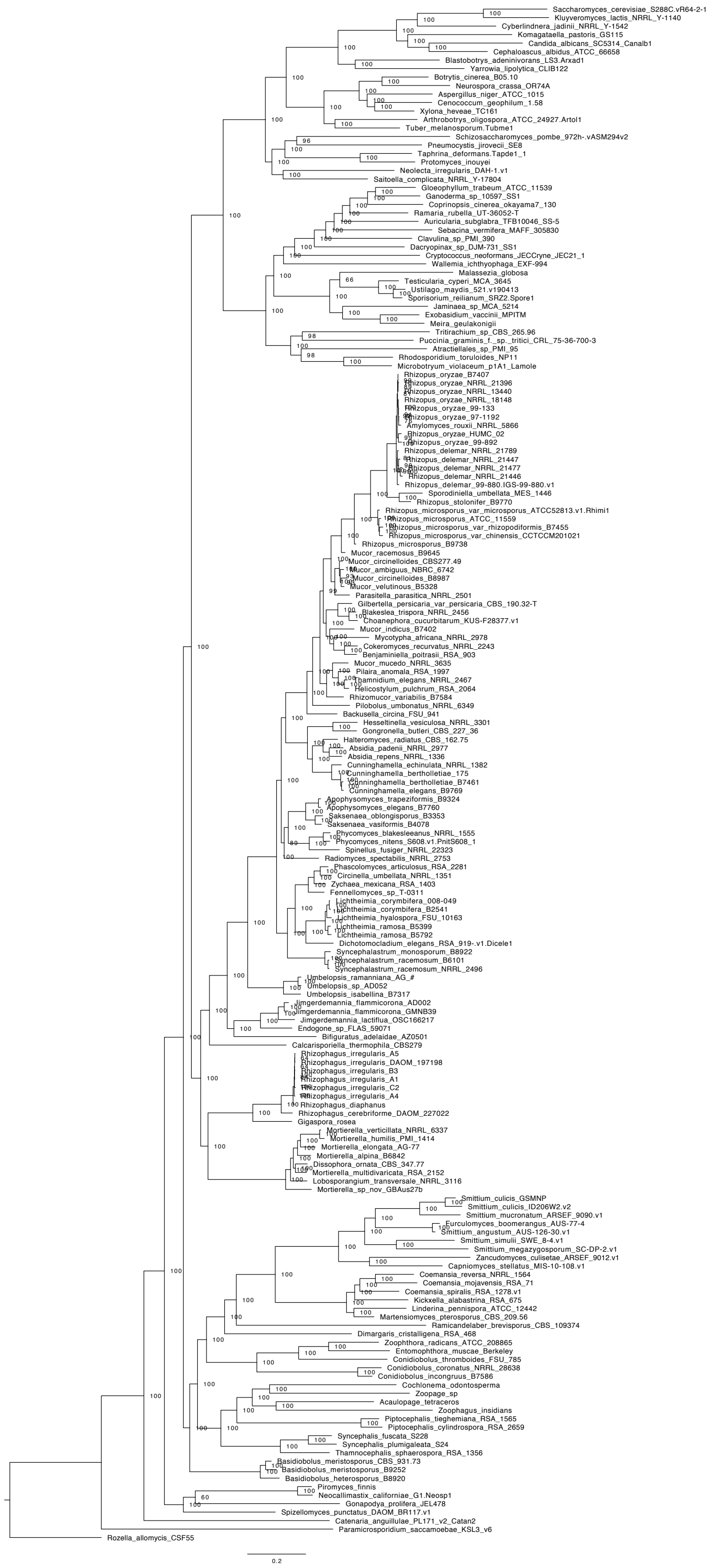

Supp. Fig. 1 The maximum-likelihood phylogenetic tree of the Kingdom Fungi using 617 markers sampled from 181 taxa. Ultrafast bootstrap values (out of 100) are indicated on each node.

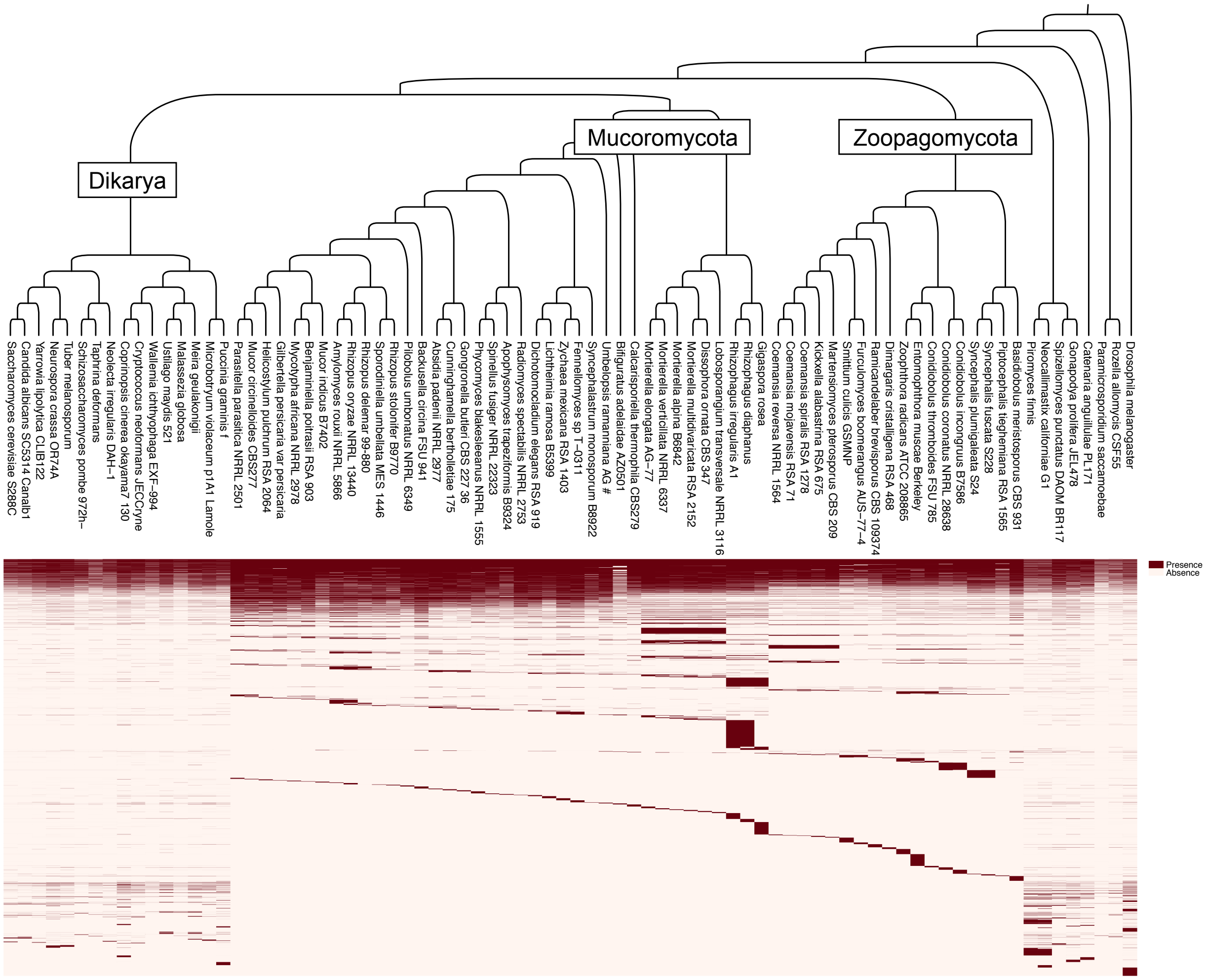

**Absence and presence of orthologous gene families across the Kingdom Fungi**

Supp. Fig. 2 Orthologous gene families across the Kingdom Fungi. A total of 62,689 gene families were found in at least one of the 80 taxa included in the backbone fungal tree of life (Fig. 2a). The dark red color indicates the presence of the gene family, while the light color represents the absence.

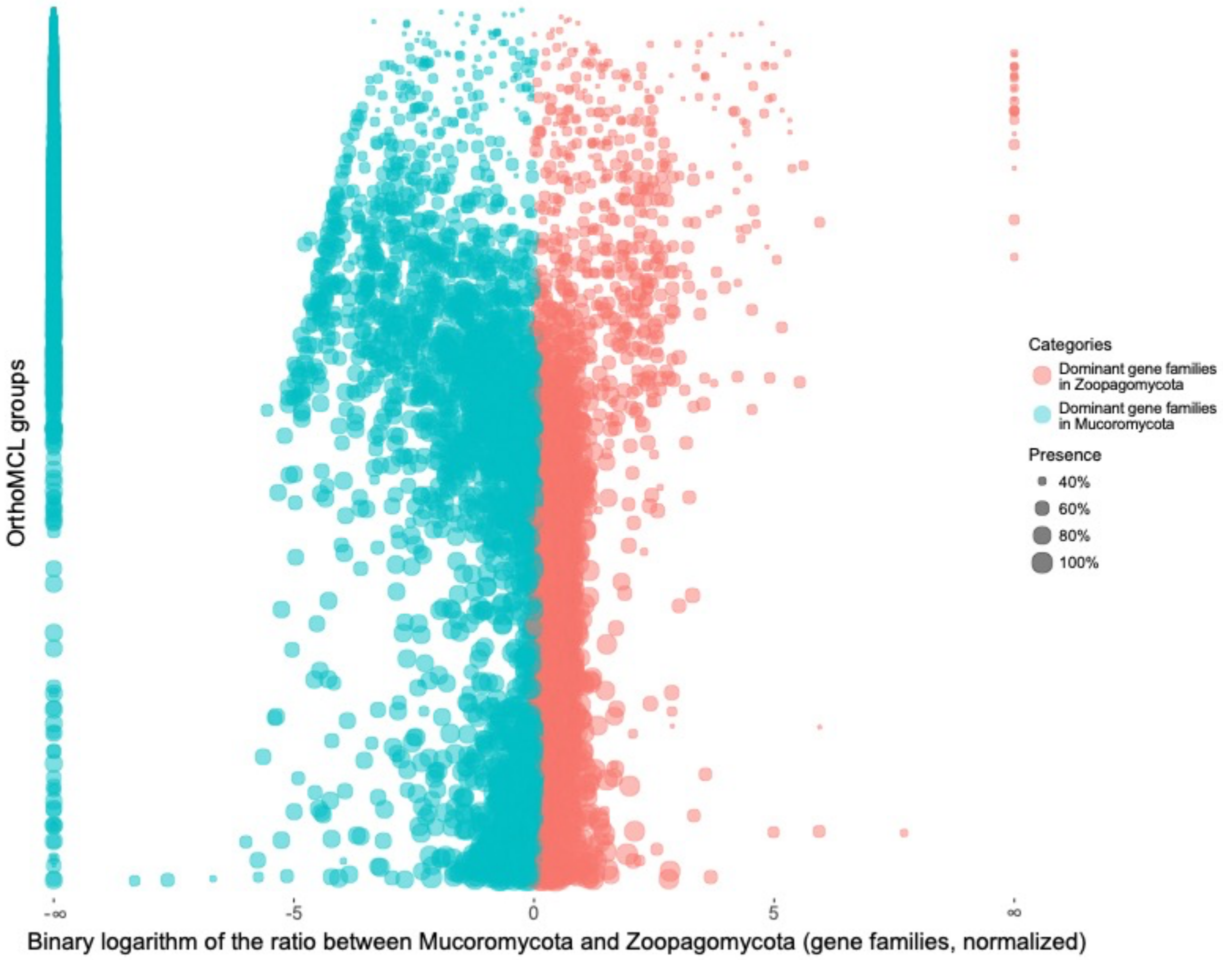

Supp. Fig. 3 Enriched gene families in Mucoromycota and Zoopagomycota. Each dot represents a gene family in zygomycete fungi. The x-axis shows the binary logarithm of the gene copy ratios between Zoopagomycota and Mucoromycota, and the y-axis shows the orthologous gene families in alphabetical order. The gene families enriched in Mucoromycota are on the left side in cyan color, and the ones enriched in Zoopagomycota are on the right side in red color. The bubbles (gene families) with bigger sizes are shared by more zygomycetes members. The dots aligned on the left edge are gene families only found in Mucoromycota and absent in Zoopagomycota, while the dots on the right edge can only be found in Zoopagomycota, but not in Mucoromycota.

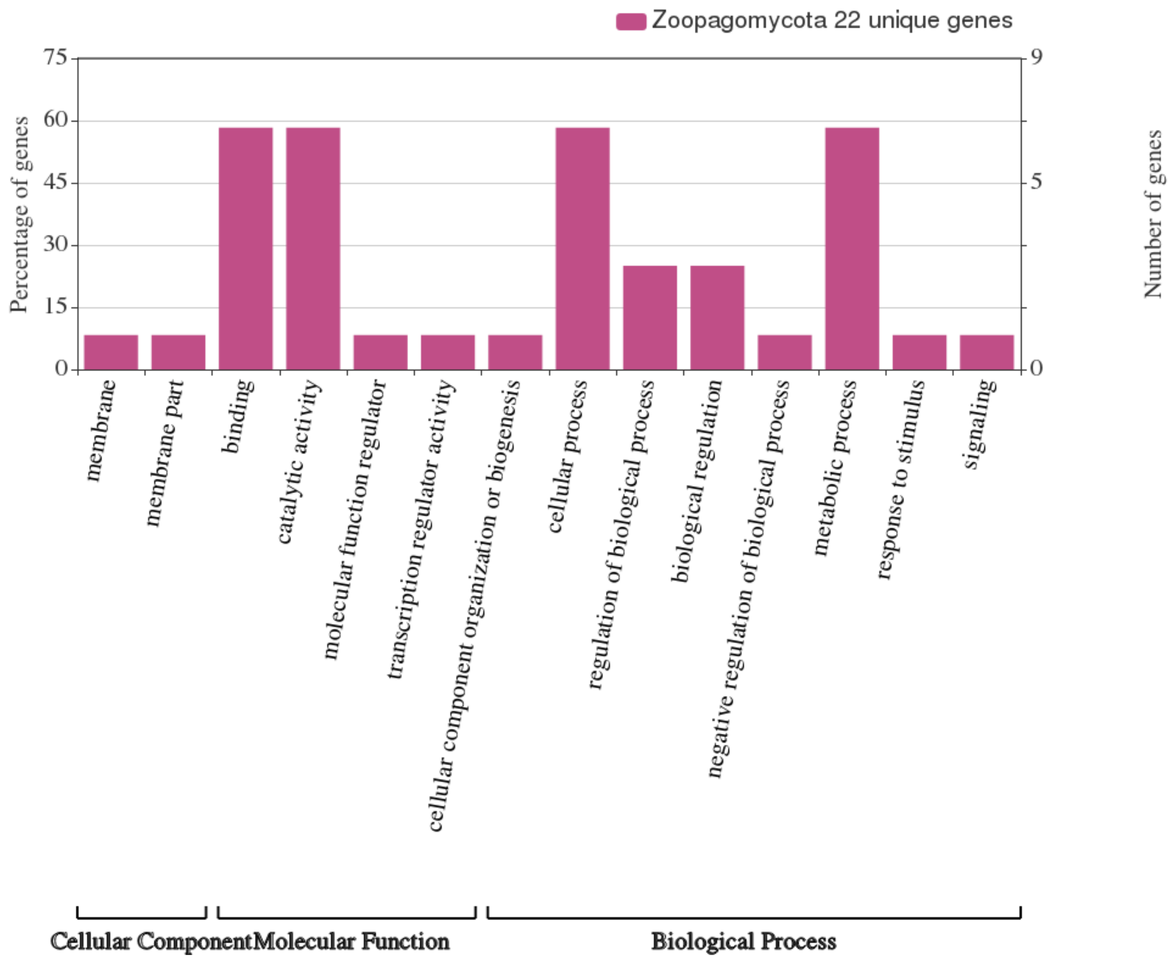

Supp. Fig. 4 Classification of Gene Ontology (GO) terms for Zoopagomycota unique genes. The x-axis indicates the sub-categories at level 2 and the y-axis shows the percentage (left) and number (right) of genes in each sub-category.

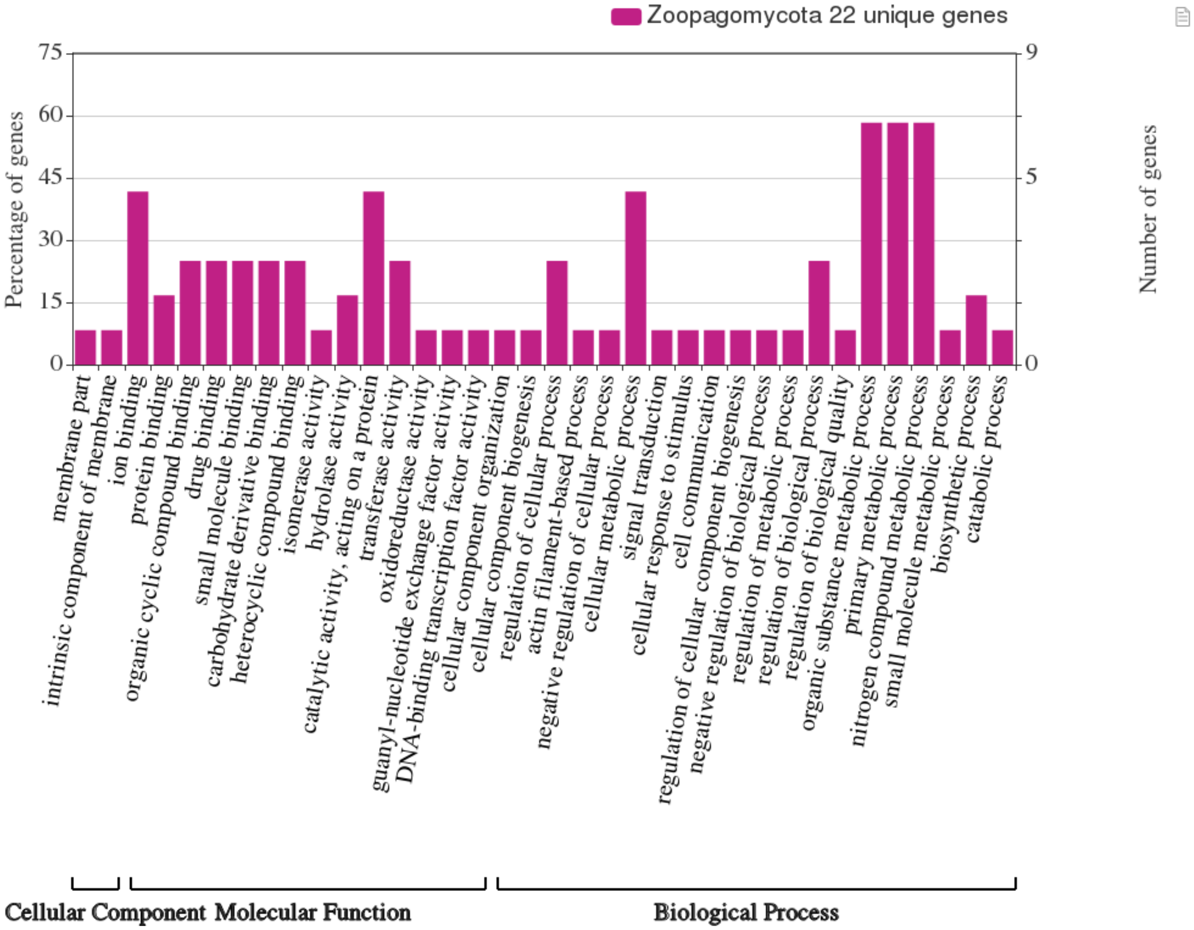

Supp. Fig. 5 A finer scale classification of GO terms for Zoopagomycota unique genes. The x-axis indicates the sub-categories at level 3 and the y-axis shows the percentage (left) and number (right) of genes in each sub-category.

Supplementary Table 1. List of non-zygomycete fungi used in the phylogenetic study in inferring kingdom-wide relationships

| Phylum | Subphylum | Species | Accession Information | DOI | Reference |
|---|---|---|---|---|---|
| Ascomycota | Saccharomycotina | Kluyveromyces lactis NRRL Y-1140 | GCF_000002515.2 | doi: 10.1038/nature02579 | Dujon et al. 2004 |
| Ascomycota | Saccharomycotina | Saccharomyces cerevisiae S288C.vR642-1 | http://yeastgenome.org | doi: 10.1126/science.274.5287.546 | Goffeau et al. 1996 |
| Ascomycota | Saccharomycotina | Cyberlindnera jadinii NRRL Y-1542 | LTAD00000000.1 | doi: 10.1073/pnas.1603941113 | Riley et al. 2016 |
| Ascomycota | Saccharomycotina | Cephaloascus albidus ATCC 66658 | https://mycocosm.jgi.doe.gov/Cepal1_1/Cepal1_1.home.html | Unpublished | |
| Ascomycota | Saccharomycotina | Candida albicans SC5314 Canalb1 | AACQ00000000.1 | doi: 10.1073/pnas.0401648101 | Jones et al. 2004 |
| Ascomycota | Saccharomycotina | Komagataella pastoris GS115 | GCA_000027005.1 | doi: 10.1038/nbt.1544 | De Schutter et al. 2009 |
| Ascomycota | Saccharomycotina | Yarrowia lipolytica CLIB122 | GCA_000002525.1 | doi: 10.1371/journal.pone.0162363 | Magnan et al. 2016 |
| Ascomycota | Saccharomycotina | Blastobotrys adeninivorans LS3 | CBZY000000000.1 | doi: 10.1186/1754-6834-7-66 | Kunze et al. 2014 |
| Ascomycota | Pezizomycotina | Xylona heveae TC161 | JXCS00000000 | doi: 10.1016/j.funbio.2015.10.002 | Gazis et al. 2016 |
| Ascomycota | Pezizomycotina | Cenococcum geophilum 1.58 | LKKR00000000.1 | doi: 10.1038/ncomms12662 | Peter et al. 2016 |
| Ascomycota | Pezizomycotina | Aspergillus niger ATCC 1015 | ACJE00000000.1 | doi: 10.1101/gr.112169.110 | Andersen et al. 2011 |
| Ascomycota | Pezizomycotina | Neurospora crassa OR74A | AABX00000000.3 | doi: 10.1038/nature01554 | Galagan et al. 2003 |
| Ascomycota | Pezizomycotina | Botrytis cinerea B05.10 | GCA_00143535.4 | doi: 10.1128/EC.00164-12 | Staats and van Kan 2012 |
| Ascomycota | Pezizomycotina | Tuber melanosporum.Tubme1 | CABJ00000000.1 | doi: 10.1038/nature08867 | Martin et al. 2010 |
| Ascomycota | Pezizomycotina | Arthrobotrys oligospora ATCC 24927 | ADOT00000000 | doi: 10.1371/journal.ppat.1002179 | Yang et al. 2011 |
| Ascomycota | Taphrinomycotina | Protomyces inouyei | https://mycocosm.jgi.doe.gov/Proin1/Proin1.home.html | Unpublished | |
| Ascomycota | Taphrinomycotina | Taphrina deformans.Tapde1 1 | CAHR00000000.2 | doi: 10.1128/mBio.00055-13 | Cissé et al. 2013 |
| Ascomycota | Taphrinomycotina | Pneumocystis jirovecii SE8 | CAKM00000000.1 | doi: 10.1128/mBio.00428-12 | Cissé et al. 2012 |
| Ascomycota | Taphrinomycotina | Schizosaccharomyces pombe 972h-.vASM294v2 | http://www.pombase.org | doi: 10.1038/nature724 | Wood et al. 2002 |
| Ascomycota | Taphrinomycotina | Saitoella complicata NRRL Y-17804 | AEUO00000000.1 | doi: 10.1073/pnas.1603941113 | Riley et al. 2016 |
| Ascomycota | Taphrinomycotina | Neolecta irregularis DAH-1.v1 | LXFE00000000 | doi: 10.1038/ncomms14444 | Nguyen et al. 2017 |
| Basidiomycota | Agaricomycotina | Ganoderma sp 10597 SS1 | ANLC00000000 | doi: 10.3852/13-003 | Binder et al. 2013 |
| Basidiomycota | Agaricomycotina | Gloeophyllum trabeum ATCC 11539 | GCA_000344685.1 | doi: 10.1126/science.1221748 | Floudas et al. 2012 |
| Basidiomycota | Agaricomycotina | Coprinopsis cinerea okayama7_130 | AACS00000000.2 | doi: 10.1073/pnas.1003391107 | Stajich et al. 2010 |
| Basidiomycota | Agaricomycotina | Ramaria rubella UT-36052-T | WIQE00000000 | doi: 10.1038/s41467-020-18795-w | Miyauchi et al. 2020 |
| Basidiomycota | Agaricomycotina | Auricularia subglabra TFB10046 SS-5 | AFVO00000000.1 | doi: 10.1126/science.1221748 | Floudas et al. 2012 |
| Basidiomycota | Agaricomycotina | Sebacina vermifera MAFF 305830 | JMDS00000000.1 | doi: 10.1038/ng.3223 | Kohler et al. 2015 |

| | | | | | |
|---|---|---|---|---|---|
| Basidiomycota | Agaricomycotina | Clavulina sp. PMI 390 | WIDE00000000.1 | doi: 10.1038/s41467-020-18795-w | Miyauchi et al. 2020 |
| Basidiomycota | Agaricomycotina | Dacryopinax sp. DJM-731 | AEUS00000000.1 | 10.1126/science.1221748 | Floudas et al. 2012 |
| Basidiomycota | Agaricomycotina | Cryptococcus neoformans JEC21 | GCA_000149245.3 | doi: 10.1126/science.1103773 | Loftus et al. 2005 |
| Basidiomycota | Wallemiomycotina | Wallemia ichthyophaga EXF-994 | APLC00000000.1 | doi: 10.1186/1471-2164-14-617 | Zajc et al. 2013 |
| Basidiomycota | Ustilaginomycotina | Sporisorium reilianum SRZ2.Spore1 | GCA_000230245.1 (ENA) | doi: 10.1126/science.1195330 | Schirawski et al. 2010 |
| Basidiomycota | Ustilaginomycotina | Ustilago maydis 521 v190413 | AACP00000000.2 | doi:10.1038/nature05248 | Kamper et al. 2006 |
| Basidiomycota | Ustilaginomycotina | Testicularia cyperi MCA 3645 | MCOH00000000.1 | doi: 10.1093/molbev/msy072 | Kijpornyongpan et al. 2018 |
| Basidiomycota | Ustilaginomycotina | Malassezia globosa | AAYY00000000.1 | doi: 10.1073/pnas.0706756104 | Xu et al. 2007 |
| Basidiomycota | Ustilaginomycotina | Meira geulakonigii | https://gold.jgi.doe.gov/project?id=Gp0120039 | Unpublished | |
| Basidiomycota | Ustilaginomycotina | Exobasidium vaccinii MPITM | https://mycocosm.jgi.doe.gov/Exova1/Exova1.home.html | Unpublished | |
| Basidiomycota | Ustilaginomycotina | Jaminaea rosea MCA 5214 | MCHB00000000.1 | doi: 10.1093/molbev/msy072 | Kijpornyongpan et al. 2018 |
| Basidiomycota | Pucciniomycotina | Microbotryum violaceum p1A1 Lamole | AEIJ00000000.1 | doi: 10.1186/s12864-015-1660-8 | Perlin et al. 2015 |
| Basidiomycota | Pucciniomycotina | Rhodosporidium toruloides NP11 | ALAU00000000.1 | doi: 10.1038/ncomms2112 | Zhu et al. 2012 |
| Basidiomycota | Pucciniomycotina | Atractiellales sp PMI 95 | https://mycocosm.jgi.doe.gov/Atrsp2/Atrsp2.home.html | Unpublished | |
| Basidiomycota | Pucciniomycotina | Puccinia graminis f. sp. tritici CRL 75-36-700-3 | AAWC0000000.1 | doi: 10.1073/pnas.1019315108 | Duplessis et al. 2011 |
| Basidiomycota | Pucciniomycotina | Tritirachium sp CBS 265.96 | https://mycocosm.jgi.doe.gov/Trisp1/Trisp1.home.html | Unpublished | |
| Chytridiomycota | N/A | Spizellomyces punctatus DAOM BR117.v1 | ACOE00000000.1 | doi: 10.1128/genomeA.00849-16 | Russ et al. 2016 |
| Chytridiomycota | N/A | Gonapodya prolifera JEL478 | LSZK00000000 | doi: 10.1093/gbe/evv090 | Chang et al. 2015 |
| Blastocladiomycota | N/A | Catenaria anguillulae PL171 | MCFL00000000.1 | doi: 10.1038/ng.3859 | Mondo et al. 2017 |
| Neocallimastigomycota | N/A | Piromyces finnis | MCFH00000000.1 | doi: 10.1038/nmicrobiol.2017.87; doi: 10.1038/ng.3859 | Haitjema et al. 2017; Mondo et al. 2017 |
| Neocallimastigomycota | N/A | Neocallimastix californiae G1.Neosp1 | MCOG00000000.1 | doi: 10.1038/nmicrobiol.2017.87 | Haitjema et al. 2017 |
| Cryptomycota | N/A | Paramicrosporidium saccamoebae KSL3 v6 | MTSL00000000.1 | doi: 10.7554/eLife.29594 | Quandt et al. 2017 |
| Cryptomycota | N/A | Rozella allomycis CSF55 | ATJD00000000.1 | doi: 10.1016/j.cub.2013.06.057 | James et al. 2013 |
| Arthropoda | N/A | Drosophila melanogaster.vr6.04 | http://flybase.org | doi:10.1126/science.287.5461.2185 | Adams et al. 2000 |

Supplementary Table 2. List of phylum-specific and subphylum-specific Pfam domains in zygomycete fungi

| Phylum-level (>10 taxa) | | Subphylum-level (>1 taxa) | | | | | |
|---|---|---|---|---|---|---|---|
| Mucoromycota | Zoopagomycota | Mucoromycotina | Mortierellomycotina | Glomeromycotina | Kickxellomycotina | Entomorphthoromycotina | Zoopagomycotina |
| C9orf72-like | N/A | Adeno_E3_CR2 | BTRD1 | Anemone_cytotox | N/A | Carbam_trans_N | DUF664 |
| CheR | | Agenet | DUF4142 | Chol_subst-bind | | GH3 | |
| | | BH3 | DUF553 | Colicin_D | | IspA | |
| | | Cytochrom_D1 | DUF655 | cpYpsA | | N6_N4_Mtase | |
| | | Dickkopf_N | GDPD_2 | Cytotoxic | | Toxin_30 | |
| | | DUF1838 | HEPN_DZIP3 | DUF2283 | | | |
| | | DUF1864 | Phage_T4_gp19 | DUF3161 | | | |
| | | DUF4070 | Pox_G5 | DUF393 | | | |
| | | DUF4611 | RHSP | DUF43 | | | |
| | | DUF711 | STELLO | DUF4371 | | | |
| | | FG-GAP_2 | YscW | DUF4379 | | | |
| | | GHL10 | | Endonuclease_7 | | | |
| | | Glyco_hydro_52 | | GNAT_C | | | |
| | | Herpes_gE | | gpD | | | |
| | | HTS | | HSDR_N | | | |
| | | Lactate_perm | | HTH_OrfB_IS605 | | | |
| | | NCU-G1 | | LXG | | | |
| | | NinF | | Methyltrn_RNA_4 | | | |
| | | NQRA | | Mrr_cat_2 | | | |
| | | Peptidase_C10 | | Phage_F | | | |
| | | Phage_int_SAM_1 | | T4_deiodinase | | | |
| | | potato_inhibit | | T4_Rnl2_C | | | |
| | | PTS-HPr | | TerD | | | |
| | | RAG2 | | TetR_N | | | |
| | | SUIM_assoc | | | | | |
| | | TaqI_C | | | | | |
| | | TcdA_TcdB | | | | | |
| | | Toprim_2 | | | | | |
| | | Toxin_38 | | | | | |
| | | Transposase_31 | | | | | |
| | | zf-3CxxC_2 | | | | | |
| | | zf-TRAF_2 | | | | | |

Supplementary Table 3. List of Pfam domains that are either abundant in Mucoromycota (with negative binary logarithm values) or Zoopagomycota (with positive binary logarithm values)

| Pfam domain name | Binary logarithm | Number of zygomycete fungi containing this domain (out of 56) | Description |
|---|---|---|---|
| Tht1 | -10 | 20 | Tht1-like nuclear fusion protein |
| Glyco_hydro_36C | -10 | 20 | Glycosyl hydrolase family 36 C-terminal domain |
| Glyco_hydro_45 | -10 | 21 | Glycoside hydrolase family 45 |
| Glyco_transf_17 | -10 | 21 | Glycosyltransferase family 17 |
| Alpha_L_fucos | -10 | 21 | Alpha-L-fucosidase |
| Lyase_8_C | -10 | 21 | Polysaccharide lyase family 8, C-terminal beta-sandwich domain |
| DUF2151 | -10 | 21 | Cell cycle and development regulator |
| DDRGK | -10 | 21 | DDRGK |
| zf-CpG_bind_C | -10 | 21 | CpG binding protein zinc finger C terminal domain |
| Glyco_hydro_28 | -10 | 22 | Glycosyl hydrolases family 28 |
| Glyco_transf_64 | -10 | 22 | Glycosyl transferase family 64 domain |
| Pox_ser-thr_kin | -10 | 22 | Poxvirus serine/threonine protein kinase |
| Pur_ac_phosph_N | -10 | 23 | Purple acid Phosphatase, N-terminal domain |
| Lyase_8 | -10 | 23 | Polysaccharide lyase family 8, super-sandwich domain |
| Lyase_8_N | -10 | 23 | Polysaccharide lyase family 8, N terminal alpha-helical domain |
| DUF1765 | -10 | 23 | Protein of unknown function (DUF1765) |
| Abp2 | -10 | 23 | autonomously replicating sequence (ARS) binding protein 2 |
| DUF1838 | -10 | 23 | Protein of unknown function (DUF1838) |
| S6PP | -10 | 24 | Sucrose-6F-phosphate phosphohydrolase |
| Metallophos_C | -10 | 24 | Iron/zinc purple acid phosphatase-like protein C |
| Methyltransf_29 | -10 | 24 | Putative S-adenosyl-L-methionine-dependent methyltransferase |
| CHDNT | -10 | 25 | CHDNT (NUC034) domain found in PHD/RING finger and chromo domain-associated helicases |
| DUF3455 | -10 | 25 | Protein of unknown function (DUF3455) |
| Glyco_hydro_36N | -10 | 25 | Glycosyl hydrolase family 36 N-terminal domain |
| FANCI_S3 | -10 | 25 | FANCI solenoid 3 |
| MR_MLE_N | -10 | 26 | Mandelate racemase / muconate lactonizing enzyme, N-terminal domain |
| Peptidase_M19 | -10 | 27 | Membrane dipeptidase (Peptidase family M19) |
| MTABC_N | -10 | 27 | Mitochondrial ABC-transporter N-terminal five TM region |
| MIOX | -10 | 27 | Myo-inositol oxygenase |
| HutD | -10 | 27 | HutD from Pseudomonas fluorescens SBW25 is a component of the histidine uptake and utilisation operon |
| DUF2834 | -10 | 27 | Protein of unknown function (DUF2834) |
| Lactate_perm | -10 | 28 | L-lactate permease |
| Methyltransf_21 | -10 | 28 | Methyltransferase FkbM domain |
| PC-Esterase | -10 | 28 | GDSL/SGNH-like Acyl-Esterase family found in Pmr5 and Cas1p |
| SASA | -10 | 28 | Carbohydrate esterase, sialic acid-specific acetylesterase |
| ERG2_Sigma1R | -10 | 28 | ERG2 and Sigma1 receptor like protein |
| PTPlike_phytase | -10 | 29 | Protein tyrosine phosphatase |
| Gln-synt_N | -10 | 29 | Glutamine synthetase, beta-Grasp domain |
| SHR3_chaperone | -10 | 29 | ER membrane protein SH3 |
| DUF1741 | -10 | 29 | Domain of unknown function (DUF1741) |
| E3_UFM1_ligase | -10 | 29 | E3 UFM1-protein ligase 1 |
| Nuc_H_symport | -10 | 30 | Nucleoside H+ symporter |
| Kei1 | -10 | 30 | Inositolphosphorylceramide synthase subunit Kei1 |
| C9orf72-like | -10 | 30 | C9orf72-like protein family |
| ASXH | -10 | 31 | Asx homology domain |
| Fmp27_SW | -10 | 31 | RNA pol II promoter Fmp27 protein domain |
| DUF3844 | -10 | 31 | Domain of unknown function (DUF3844) |
| uDENN | -10 | 32 | uDENN domain, regulators of practically all membrane trafficking events in eukaryotes |
| TM2 | -10 | 32 | TM2 domain (a pair of transmembrane alpha helices connected by a short linker) |
| DUF2461 | -10 | 32 | Conserved hypothetical protein (DUF2461) |
| Exostosin | -10 | 33 | Exostosin family |
| NCBP3 | -10 | 33 | Nuclear cap-binding protein subunit 3 |
| NT5C | -10 | 33 | 5' nucleotidase, deoxy (Pyrimidine), cytosolic type C protein (NT5C) |
| PLDc_N | -10 | 34 | Phospholipase_D-nuclease N-terminal |
| dDENN | -10 | 34 | dDENN domain, regulators of practically all membrane trafficking events in eukaryotes |
| ANF_receptor | -10 | 35 | Receptor family ligand binding region |
| DENN | -10 | 35 | DENN (AEX-3) domain, regulators of practically all membrane trafficking events in eukaryotes |
| FUN14 | -10 | 35 | FUN14 family |
| DUF1691 | -10 | 35 | Protein of unknown function (DUF1691) |
| NUC194 | -10 | 35 | NUC194 domain |
| SMG1 | -10 | 35 | Serine/threonine-protein kinase smg-1, a key regulator of growth (loss of SMG1 leads to hyperactive responses to injury and subsequent growth that continues out of control) |
| SUR7 | -10 | 36 | SUR7/PalI family |
| Peripla_BP_6 | -10 | 36 | Periplasmic binding protein |
| DUF3425 | -10 | 37 | Domain of unknown function (DUF3425) |
| RTC4 | -10 | 37 | RTC4-like domain |
| E2F_TDP | -10 | 37 | E2F/DP family winged-helix DNA-binding domain |
| Melibiase_2 | -10 | 37 | Glycoside hydrolase family 27 |
| Melibiase_C | -10 | 37 | Alpha galactosidase C-terminal beta sandwich domain |
| PRA1 | -10 | 37 | PRA1 family protein (Prenylated rab acceptor) |
| Pil1 | -10 | 38 | Eisosome component PIL1 |
| CotH | -7.286497299 | 34 | |
| SBF_like | -6.226502813 | 35 | |
| TrkH | -5.717997045 | 30 | |
| Smg8_Smg9 | -5.673941017 | 36 | |
| PAP1 | -5.554790806 | 39 | |
| DUF4442 | -5.366717472 | 39 | |
| DUF2235 | -5.187414783 | 27 | |
| WRKY | -5.056959876 | 37 | |
| An_peroxidase | -4.996401753 | 31 | |
| DNA_pol_lambd_f | -4.91887188 | 33 | |
| RicinB_lectin_2 | -4.888969948 | 40 | |
| Cons_hypoth698 | -4.835212988 | 36 | |
| DNA_pol_B_thumb | -4.795091121 | 29 | |
| DNA_pol_B_palm | -4.784545885 | 28 | |
| HHH_8 | -4.780627268 | 30 | |
| Senescence | -4.687956684 | 31 | |
| Sulfotransfer_4 | -4.647708709 | 28 | |
| LRR_6 | -4.645629139 | 54 | |
| EHN | -4.604049901 | 36 | |

| | | |
|---|---|---|
| Melibiase | -4.598284062 | 39 |
| Caleosin | -4.571681845 | 36 |
| Nucleotid_trans | -4.540551321 | 34 |
| MMPL | -4.478953458 | 34 |
| UPF0052 | -4.478535182 | 37 |
| Gb3_synth | -4.462587203 | 33 |
| DNA_ligase_OB_2 | -4.41018714 | 34 |
| KAR9 | -4.401188319 | 36 |
| DUF2205 | -4.36890957 | 34 |
| DDE_Tnp_4 | -4.36285472 | 20 |
| Ricin_B_lectin | -4.33881613 | 40 |
| YBD | -4.298446015 | 34 |
| Reticulon | -4.257807461 | 39 |
| DHDPS | -4.255537848 | 28 |
| PAC1 | -4.225595845 | 31 |
| zf-primase | -4.208463369 | 29 |
| CK2S | -4.189541707 | 24 |
| DUF1264 | -4.133846046 | 26 |
| Med12-LCEWAV | -4.128572188 | 28 |
| Cu_amine_oxid | -4.086097934 | 29 |
| DUF5321 | -4.081428992 | 28 |
| Glyco_transf_10 | -4.068451435 | 33 |
| Glyco_transf_49 | -4.02260694 | 42 |
| BACK | -3.977097915 | 28 |
| Cu_amine_oxidN3 | -3.963460282 | 28 |
| RTP1_C2 | -3.909464203 | 24 |
| Sulfotransfer_1 | -3.893958683 | 21 |
| GET2 | -3.874480402 | 32 |
| Haem_degrading | -3.862539271 | 37 |
| Cu_amine_oxidN2 | -3.842783119 | 28 |
| zf-RVT | -3.800417775 | 31 |
| DUF2838 | -3.782808655 | 35 |
| ADP_ribosyl_GH | -3.778272854 | 37 |
| EOS1 | -3.762564422 | 39 |
| Med5 | -3.746089138 | 26 |
| LINES_N | -3.734042083 | 23 |
| Dis3l2_C_term | -3.691840331 | 42 |
| DUF273 | -3.680657901 | 27 |
| Gemin6 | -3.676618296 | 23 |
| PFOR_II | -3.644888585 | 29 |
| AATase | -3.610170789 | 34 |
| MHYT | -3.606884684 | 29 |
| AGTRAP | -3.565556823 | 29 |
| DDE_3 | -3.538907589 | 27 |
| Ku_PK_bind | -3.518925901 | 36 |
| MBOAT_2 | -3.509812699 | 42 |
| MTP18 | -3.478885758 | 35 |
| CDKN3 | -3.459035728 | 30 |
| Thaumatin | -3.434057239 | 29 |
| Nup188 | -3.419590912 | 41 |
| GAS2 | -3.417688952 | 43 |
| O-FucT | -3.396180147 | 41 |
| Exo_endo_phos_2 | -3.372298451 | 29 |
| DUF1929 | -3.352738952 | 40 |
| Polysacc_synt_C | -3.311655319 | 34 |
| Glyoxal_oxid_N | -3.288178467 | 40 |
| Acetyltransf_CG | -3.259159846 | 23 |
| AcetylCoA_hydro | -3.23062002 | 31 |
| MID_MedPIWI | -3.213467562 | 40 |
| SOG2 | -3.181559184 | 36 |
| GCR1_C | -3.177321651 | 33 |
| BTB_2 | -3.129516934 | 42 |
| OATP | -3.113200129 | 27 |
| SWIM | -3.104829042 | 34 |
| AtuA | -3.075008893 | 37 |
| NACHT | -3.071980364 | 31 |
| BetaGal_dom4_5 | -3.055401 | 30 |
| DUF4396 | -3.04743991 | 27 |
| Dpy-30 | -3.042288187 | 30 |
| Dsl1_C | -2.99807987 | 27 |
| Tubulin-binding | -2.995416288 | 35 |
| XLF | -2.969347067 | 24 |
| NUP214 | -2.958766567 | 24 |
| Eco57I | -2.943475117 | 25 |
| Frizzled | -2.94136386 | 21 |
| GRAB | -2.900746103 | 27 |
| OSR1_C | -2.896705303 | 30 |
| Rubis-subs-bind | -2.869564717 | 36 |
| CTD | -2.864876134 | 37 |
| RBFA | -2.85116806 | 21 |
| AcetylCoA_hyd_C | -2.844305915 | 32 |
| Phi_1 | -2.83665619 | 29 |
| PH_4 | -2.806192692 | 33 |
| NAS | -2.802699345 | 23 |
| Preseq_ALAS | -2.797216806 | 21 |
| HSBP1 | -2.793410214 | 24 |
| Aminotran_MocR | -2.780033515 | 39 |
| DUF900 | -2.764699475 | 28 |
| Peptidase_M13 | -2.744188471 | 42 |
| BTB | -2.731237214 | 53 |
| TFIIE-A_C | -2.727724247 | 22 |
| FAM178 | -2.705363932 | 24 |
| Peptidase_M13_N | -2.69081016 | 42 |
| LtrA | -2.679109785 | 41 |
| DOCK-C2 | -2.678741433 | 34 |
| DUF262 | -2.659991323 | 25 |
| DUF1687 | -2.624936508 | 41 |
| Promethin | -2.623763161 | 21 |

| | | |
|---|---|---|
| UvdE | -2.612726657 | 25 |
| DUF4449 | -2.601243127 | 37 |
| Gryzun-like | -2.567802269 | 20 |
| ABC_trans_N | -2.556172748 | 39 |
| COG7 | -2.546555374 | 39 |
| FHA_2 | -2.546279453 | 28 |
| AIG1 | -2.530662572 | 46 |
| TLD | -2.495658461 | 56 |
| DUF1206 | -2.48677723 | 29 |
| Integrase_H2C2 | -2.479397378 | 44 |
| zf-PARP | -2.47543716 | 37 |
| TTKRSYEDQ | -2.474521796 | 35 |
| Presenilin | -2.471226841 | 39 |
| Lip_prot_lig_C | -2.462793795 | 35 |
| eIF_4G1 | -2.462302573 | 34 |
| RPE65 | -2.411496236 | 48 |
| Mannosyl_trans3 | -2.408725588 | 38 |
| Catalase_C | -2.395082609 | 30 |
| MFS_1_like | -2.376235464 | 47 |
| JHD | -2.376127564 | 39 |
| RVT_1 | -2.360528505 | 24 |
| DOCK_N | -2.345225784 | 35 |
| Glyco_transf_28 | -2.343056719 | 44 |
| N-SET | -2.335749394 | 30 |
| DUF563 | -2.318735497 | 32 |
| Lactonase | -2.316243114 | 30 |
| DUF5614 | -2.31500336 | 22 |
| ArsC | -2.296940799 | 33 |
| Thi4 | -2.287009055 | 37 |
| AAA_26 | -2.283677419 | 32 |
| UBA_2 | -2.266741654 | 36 |
| PhosphMutase | -2.26032568 | 27 |
| Glu_dehyd_C | -2.243391741 | 30 |
| GAF | -2.193058352 | 46 |
| PAS_4 | -2.177380459 | 41 |
| Ku_C | -2.168033138 | 36 |
| VTC | -2.159754528 | 54 |
| Stk19 | -2.125461961 | 34 |
| OCD_Mu_crystall | -2.118542739 | 39 |
| SBF | -2.112232018 | 41 |
| Sel1 | -2.112059837 | 56 |
| Methyltransf_2 | -2.110069438 | 38 |
| TIG | -2.102835953 | 52 |
| DUF1752 | -2.09280394 | 50 |
| Ins134_P3_kin | -2.078759601 | 30 |
| SLS | -2.077021862 | 37 |
| YTH | -2.0688735 | 26 |
| zf-C4H2 | -2.061364343 | 26 |
| RT_RNaseH_2 | -2.059725664 | 20 |
| DUF2362 | -2.047936377 | 44 |
| Ser_hydrolase | -2.047363865 | 26 |
| LacY_symp | -2.036295371 | 30 |
| Retrotrans_gag | -2.027861857 | 29 |
| UvrD_C_2 | -2.012592786 | 23 |
| ArAE_2_N | -2.003175948 | 50 |
| RBP_receptor | -2.001432596 | 31 |
| TRAP_alpha | -1.971213733 | 39 |
| BAF250_C | -1.967418662 | 36 |
| KIF1B | -1.962489695 | 45 |
| zf-TRAF | -1.960834099 | 36 |
| CTC1 | -1.958064347 | 28 |
| BATS | -1.956783108 | 42 |
| TMEM18 | -1.934805101 | 34 |
| TMEM164 | -1.933494528 | 40 |
| REV1_C | -1.933334657 | 25 |
| Lectin_legB | -1.930237472 | 29 |
| MAP65_ASE1 | -1.923820429 | 42 |
| WHIM1 | -1.900468623 | 32 |
| TFIIIC_delta | -1.890230126 | 33 |
| TCO89 | -1.887474959 | 38 |
| DUF1771 | -1.877499691 | 47 |
| F-box-like | -1.870264397 | 56 |
| PQQ_2 | -1.865430064 | 47 |
| DUF5102 | -1.860682042 | 37 |
| Chalcone_2 | -1.859978529 | 43 |
| ChlI | -1.85554578 | 43 |
| Kinetochor_Ybp2 | -1.852096499 | 36 |
| Glyco_transf_5 | -1.844336203 | 48 |
| RNA_ligase | -1.840097023 | 39 |
| Methyltransf_8 | -1.836587944 | 56 |
| Thioredox_DsbH | -1.831334515 | 28 |
| TAXi_N | -1.831304031 | 56 |
| CSTF1_dimer | -1.824351082 | 24 |
| CENP-P | -1.820222545 | 22 |
| GRDP-like | -1.818167139 | 26 |
| Glyco_hydro_5_C | -1.817075152 | 44 |
| FAD_binding_3 | -1.815905619 | 56 |
| ArAE_2 | -1.812729507 | 50 |
| RT_RNaseH | -1.811446612 | 22 |
| Asp | -1.809427916 | 56 |
| RdRP | -1.808236186 | 44 |
| zf-NADH-PPase | -1.802814138 | 28 |
| DUF3361 | -1.794758853 | 35 |
| Alginate_lyase | -1.792760471 | 41 |
| DRY_EERY | -1.787784136 | 40 |
| Fer2_BFD | -1.782046864 | 21 |
| DUF3712 | -1.778506476 | 49 |
| Ncstrn_small | -1.771805901 | 39 |

| | | |
|---|---|---|
| Anp1 | -1.771308259 | 40 |
| Hemerythrin | -1.768302465 | 20 |
| Zeta_toxin | -1.768168788 | 22 |
| SPOC | -1.767765149 | 36 |
| Avl9 | -1.757532033 | 47 |
| CRT10 | -1.753629293 | 30 |
| EI24 | -1.749911303 | 47 |
| FUSC_2 | -1.748631522 | 54 |
| DUF3759 | -1.748472429 | 50 |
| SPARC_Ca_bdg | -1.746595878 | 42 |
| Glyco_hydro_85 | -1.744583921 | 28 |
| 7tm_2 | -1.741757591 | 28 |
| DUSP | -1.736458534 | 41 |
| Glyco_transf_34 | -1.717610449 | 40 |
| DUF3808 | -1.713729963 | 55 |
| NST1 | -1.713235363 | 39 |
| Glyco_hydro_9 | -1.707507961 | 42 |
| PEN-2 | -1.706575318 | 30 |
| Tropomyosin_1 | -1.702022624 | 55 |
| Cyclase | -1.701134716 | 32 |
| Importin_rep | -1.696580519 | 27 |
| PTEN_C2 | -1.695168383 | 23 |
| Aph-1 | -1.692254001 | 38 |
| PLDc | -1.688772178 | 55 |
| ATG11 | -1.685519015 | 32 |
| LysM | -1.68353842 | 41 |
| Glyco_hydro_125 | -1.676801068 | 36 |
| DUF872 | -1.676153724 | 38 |
| Sec15 | -1.675081211 | 41 |
| Fra10Ac1 | -1.670406149 | 42 |
| NDT80_PhoG | -1.655177266 | 55 |
| DUF1772 | -1.640762817 | 38 |
| Mtc | -1.639691923 | 36 |
| TRAM | -1.616543052 | 30 |
| VIT1 | -1.608068564 | 37 |
| NMT1 | -1.604937951 | 43 |
| D-ser_dehydrat | -1.601022352 | 27 |
| Gryzun | -1.59328004 | 42 |
| CYSTM | -1.588069346 | 25 |
| Fe_hyd_SSU | -1.577966788 | 35 |
| VASt | -1.577890145 | 54 |
| Ppx-GppA | -1.552651327 | 45 |
| GAT | -1.551070313 | 52 |
| Acetyltransf_4 | -1.547823347 | 42 |
| CHZ | -1.545354221 | 32 |
| GRIP | -1.536154384 | 41 |
| TIMELESS_C | -1.528465535 | 24 |
| SDH_C | -1.52818949 | 34 |
| DUF2470 | -1.517282118 | 42 |
| Glyco_hydro_8 | -1.516388971 | 38 |
| FUSC | -1.515426858 | 51 |
| Peptidase_C97 | -1.514977248 | 50 |
| HUN | -1.512521234 | 36 |
| Flavin_Reduct | -1.492829438 | 40 |
| Xan_ur_permease | -1.485026374 | 49 |
| DUF810 | -1.484319354 | 35 |
| PIF1 | -1.475531278 | 45 |
| DUF2046 | -1.473299597 | 46 |
| PABP | -1.463796461 | 56 |
| ALMT | -1.455211135 | 53 |
| Hamartin | -1.452492457 | 29 |
| DUF4704 | -1.448368957 | 33 |
| FH2 | -1.444216108 | 56 |
| Vac7 | -1.437215682 | 43 |
| TauD | -1.433599431 | 54 |
| DNA_pol_delta_4 | -1.431491073 | 30 |
| CDC24_OB2 | -1.426095186 | 32 |
| SIKE | -1.425858916 | 22 |
| Polysacc_deac_1 | -1.424123267 | 48 |
| Cas1_AcylT | -1.415234961 | 40 |
| RED_C | -1.411633109 | 42 |
| GAF_2 | -1.4085318 | 46 |
| Acid_PPase | -1.404991206 | 29 |
| Amidohydro_2 | -1.40157158 | 47 |
| HTH_23 | -1.401438016 | 23 |
| RIIa | -1.400086356 | 47 |
| F-box | -1.393304114 | 56 |
| DUF1688 | -1.391601762 | 48 |
| CHIP_TPR_N | -1.389983131 | 34 |
| Ribosomal_L50 | -1.384265682 | 36 |
| TFR_dimer | -1.383633918 | 39 |
| DUF2433 | -1.38155525 | 44 |
| DDT | -1.380170992 | 48 |
| LigB | -1.380045133 | 43 |
| DUF1308 | -1.372435974 | 40 |
| AAA_PrkA | -1.371454941 | 51 |
| HI0933_like | -1.361479996 | 35 |
| Phosphoesterase | -1.350830139 | 40 |
| PXB | -1.350570391 | 44 |
| PH_5 | -1.342568761 | 56 |
| Fer4_NifH | -1.33799764 | 23 |
| FMO-like | -1.332569833 | 51 |
| Methyltransf_9 | -1.332151387 | 48 |
| NicO | -1.330460249 | 31 |
| HisKA | -1.329316442 | 54 |
| Gly_transf_sug | -1.319479543 | 43 |
| Pik1 | -1.31224225 | 45 |
| Suc_Fer-like | -1.308911354 | 53 |

| | | |
|---|---|---|
| DUF3694 | -1.307958015 | 53 |
| DUF4807 | -1.305520996 | 29 |
| BAR_3 | -1.302814927 | 51 |
| PAH | -1.301168408 | 56 |
| Drf_GBD | -1.298285909 | 56 |
| Ribosomal_L6e_N | -1.296549233 | 44 |
| Cas_Cas4 | -1.291942216 | 32 |
| COesterase | -1.289458129 | 55 |
| DUF4061 | -1.283842669 | 20 |
| DUF2828 | -1.27787561 | 33 |
| DNA_repr_REX1B | -1.274581908 | 38 |
| PA26 | -1.274509626 | 41 |
| Ku_N | -1.271239164 | 45 |
| Ribosomal_S27 | -1.267634622 | 48 |
| START | -1.263917031 | 54 |
| DDHD | -1.257895513 | 52 |
| DMAP1 | -1.256089511 | 39 |
| Linker_histone | -1.253802251 | 47 |
| PLA2_B | -1.251541966 | 48 |
| RNA_Me_trans | -1.249502695 | 43 |
| Chitin_synth_1 | -1.248615411 | 56 |
| Rubrerythrin | -1.247486581 | 41 |
| NUDIX-like | -1.243924025 | 40 |
| LON_substr_bdg | -1.243621819 | 55 |
| B12D | -1.237984368 | 54 |
| DUF1358 | -1.229227144 | 22 |
| Tropomyosin | -1.228167892 | 28 |
| DUF1479 | -1.223361015 | 21 |
| Ubie_methyltran | -1.222245523 | 56 |
| CP2 | -1.221079376 | 53 |
| TPR_12 | -1.220977088 | 55 |
| AidB_N | -1.218777311 | 35 |
| Ost5 | -1.218731713 | 36 |
| Sec2p | -1.217923409 | 56 |
| BPL_C | -1.217231764 | 30 |
| UAE_UbL | -1.20962825 | 23 |
| Ribosomal_L40e | -1.204676831 | 31 |
| Chitin_synth_1N | -1.19511176 | 56 |
| POR_N | -1.190147942 | 46 |
| Methyltransf_4 | -1.185454139 | 56 |
| DNA_ligase_A_M | -1.18515119 | 55 |
| DUF3818 | -1.185003801 | 45 |
| Transglut_core | -1.183083687 | 42 |
| VanZ | -1.181549784 | 25 |
| Response_reg | -1.179121494 | 56 |
| BCS1_N | -1.175383838 | 56 |
| Tmemb_170 | -1.175173375 | 38 |
| LEA_2 | -1.174372456 | 28 |
| DUF2347 | -1.172816166 | 42 |
| Copper-fist | -1.172395854 | 54 |
| MOR2-PAG1_mid | -1.161465637 | 47 |
| Gpr1_Fun34_YaaH | -1.160646753 | 46 |
| Glyco_transf_21 | -1.160597308 | 52 |
| F_bP_aldolase | -1.159817882 | 47 |
| Kelch_5 | -1.156182411 | 56 |
| UNC45-central | -1.156180674 | 43 |
| Smr | -1.152977244 | 52 |
| Lipin_mid | -1.14538294 | 41 |
| DUF3512 | -1.144871359 | 22 |
| Stc1 | -1.136530181 | 37 |
| IIGP | -1.135670098 | 50 |
| MetW | -1.13112453 | 41 |
| Nucleoporin_FG2 | -1.126997822 | 21 |
| TOM20_plant | -1.119123888 | 28 |
| PAP_central | -1.117754769 | 54 |
| Chromo | -1.117023237 | 56 |
| Kelch_3 | -1.11136169 | 56 |
| Aft1_OSA | -1.108625633 | 46 |
| DUF4504 | -1.103683813 | 30 |
| Trs65 | -1.10358564 | 40 |
| Drf_FH3 | -1.101081722 | 56 |
| Cnn_1N | -1.100452094 | 42 |
| Heme_oxygenase | -1.098559367 | 43 |
| Calponin | -1.097363662 | 28 |
| Nup35_RRM_2 | -1.095321918 | 23 |
| FTR1 | -1.093630659 | 44 |
| COG5 | -1.089533174 | 46 |
| bZIP_1 | -1.086196397 | 56 |
| ParBc | -1.086074669 | 28 |
| HLH | -1.084061842 | 56 |
| DHR-2 | -1.083826845 | 36 |
| DUF5600 | -1.078371326 | 50 |
| SAP18 | -1.076014479 | 46 |
| Kelch_4 | -1.075810758 | 56 |
| ACT_7 | -1.075426763 | 55 |
| RXT2_N | -1.075095234 | 42 |
| Vezatin | -1.07404463 | 49 |
| Ribosomal_L15e | -1.071260729 | 55 |
| Rad54_N | -1.070063177 | 41 |
| KLRAQ | -1.06984809 | 40 |
| SHD1 | -1.068329374 | 54 |
| SET_assoc | -1.065370298 | 21 |
| B56 | -1.062805396 | 56 |
| HOOK | -1.058524211 | 52 |
| TRAPPC10 | -1.057117871 | 40 |
| PIH1 | -1.054469617 | 42 |
| Ku | -1.05094087 | 47 |
| NodS | -1.04724251 | 34 |

| | | |
|---|---|---|
| USP7_C2 | -1.046873669 | 55 |
| Syntaxin_2 | -1.043336034 | 54 |
| TPR_10 | -1.039330114 | 53 |
| USP7_ICP0_bdg | -1.038950917 | 56 |
| LAG1-DNAbind | -1.037168777 | 56 |
| OB_Dis3 | -1.034375508 | 56 |
| Ceramidse_alk_C | -1.030852272 | 34 |
| PARP_reg | -1.028962138 | 38 |
| Gemin7 | -1.02764079 | 33 |
| G-alpha | -1.027265698 | 56 |
| Hid1 | -1.026203172 | 49 |
| PH_10 | -1.02593802 | 55 |
| TPR_7 | -1.024542503 | 56 |
| Ytp1 | -1.021388173 | 46 |
| Gelsolin | -1.018234508 | 56 |
| STPPase_N | -1.01557797 | 56 |
| ING | -1.014122881 | 53 |
| DEP | -1.013932071 | 55 |
| ANAPC1 | -1.013642949 | 35 |
| EF-hand_13 | -1.012914955 | 36 |
| Lon_C | -1.011490284 | 55 |
| RA | -1.011026775 | 56 |
| Na_Ca_ex | -1.010483908 | 56 |
| Ras_bdg_2 | -1.008934696 | 47 |
| PAS_11 | -1.00552841 | 36 |
| GRAM | -1.005103191 | 56 |
| PAP_RNA-bind | -1.004011926 | 54 |
| AAA_lid_2 | -0.999718035 | 23 |
| Ceramidase_alk | -0.997681739 | 34 |
| EHD_N | -0.995977024 | 50 |
| ELMO_CED12 | -0.995545846 | 46 |
| NMO | -0.992690057 | 48 |
| SAP | -0.991070233 | 54 |
| DUF202 | -0.990836758 | 55 |
| DnaJ-X | -0.989420115 | 55 |
| zf-CCCH_2 | -0.989034207 | 53 |
| Kinesin_assoc | -0.988778255 | 54 |
| Folliculin | -0.988482189 | 32 |
| PAS_3 | -0.987119172 | 46 |
| ANAPC3 | -0.987075796 | 51 |
| UBN_AB | -0.984103289 | 43 |
| Ecl1 | -0.983137317 | 54 |
| Asp_protease_2 | -0.98158392 | 38 |
| Sin3a_C | -0.980022523 | 55 |
| Aft1_HRA | -0.979054452 | 23 |
| CobW_C | -0.976267968 | 50 |
| BC10 | -0.972540289 | 26 |
| DUF2052 | -0.964999404 | 41 |
| Not1 | -0.963998273 | 55 |
| Ferritin | -0.96175637 | 45 |
| TENA_THI-4 | -0.961573962 | 42 |
| N1221 | -0.95779374 | 43 |
| PK_C | -0.955429158 | 54 |
| N6_Mtase | -0.95347415 | 45 |
| Galactosyl_T | -0.952200204 | 44 |
| DSPc | -0.949286408 | 56 |
| PH_9 | -0.949088727 | 56 |
| Glu-tRNAGln | -0.946950342 | 20 |
| Chromosome_seg | -0.94394038 | 42 |
| Cript | -0.943741875 | 40 |
| fn3_5 | -0.94366136 | 47 |
| TPR_11 | -0.942408577 | 56 |
| Patched | -0.941302088 | 51 |
| PNRC | -0.941019029 | 46 |
| Mitofilin | -0.940154828 | 55 |
| Nup35_RRM | -0.939710732 | 36 |
| End3 | -0.938585314 | 45 |
| Trehalose_PPase | -0.938276846 | 56 |
| Ribonuc_red_lgN | -0.933648239 | 56 |
| TYW3 | -0.932992764 | 37 |
| RSB_motif | -0.932143812 | 27 |
| CRM1_C | -0.932058608 | 55 |
| Ribosomal_L30_N | -0.931866207 | 55 |
| Rieske | -0.930065928 | 56 |
| Sin3_corepress | -0.928787576 | 56 |
| Amidohydro_3 | -0.926572403 | 52 |
| Biotin_lipoyl_2 | -0.926330604 | 26 |
| BET | -0.925345504 | 55 |
| CENP-K | -0.925011535 | 45 |
| TPR_14 | -0.919780377 | 31 |
| GPI-anchored | -0.918817938 | 55 |
| MOR2-PAG1_C | -0.917376275 | 47 |
| ATG2_CAD | -0.915107115 | 42 |
| PAS | -0.913379706 | 49 |
| CDC24 | -0.910678306 | 55 |
| Syntaxin-5_N | -0.908550506 | 37 |
| Ribosomal_S6e | -0.898740866 | 56 |
| TPR_17 | -0.898574397 | 49 |
| DUF2407 | -0.896533454 | 47 |
| Ribonuc_red_lgC | -0.895577912 | 56 |
| DUF3684 | -0.895017537 | 43 |
| FNIP_N | -0.894269318 | 32 |
| CENP-H | -0.893715409 | 39 |
| NTP_transf_3 | -0.893452878 | 56 |
| NPC1_N | -0.892256115 | 45 |
| DUF3402 | -0.889581458 | 51 |
| MULE | -0.885914098 | 24 |
| FAR1 | -0.884874249 | 45 |

| | | |
|---|---|---|
| FNIP_C | -0.884792872 | 38 |
| TPR_2 | -0.883851723 | 56 |
| Wyosine_form | -0.883029301 | 39 |
| NAD_Gly3P_dh_N | -0.880053806 | 56 |
| Ribosomal_L21e | -0.878801157 | 54 |
| Sds3 | -0.878634546 | 56 |
| NUDIX_4 | -0.878187343 | 37 |
| PK | -0.873388197 | 56 |
| CENP-I | -0.872373223 | 42 |
| Ubiquitin_4 | -0.871202563 | 33 |
| RhoGAP | -0.868021837 | 56 |
| Kelch_2 | -0.864918986 | 56 |
| DIL | -0.861626887 | 56 |
| RasGEF | -0.860406992 | 56 |
| Hexokinase_1 | -0.859478768 | 56 |
| Na_sulph_symp | -0.858664989 | 56 |
| PSP1 | -0.85763461 | 54 |
| PDEase_I | -0.854785225 | 56 |
| Methyltrn_RNA_3 | -0.854016894 | 43 |
| Disintegrin | -0.853857592 | 53 |
| DUF4050 | -0.849063533 | 40 |
| Vps54_N | -0.847937579 | 51 |
| Nup88 | -0.846192774 | 50 |
| Glyco_hydro_57 | -0.844970609 | 50 |
| CENP-M | -0.843999134 | 29 |
| TPR_1 | -0.843577663 | 56 |
| Ribosomal_L18 | -0.843476665 | 56 |
| Ubiquitin_2 | -0.840843843 | 55 |
| PDR_CDR | -0.839782392 | 45 |
| Arrestin_C | -0.838678661 | 53 |
| PH_12 | -0.836531103 | 35 |
| SWIRM | -0.833516457 | 56 |
| Ribosomal_L27e | -0.831311205 | 55 |
| TPR_6 | -0.831297953 | 54 |
| FNIP_M | -0.830671231 | 37 |
| Membr_traf_MHD | -0.829736845 | 48 |
| PH_11 | -0.828558886 | 36 |
| Tup_N | -0.82659214 | 54 |
| IF3_C | -0.824959746 | 38 |
| Pex24p | -0.822551979 | 55 |
| Ribosomal_L18A | -0.81707402 | 56 |
| Isochorismatase | -0.816289656 | 54 |
| FGGY_N | -0.814237859 | 56 |
| SCP2 | -0.813943638 | 56 |
| DNA_pol_A_exo1 | -0.810558239 | 55 |
| FGE-sulfatase | -0.809005378 | 49 |
| Fn3-like | -0.806338439 | 47 |
| NDC10_II | -0.804476861 | 21 |
| CNH | -0.80179628 | 56 |
| Hexokinase_2 | -0.799843473 | 56 |
| AAA_16 | -0.799483042 | 55 |
| Tmemb_185A | -0.798996389 | 49 |
| PPR_long | -0.798462675 | 55 |
| BTD | -0.795938452 | 56 |
| DUF3700 | -0.794555844 | 47 |
| Kelch_1 | -0.793680598 | 56 |
| muHD | -0.793066926 | 50 |
| PH_3 | -0.79268186 | 43 |
| STIMATE | -0.789411859 | 55 |
| DREV | -0.789003926 | 32 |
| PLDc_2 | -0.788794668 | 56 |
| Reprolysin_5 | -0.788116709 | 54 |
| Reprolysin_4 | -0.788116709 | 54 |
| Swi3 | -0.787860368 | 44 |
| Nuc_sug_transp | -0.787250486 | 55 |
| FtsH_ext | -0.785394481 | 48 |
| ACBP | -0.783502655 | 56 |
| CitMHS | -0.783473885 | 56 |
| Mtr2 | -0.783189337 | 37 |
| Ysc84 | -0.782065041 | 55 |
| bZIP_2 | -0.77730567 | 56 |
| ER | -0.776796654 | 41 |
| FDF | -0.774250377 | 50 |
| Mcp5_PH | -0.772764295 | 54 |
| UNC-50 | -0.772711353 | 38 |
| DDE_1 | -0.768419626 | 21 |
| Proteasom_Rpn13 | -0.767901669 | 50 |
| UDPG_MGDP_dh_C | -0.767775646 | 48 |
| RasGEF_N | -0.767383672 | 56 |
| Glyco_hydro_35 | -0.763708307 | 49 |
| Gp_dh_C | -0.763201816 | 55 |
| DUF2407_C | -0.761860031 | 46 |
| Atg29_N | -0.759954374 | 32 |
| ATP-cone | -0.759744854 | 56 |
| Whi5 | -0.759428063 | 25 |
| Fmp27_GFWDK | -0.759196392 | 42 |
| ILEI | -0.758742743 | 45 |
| Trehalase_Ca-bi | -0.758709969 | 55 |
| Ribosomal_L5 | -0.757673844 | 54 |
| Gp_dh_N | -0.756864118 | 55 |
| zf-MYND | -0.755716886 | 56 |
| DUF971 | -0.755435149 | 48 |
| RNB | -0.752787822 | 56 |
| RINGv | -0.752120665 | 53 |
| SWIRM-assoc_1 | -0.751076962 | 55 |
| CENP-X | -0.750605444 | 35 |
| CAP_GLY | -0.750314352 | 56 |
| SpoU_methylase | -0.749258788 | 53 |

| | | |
|---|---|---|
| Reprolysin_2 | -0.747354194 | 54 |
| DUF1746 | -0.746401414 | 35 |
| GST_C_5 | -0.744646597 | 35 |
| Arb1 | -0.744253065 | 49 |
| NAPRTase_N | -0.744244041 | 42 |
| GATA | -0.743929215 | 56 |
| TPR_19 | -0.743056577 | 56 |
| GBP | -0.742775139 | 51 |
| Pept_tRNA_hydro | -0.742371512 | 49 |
| CENP-N | -0.741766493 | 41 |
| Sterol-sensing | -0.741738737 | 52 |
| SPX | -0.740407651 | 56 |
| Gln-synt_C | -0.740310487 | 56 |
| Kelch_6 | -0.737538621 | 56 |
| Rad4 | -0.737333927 | 51 |
| UDPG_MGDP_dh | -0.735778883 | 48 |
| ATP-synt_ab_C | -0.734887425 | 55 |
| PAP2_3 | -0.734121926 | 46 |
| Sirohm_synth_C | -0.733626482 | 51 |
| Ribosomal_S25 | -0.733519345 | 54 |
| UPF0183 | -0.733350202 | 48 |
| DUF3819 | -0.733288793 | 55 |
| Ribosomal_L32e | -0.731074568 | 55 |
| TPT | -0.730504388 | 56 |
| PHM7_ext | -0.729901644 | 42 |
| Zds_C | -0.728555568 | 55 |
| TEA | -0.727075747 | 41 |
| Ribosomal_S26e | -0.72674061 | 56 |
| DUF3020 | -0.724516412 | 53 |
| Peptidase_C12 | -0.723858399 | 56 |
| DUF2405 | -0.722685907 | 41 |
| BAG | -0.722375225 | 54 |
| CDC50 | -0.721208029 | 56 |
| Fasciclin | -0.720731886 | 56 |
| CSN5_C | -0.716550257 | 45 |
| DUF1899 | -0.716539748 | 55 |
| DUF3474 | -0.714534307 | 37 |
| PH | -0.713660162 | 56 |
| Reprolysin_3 | -0.710676681 | 54 |
| PAS_9 | -0.70597304 | 48 |
| Frag1 | -0.705433401 | 50 |
| Ribosomal_60s | -0.705049988 | 56 |
| Actin_micro | -0.704640882 | 49 |
| EF-hand_11 | -0.702347777 | 23 |
| Ribosomal_S7e | -0.702335986 | 56 |
| MUG113 | -0.702265199 | 34 |
| MIT | -0.701855984 | 54 |
| NIR_SIR_ferr | -0.699833821 | 50 |
| PUF | -0.699733659 | 56 |
| PXA | -0.699102854 | 48 |
| CLIP1_ZNF | -0.698674847 | 41 |
| Ank | -0.696900679 | 56 |
| CTD_bind | -0.695901822 | 48 |
| SRF-TF | -0.69371886 | 56 |
| Arm_2 | -0.693040448 | 47 |
| FGGY_C | -0.691743408 | 56 |
| SH3_2 | -0.691428322 | 56 |
| CNOT1_TTP_bind | -0.690093995 | 55 |
| PHD | -0.68887188 | 56 |
| Peroxidase_2 | -0.688471407 | 29 |
| MAS20 | -0.688184664 | 56 |
| Septin | -0.687871434 | 56 |
| zf-CCHC | -0.687521668 | 56 |
| HSA | -0.687169992 | 56 |
| Ank_5 | -0.686495085 | 56 |
| HAMP | -0.686212073 | 53 |
| TMEM135_C_rich | -0.684305428 | 45 |
| CTP_transf_1 | -0.682040079 | 56 |
| PHD_2 | -0.681431432 | 53 |
| Methyltransf_32 | -0.681250368 | 53 |
| Rap_GAP | -0.67835764 | 56 |
| KA1 | -0.676156616 | 54 |
| RIH_assoc | -0.675880127 | 21 |
| Rieske_2 | -0.67581665 | 26 |
| CNOT1_HEAT | -0.675815238 | 54 |
| MgtE | -0.675310905 | 49 |
| HR1 | -0.674970669 | 41 |
| C1_1 | -0.674324219 | 56 |
| SRAP | -0.672213898 | 42 |
| WW | -0.672067441 | 56 |
| UBM | -0.67067636 | 51 |
| Ribosomal_S24e | -0.670658555 | 56 |
| Vps5 | -0.670337124 | 55 |
| Ribosomal_L37ae | -0.665845847 | 55 |
| UDPG_MGDP_dh_N | -0.665277906 | 49 |
| AA_permease_2 | -0.665254682 | 56 |
| Ribosomal_L26 | -0.663717052 | 56 |
| GGACT | -0.66306516 | 41 |
| GFO_IDH_MocA | -0.662794123 | 50 |
| FYRN | -0.662448899 | 56 |
| TPR_8 | -0.662206962 | 56 |
| Sterol_MT_C | -0.661670873 | 53 |
| NAD_binding_7 | -0.660149255 | 51 |
| NAD_Gly3P_dh_C | -0.659966149 | 56 |
| Clathrin_propel | -0.659820047 | 52 |
| NPL | -0.6586672 | 54 |
| Ribosomal_L28e | -0.655204125 | 55 |
| Acyltransferase | -0.653594588 | 56 |

| | | |
|---|---|---|
| XPG_I_2 | -0.652756419 | 47 |
| RecA | -0.651015134 | 43 |
| Ribosomal_S3Ae | -0.651007919 | 55 |
| Ribosomal_L23eN | -0.648266574 | 54 |
| HEAT | -0.647528706 | 56 |
| FCH | -0.646025726 | 56 |
| MATH | -0.645383809 | 53 |
| HAGH_C | -0.644995461 | 54 |
| Pirin_C | -0.644736925 | 52 |
| Glyco_transf_20 | -0.644220697 | 56 |
| SH3_9 | -0.643960643 | 56 |
| zf-A20 | -0.643875147 | 42 |
| zf_UBZ | -0.643623022 | 26 |
| Ribosomal_L39 | -0.643465694 | 54 |
| N_BRCA1_IG | -0.639757467 | 46 |
| Pkinase_Tyr | -0.639294822 | 56 |
| NUDE_C | -0.634823573 | 42 |
| Glyco_hydro_42 | -0.633300666 | 41 |
| SWIRM-assoc_2 | -0.630769544 | 43 |
| SPRY | -0.630426794 | 55 |
| Arm_3 | -0.630344443 | 56 |
| PX | -0.626899247 | 56 |
| Ribosomal_L34e | -0.625373088 | 55 |
| RFX_DNA_binding | -0.624223055 | 26 |
| GRASP55_65 | -0.623494032 | 51 |
| EMP70 | -0.622573252 | 56 |
| PDCD7 | -0.62215102 | 28 |
| IBB | -0.620120388 | 56 |
| DAO | -0.619194823 | 56 |
| Peptidase_S15 | -0.61614545 | 47 |
| PPR_3 | -0.615526836 | 56 |
| Ribosomal_L44 | -0.61498585 | 55 |
| DPBB_1 | -0.614462643 | 54 |
| TRM13 | -0.613399782 | 49 |
| PB1 | -0.612836817 | 56 |
| Ribosomal_S19e | -0.612757941 | 54 |
| Fmp27_WPPW | -0.611827185 | 42 |
| DUF2358 | -0.611757294 | 33 |
| POT1PC | -0.609814651 | 41 |
| Spc7 | -0.608603694 | 42 |
| Usp | -0.607755261 | 56 |
| zf-U11-48K | -0.607589718 | 46 |
| Arginase | -0.604095489 | 54 |
| NIR_SIR | -0.602303724 | 50 |
| CENP-L | -0.602290564 | 39 |
| Ion_trans | -0.602261608 | 56 |
| G6PD_N | -0.598273245 | 55 |
| CNOT1_CAF1_bind | -0.597590532 | 55 |
| TPR_16 | -0.596081063 | 56 |
| VPS13 | -0.595400472 | 54 |
| VHS | -0.594818745 | 56 |
| IQ | -0.594642151 | 56 |
| SH3_1 | -0.594467649 | 56 |
| Cullin_binding | -0.593515279 | 54 |
| RPAP3_C | -0.593203463 | 43 |
| DAGK_cat | -0.591003781 | 54 |
| SAM_2 | -0.589774937 | 56 |
| DAGK_acc | -0.586452583 | 34 |
| PBD | -0.585553963 | 56 |
| RsgA_GTPase | -0.584743368 | 22 |
| DUF5427 | -0.584220547 | 47 |
| Ribosomal_S3_C | -0.581748856 | 55 |
| Velvet | -0.580483878 | 56 |
| LDB19 | -0.579578409 | 38 |
| AWS | -0.578451986 | 55 |
| Bap31 | -0.578342348 | 55 |
| VPS13_C | -0.577025402 | 54 |
| DNA_ligase_A_N | -0.574153706 | 55 |
| PI31_Prot_N | -0.572216211 | 46 |
| Lgl_C | -0.567377593 | 38 |
| Ribosomal_S28e | -0.566556814 | 55 |
| S36_mt | -0.566395029 | 39 |
| GMC_oxred_C | -0.566286403 | 46 |
| DIOX_N | -0.565541817 | 54 |
| T5orf172 | -0.565492703 | 38 |
| Pkinase | -0.564709759 | 56 |
| SRA1 | -0.564238397 | 39 |
| C2 | -0.56283569 | 56 |
| SAM_1 | -0.562381561 | 56 |
| NCE101 | -0.56216567 | 28 |
| EcKinase | -0.561063154 | 43 |
| ZZ | -0.559745301 | 56 |
| Nicastrin | -0.558804321 | 41 |
| FYRC | -0.558604923 | 56 |
| LIM | -0.558125476 | 56 |
| NAD_binding_10 | -0.556571371 | 56 |
| Abhydro_lipase | -0.555702634 | 52 |
| NAD_binding_8 | -0.554637383 | 56 |
| Chromo_shadow | -0.554198376 | 53 |
| TRP | -0.550454709 | 45 |
| GIT_SHD | -0.546122472 | 53 |
| G6PD_C | -0.545132435 | 55 |
| Sirohm_synth_M | -0.544584491 | 51 |
| Ribosomal_L6e | -0.542845544 | 56 |
| PBP | -0.542225331 | 55 |
| Peptidase_C39 | -0.541732092 | 23 |
| PA | -0.537843577 | 56 |
| Hydrolase_6 | -0.537174161 | 56 |

| | | |
|---|---|---|
| S10_plectin | -0.536168902 | 56 |
| malic | -0.535674327 | 55 |
| Ank_4 | -0.534783365 | 56 |
| SnAC | -0.533624218 | 55 |
| CBM_21 | -0.533224487 | 54 |
| PLU-1 | -0.532026616 | 41 |
| PIP5K | -0.531612013 | 56 |
| WAC_Acf1_DNA_bd | -0.530877272 | 53 |
| Rib_5-P_isom_A | -0.530791648 | 50 |
| Ribosomal_L35Ae | -0.52782841 | 55 |
| CRT-like | -0.525604223 | 46 |
| Ribosomal_L38e | -0.523023601 | 54 |
| Forkhead | -0.521591587 | 56 |
| gag-asp_proteas | -0.520222013 | 34 |
| Ribosomal_S27e | -0.519609707 | 55 |
| AAA_30 | -0.518720331 | 56 |
| Rax2 | -0.517170841 | 55 |
| Mg_trans_NIPA | -0.516750367 | 54 |
| Rad60-SLD | -0.51515487 | 56 |
| DUF3336 | -0.514634571 | 52 |
| Malic_M | -0.514537324 | 55 |
| Methyltransf_23 | -0.51378639 | 56 |
| Ribosomal_S8e | -0.513690935 | 56 |
| tRNA_int_endo_N | -0.511958362 | 20 |
| MFS_2 | -0.511094468 | 56 |
| EF1_GNE | -0.5093042 | 48 |
| Pkinase_C | -0.509199039 | 56 |
| VPS9 | -0.508793922 | 56 |
| MIF | -0.50873895 | 34 |
| Abhydrolase_2 | -0.50657893 | 55 |
| Acetyltransf_3 | -0.505737479 | 55 |
| ERCC3_RAD25_C | -0.504715891 | 56 |
| DUF155 | -0.501680678 | 56 |
| Bac_surface_Ag | -0.500997283 | 55 |
| CSD | -0.500804669 | 39 |
| KH_2 | -0.498734477 | 55 |
| PPR_2 | -0.495940618 | 56 |
| Acetyltransf_1 | -0.495551431 | 56 |
| Ribosomal_L37e | -0.495459519 | 55 |
| Ank_3 | -0.494659885 | 56 |
| DNA_ligase_A_C | -0.494647519 | 55 |
| DUF2043 | -0.494363734 | 38 |
| Bac_GDH | -0.492939908 | 55 |
| NmrA | -0.492593036 | 56 |
| AdoHcyase_NAD | -0.490213809 | 55 |
| ATG13 | -0.489431508 | 50 |
| Peroxin-13_N | -0.48913324 | 55 |
| Pyrophosphatase | -0.487547112 | 56 |
| Ribonucleas_3_3 | -0.486696407 | 55 |
| THUMP | -0.486655261 | 54 |
| RabGAP-TBC | -0.486622084 | 56 |
| CENP-O | -0.486600472 | 41 |
| Opi1 | -0.485774766 | 54 |
| DUF908 | -0.48561351 | 53 |
| cNMP_binding | -0.483576953 | 56 |
| Ribosomal_L5_C | -0.481579196 | 56 |
| ArfGap | -0.48096275 | 56 |
| BAR | -0.477805748 | 56 |
| Ribosomal_L29e | -0.47708034 | 53 |
| DUF3419 | -0.476082571 | 42 |
| VPS13_mid_rpt | -0.475788051 | 54 |
| FoP_duplication | -0.475386323 | 53 |
| zf-RanBP | -0.474222431 | 53 |
| Pyr_redox_3 | -0.472266327 | 56 |
| DUF4210 | -0.471595113 | 43 |
| Bin3 | -0.468165106 | 41 |
| DUF4042 | -0.467370014 | 48 |
| RhoGEF | -0.466757136 | 56 |
| Pep_M12B_propep | -0.465902857 | 33 |
| Ribosomal_L19e | -0.464852942 | 55 |
| DUF913 | -0.464821344 | 49 |
| Ribosomal_L36e | -0.464741962 | 56 |
| Nucleopor_Nup85 | -0.463835753 | 51 |
| Med13_C | -0.462910207 | 50 |
| HATPase_c | -0.461686052 | 56 |
| B-block_TFIIIC | -0.460619821 | 49 |
| DLIC | -0.45984843 | 56 |
| Nexin_C | -0.459841618 | 48 |
| Itfg2 | -0.459361012 | 47 |
| Polysacc_synt_2 | -0.45929592 | 56 |
| CoA_transf_3 | -0.458619435 | 55 |
| Say1_Mug180 | -0.458541355 | 55 |
| SNARE_assoc | -0.457449988 | 55 |
| GDP_Man_Dehyd | -0.453608835 | 56 |
| Pyrid_oxidase_2 | -0.453433351 | 37 |
| AA_permease | -0.4531186 | 56 |
| Cyt-b5 | -0.45230501 | 56 |
| DUF2408 | -0.451026556 | 54 |
| EF-1_beta_acid | -0.450397169 | 48 |
| EF-hand_4 | -0.450058095 | 56 |
| HIRAN | -0.449633809 | 44 |
| SRI | -0.448565079 | 45 |
| UPF0176_N | -0.447191009 | 54 |
| CTP_transf_like | -0.446711552 | 56 |
| FG-GAP | -0.446675966 | 50 |
| MMS19_N | -0.443355834 | 54 |
| Nt_Gln_amidase | -0.441913241 | 36 |
| DUF2075 | -0.441708969 | 39 |

| | | |
|---|---|---|
| Ribosomal_S21e | -0.439805912 | 56 |
| Methyltransf_31 | -0.439798729 | 56 |
| BRCT_2 | -0.43928619 | 55 |
| Arrestin_N | -0.439095283 | 55 |
| Sec3-PIP2_bind | -0.436851539 | 42 |
| Beach | -0.436609645 | 54 |
| DUF2427 | -0.435386085 | 46 |
| Methyltransf_25 | -0.43405609 | 56 |
| Rhodanese_C | -0.433971933 | 53 |
| Phytochelatin | -0.432084684 | 50 |
| SPA | -0.431531153 | 56 |
| KCH | -0.431111994 | 46 |
| Ribosomal_S2 | -0.431068372 | 56 |
| Ribosomal_S19 | -0.430329751 | 56 |
| SM-ATX | -0.429474515 | 54 |
| TBCC_N | -0.429368656 | 44 |
| RimK | -0.428685208 | 51 |
| FR47 | -0.428563895 | 56 |
| MMS19_C | -0.427559277 | 54 |
| Peptidase_C65 | -0.427471874 | 48 |
| Med15_fungi | -0.424667689 | 44 |
| Aldo_ket_red | -0.424346464 | 56 |
| Nramp | -0.424254424 | 53 |
| CRM1_repeat_2 | -0.42322145 | 55 |
| zf-C2H2 | -0.423126759 | 56 |
| IATP | -0.423032224 | 51 |
| peroxidase | -0.422863517 | 54 |
| PC4 | -0.422408959 | 41 |
| LIM_bind | -0.422277768 | 48 |
| Ribosomal_L27A | -0.419873841 | 56 |
| ubiquitin | -0.419707247 | 56 |
| 2-Hacid_dh | -0.418488824 | 56 |
| Fungal_trans | -0.417616533 | 56 |
| Chorein_N | -0.417001834 | 55 |
| Ribosomal_S30 | -0.416264843 | 53 |
| Ribosomal_L31e | -0.414733354 | 55 |
| Thioredoxin_9 | -0.414450617 | 23 |
| Peptidase_A22B | -0.413222366 | 54 |
| ALG3 | -0.412333274 | 49 |
| HMG_box_2 | -0.412276678 | 56 |
| Sec23_BS | -0.411376704 | 56 |
| Abhydrolase_3 | -0.410697816 | 56 |
| HEAT_2 | -0.410407448 | 56 |
| MFS_5 | -0.409972631 | 43 |
| Peptidase_C78 | -0.409035803 | 43 |
| Sec23_helical | -0.408951701 | 56 |
| Methyltransf_11 | -0.408708978 | 56 |
| HSP70 | -0.408612653 | 56 |
| ZT_dimer | -0.407541462 | 55 |
| AAA_19 | -0.405463867 | 56 |
| Glyco_trans_2_3 | -0.405318072 | 56 |
| Pho88 | -0.404861424 | 55 |
| zf-RING-like | -0.400207068 | 45 |
| PPIP5K2_N | -0.399826333 | 55 |
| 2OG-FeII_Oxy | -0.39902294 | 53 |
| RHD3 | -0.397437772 | 55 |
| Ribosomal_L30 | -0.396916924 | 56 |
| MARVEL | -0.3967595 | 43 |
| Chitin_synth_2 | -0.396229216 | 56 |
| Motile_Sperm | -0.395687959 | 56 |
| UPF0220 | -0.388456064 | 54 |
| LRR_8 | -0.38752155 | 56 |
| Cdh1_DBD_1 | -0.387077187 | 30 |
| Ebp2 | -0.38694798 | 55 |
| FANCI_HD2 | -0.386710303 | 47 |
| Arm | -0.38637218 | 56 |
| CSD2 | -0.384614268 | 43 |
| G-gamma | -0.383949558 | 55 |
| DUF1681 | -0.383872552 | 45 |
| Glyco_hydro_3_C | -0.383129031 | 48 |
| Mon2_C | -0.382871083 | 54 |
| SprT-like | -0.382662702 | 36 |
| EPL1 | -0.381619305 | 55 |
| Kdo | -0.38136284 | 56 |
| Glycogen_syn | -0.381301266 | 54 |
| HTH_3 | -0.381207954 | 54 |
| Transketolase_N | -0.381152085 | 55 |
| Y_phosphatase | -0.380428397 | 56 |
| Y_phosphatase3 | -0.380357974 | 56 |
| NAD_binding_2 | -0.380315622 | 56 |
| F420_oxidored | -0.380113261 | 56 |
| tRNA-synt_1d | -0.379936439 | 54 |
| ANTH | -0.378795478 | 56 |
| Acetyltransf_7 | -0.378637621 | 56 |
| PFK | -0.378437181 | 55 |
| MMtag | -0.377319956 | 44 |
| CUE | -0.376630544 | 56 |
| Acyl-CoA_ox_N | -0.376373885 | 56 |
| Sec16 | -0.37579982 | 47 |
| Glyco_hydro_15 | -0.374900875 | 56 |
| FKS1_dom1 | -0.374653375 | 54 |
| SurE | -0.374582407 | 24 |
| Methyltransf_16 | -0.374479226 | 56 |
| Ribosomal_L2_C | -0.373197012 | 56 |
| UPF0020 | -0.371710191 | 56 |
| Thioredoxin_7 | -0.371025729 | 51 |
| Cyclin | -0.370952306 | 56 |
| CRM1_repeat_3 | -0.370751174 | 55 |

| | | |
|---|---|---|
| MyTH4 | -0.369591596 | 52 |
| UAA | -0.368513466 | 56 |
| Ribosomal_L13e | -0.367664563 | 55 |
| zf-C3H1 | -0.366405625 | 31 |
| Myb_DNA-bind_6 | -0.364302945 | 56 |
| Creatinase_N | -0.364187278 | 55 |
| Tetraspanin | -0.362568007 | 55 |
| Pirin | -0.362464647 | 52 |
| PPR | -0.360949871 | 56 |
| CRM1_repeat | -0.360704598 | 55 |
| His_Phos_1 | -0.360498795 | 56 |
| ACOX | -0.360313113 | 56 |
| DUF647 | -0.359295875 | 50 |
| Cupin_2 | -0.359186685 | 54 |
| HEAT_EZ | -0.358036082 | 56 |
| EF-hand_7 | -0.35793016 | 56 |
| EF-hand_1 | -0.357537298 | 56 |
| Glucan_synthase | -0.355485428 | 54 |
| HHH_5 | -0.355243751 | 47 |
| ELL | -0.354879669 | 24 |
| Bromodomain | -0.353984664 | 56 |
| RTA1 | -0.35358199 | 33 |
| zf-RING_4 | -0.353559139 | 53 |
| NTP_transf_2 | -0.353484837 | 56 |
| WH1 | -0.353309441 | 55 |
| DAHP_synth_1 | -0.353092898 | 56 |
| Mit_ribos_Mrp51 | -0.352858935 | 54 |
| BSD | -0.352682819 | 54 |
| HMG-CoA_red | -0.351824329 | 54 |
| Ribosomal_L14e | -0.350042451 | 56 |
| Peptidase_C1_2 | -0.350041338 | 50 |
| Clathrin-link | -0.348650138 | 51 |
| Ank_2 | -0.348538356 | 56 |
| Glu_synthase | -0.344382796 | 56 |
| BHD_2 | -0.344192443 | 46 |
| SHR-BD | -0.342011353 | 55 |
| IMS_C | -0.341930408 | 51 |
| Arf | -0.341855587 | 56 |
| EFhand_Ca_insen | -0.34171493 | 55 |
| zf-Sec23_Sec24 | -0.339449489 | 56 |
| Pex16 | -0.339314385 | 54 |
| EB1 | -0.33794176 | 53 |
| HK | -0.337789333 | 41 |
| ClpS | -0.337628795 | 56 |
| Ribosomal_S17e | -0.33717851 | 56 |
| Hydrolase_3 | -0.335487037 | 56 |
| LSM14 | -0.335119901 | 51 |
| NUDIX | -0.334900276 | 56 |
| Dynactin | -0.334821865 | 47 |
| V-ATPase_H_N | -0.334239193 | 56 |
| SEN1_N | -0.333969124 | 37 |
| Kinesin | -0.333764015 | 56 |
| ACT_6 | -0.332428583 | 40 |
| Alpha_adaptinC2 | -0.33235413 | 56 |
| DUF1749 | -0.331537207 | 35 |
| PH_8 | -0.330649388 | 50 |
| MR_MLE_C | -0.330624984 | 47 |
| zf-C2H2_6 | -0.327315766 | 56 |
| Not3 | -0.326877453 | 55 |
| DUF2183 | -0.326743448 | 53 |
| CK_II_beta | -0.326194491 | 56 |
| Ammonium_transp | -0.325605146 | 56 |
| UPF0121 | -0.325366802 | 56 |
| Acetyltransf_10 | -0.322550003 | 56 |
| Methyltransf_12 | -0.322538903 | 56 |
| HMG_box | -0.322403756 | 56 |
| Bap31_Bap29_C | -0.32237939 | 45 |
| PS_Dcarbxylase | -0.322128841 | 56 |
| AAA_12 | -0.320898662 | 56 |
| Ribosomal_L2 | -0.320783184 | 56 |
| RecQ_Zn_bind | -0.318710688 | 53 |
| PAZ | -0.318192069 | 55 |
| ATP-synt_ab | -0.31796401 | 56 |
| C12orf66_like | -0.317695025 | 43 |
| Fer4_4 | -0.316866037 | 48 |
| EMG1 | -0.316772184 | 56 |
| AIP3 | -0.316714786 | 56 |
| Sec23_trunk | -0.315818286 | 56 |
| GXGXG | -0.315604984 | 53 |
| ADH_N_2 | -0.315415906 | 51 |
| Syntaxin | -0.313568707 | 56 |
| MS_channel | -0.310222697 | 56 |
| Rad60-SLD_2 | -0.310065089 | 53 |
| Mo-co_dimer | -0.309893141 | 53 |
| Mitochondr_Som1 | -0.308362513 | 36 |
| RNase_H2-Ydr279 | -0.306566052 | 40 |
| RasGAP_C | -0.305624418 | 56 |
| Peptidase_M41 | -0.305172768 | 56 |
| FHA | -0.305114413 | 56 |
| Glu_syn_central | -0.304940769 | 53 |
| Aa_trans | -0.304894027 | 56 |
| Meth_synt_2 | -0.303927743 | 55 |
| UMPH-1 | -0.303858045 | 40 |
| CDT1 | -0.303699035 | 39 |
| RraA-like | -0.302400013 | 49 |
| CBS | -0.302372199 | 56 |
| Cons_hypoth95 | -0.301251024 | 45 |
| AAA_lid_1 | -0.300468011 | 33 |

| | | |
|---|---|---|
| IF4E | -0.300402861 | 56 |
| Ribosomal_L23 | -0.299219405 | 56 |
| Laminin_G_3 | -0.299067715 | 33 |
| TniB | -0.298658428 | 43 |
| AZUL | -0.2974528 | 47 |
| Med23 | -0.296627557 | 40 |
| eIF3m_C_helix | -0.295892076 | 48 |
| Fibrillarin | -0.294246596 | 55 |
| La | -0.293496512 | 56 |
| Helicase_PWI | -0.293331714 | 52 |
| Oxidored_molyb | -0.292374105 | 54 |
| Nup54 | -0.29223764 | 53 |
| ATP-synt_ab_N | -0.287668825 | 56 |
| Ribosomal_S9 | -0.28616401 | 56 |
| Aminotran_3 | -0.285505324 | 56 |
| Glyco_transf_8 | -0.285257409 | 56 |
| GPP34 | -0.284489807 | 52 |
| AraC_binding | -0.283845232 | 50 |
| IlvC | -0.283681976 | 55 |
| INTS2 | -0.281446707 | 48 |
| DUF5601 | -0.281350398 | 52 |
| CBFB_NFYA | -0.28115934 | 56 |
| SQS_PSY | -0.280271728 | 56 |
| EXS | -0.278103342 | 56 |
| ER_lumen_recept | -0.277115754 | 56 |
| EF-hand_5 | -0.276945516 | 56 |
| RPAP1_N | -0.276694428 | 45 |
| STE | -0.275893326 | 56 |
| Fer4_20 | -0.274721781 | 53 |
| Vps35 | -0.273314512 | 55 |
| Gtr1_RagA | -0.272800097 | 56 |
| Glyco_trans_1_4 | -0.271163196 | 56 |
| Glyco_hydro_31 | -0.269994046 | 56 |
| Rrp44_CSD1 | -0.269237049 | 55 |
| TFIIS_M | -0.26906949 | 55 |
| P5CR_dimer | -0.267727659 | 56 |
| COX7a | -0.266069869 | 51 |
| 6PF2K | -0.265890817 | 56 |
| 4HBT_2 | -0.265864651 | 38 |
| SecY | -0.265492271 | 56 |
| DNA_pol_B | -0.264981543 | 56 |
| AAA_25 | -0.264884007 | 54 |
| Ribosomal_S5 | -0.264862516 | 56 |
| U3snoRNP10 | -0.264628864 | 43 |
| EamA | -0.264591414 | 56 |
| zf-HC5HC2H | -0.264523495 | 55 |
| WSD | -0.262442428 | 55 |
| Ribosomal_L22 | -0.262000603 | 56 |
| SET | -0.260593845 | 56 |
| Topoisom_bac | -0.260263765 | 55 |
| WD40 | -0.25993307 | 56 |
| RNA12 | -0.259676921 | 52 |
| Oxysterol_BP | -0.258429806 | 56 |
| Pan3_PK | -0.258181895 | 51 |
| Cupin_8 | -0.257482198 | 56 |
| GLTP | -0.255497381 | 53 |
| DUF2370 | -0.255164893 | 56 |
| CRC_subunit | -0.254729927 | 55 |
| 40S_S4_C | -0.253865158 | 56 |
| ADH_N | -0.253609121 | 56 |
| HSP90 | -0.253564033 | 55 |
| ECH_2 | -0.251537713 | 56 |
| UDPGP | -0.25087194 | 56 |
| SWIB | -0.25085411 | 56 |
| RS4NT | -0.250653077 | 55 |
| Sec39 | -0.250047594 | 49 |
| Lactamase_B | -0.248501846 | 56 |
| Microtub_bd | -0.248451311 | 56 |
| Nup160 | -0.248235735 | 51 |
| PARP | -0.246783195 | 43 |
| Ribosomal_L24e | -0.24661091 | 55 |
| DUF676 | -0.24659224 | 52 |
| Ada3 | -0.24608698 | 55 |
| Kinase-like | -0.24370054 | 56 |
| CAF1C_H4-bd | -0.242953264 | 56 |
| Guanylate_cyc | -0.24197658 | 56 |
| HATPase_c_3 | -0.241614738 | 56 |
| RCC1 | -0.241333935 | 56 |
| Rhomboid | -0.241208313 | 56 |
| AA_kinase | -0.241200489 | 56 |
| CH | -0.240917648 | 56 |
| Ribosomal_S4e | -0.240573444 | 56 |
| zf-HC5HC2H_2 | -0.240390925 | 55 |
| Ribosomal_L18_c | -0.240120709 | 56 |
| Dynamin_N | -0.239994685 | 56 |
| LisH_TPL | -0.239905853 | 53 |
| TBP-binding | -0.239822816 | 35 |
| GalP_UDP_transf | -0.238976642 | 51 |
| DUF1977 | -0.237859352 | 54 |
| Yop-YscD_cpl | -0.23782805 | 25 |
| ALIX_LYPXL_bnd | -0.236006262 | 56 |
| MMR_HSR1 | -0.235931756 | 56 |
| Myosin_N | -0.234791593 | 44 |
| Ribosomal_L5e | -0.23392339 | 56 |
| W2 | -0.233888313 | 56 |
| V-ATPase_H_C | -0.233576525 | 56 |
| Phos_pyr_kin | -0.232009566 | 54 |
| Malate_synthase | -0.231885695 | 56 |

| | | |
|---|---|---|
| Ribosomal_L11_N | -0.231830949 | 56 |
| Xpo1 | -0.231375864 | 56 |
| Glucosamine_iso | -0.227531488 | 56 |
| Shikimate_DH | -0.227527373 | 56 |
| Fer4_8 | -0.226670781 | 48 |
| K_oxygenase | -0.226552099 | 54 |
| TFIID_20kDa | -0.22609583 | 55 |
| FAD-oxidase_C | -0.225896509 | 56 |
| ArgoL2 | -0.225886481 | 54 |
| Myosin_head | -0.225882817 | 56 |
| Glyco_trans_1_2 | -0.225867248 | 31 |
| Glyco_tranf_2_3 | -0.22570013 | 56 |
| FAD_binding_2 | -0.225396753 | 56 |
| Rxt3 | -0.224935702 | 37 |
| UCH_1 | -0.223813514 | 56 |
| MBA1 | -0.223786545 | 47 |
| Roc | -0.222844729 | 56 |
| GCFC | -0.221914008 | 55 |
| UvrD-helicase | -0.221765295 | 50 |
| Mt_ATP-synt_B | -0.221688998 | 55 |
| Cg6151-P | -0.220773418 | 53 |
| Na_H_Exchanger | -0.220616566 | 55 |
| EF1G | -0.220562144 | 48 |
| WD40_4 | -0.219871146 | 55 |
| Complex1_LYR_1 | -0.21957791 | 24 |
| WGR | -0.219520504 | 38 |
| EF-hand_8 | -0.218260007 | 56 |
| Uds1 | -0.218181563 | 47 |
| Spt20 | -0.218015414 | 43 |
| Enolase_C | -0.217775408 | 56 |
| Flavodoxin_1 | -0.217160797 | 56 |
| DUF1640 | -0.217068939 | 55 |
| Myb_DNA-binding | -0.217060619 | 56 |
| CTP_synth_N | -0.216172078 | 55 |
| VWA | -0.215166502 | 26 |
| Complex1_LYR_2 | -0.214128772 | 56 |
| DUF1295 | -0.213259732 | 53 |
| IlvN | -0.212553995 | 56 |
| UCR_TM | -0.212308865 | 55 |
| Asp_protease | -0.212149142 | 33 |
| zf-GRF | -0.210601052 | 50 |
| 40S_SA_C | -0.210460661 | 42 |
| Enolase_N | -0.209051785 | 56 |
| DUF4604 | -0.208518052 | 46 |
| 6PGD | -0.207882324 | 56 |
| Ribosomal_L11 | -0.207726315 | 56 |
| Ribosomal_L7Ae | -0.207426803 | 56 |
| Vps39_1 | -0.206896138 | 53 |
| Oxidored_FMN | -0.206572262 | 54 |
| CBM53 | -0.205933829 | 52 |
| Lactamase_B_3 | -0.20551086 | 53 |
| Ribosomal_L13 | -0.205109291 | 56 |
| YhhN | -0.204211516 | 22 |
| FAD_binding_1 | -0.20409635 | 56 |
| DeoC | -0.203983469 | 36 |
| DUF4187 | -0.203172936 | 52 |
| Plug_translocon | -0.203060244 | 56 |
| DASH_Dad2 | -0.203038853 | 42 |
| YchF-GTPase_C | -0.201998455 | 56 |
| FumaraseC_C | -0.201896254 | 56 |
| RNase_T | -0.201525952 | 56 |
| Ras | -0.200708091 | 56 |
| DALR_1 | -0.200423475 | 54 |
| HMA | -0.200345316 | 56 |
| DUF2009 | -0.200249033 | 51 |
| IBR | -0.198963516 | 56 |
| zf-UBR | -0.198315609 | 56 |
| Thioredoxin_15 | -0.198309827 | 47 |
| QLQ | -0.197712768 | 52 |
| DUF727 | -0.196636063 | 52 |
| Vps51 | -0.196319074 | 56 |
| RAI16-like | -0.196313992 | 56 |
| RNase_P_p30 | -0.196214999 | 56 |
| ERG4_ERG24 | -0.195570273 | 55 |
| GFO_IDH_MocA_C | -0.195324155 | 46 |
| PH_TFIIH | -0.195118416 | 44 |
| Bromo_TP | -0.194938453 | 56 |
| UIM | -0.194542422 | 53 |
| Acyl-CoA_dh_M | -0.194307406 | 56 |
| NTP_transferase | -0.194214457 | 56 |
| Histone_H2A_C | -0.191996082 | 56 |
| THF_DHG_CYH_C | -0.191138868 | 56 |
| Ecm29 | -0.19089519 | 49 |
| Amino_oxidase | -0.190646194 | 56 |
| FA_hydroxylase | -0.188007512 | 56 |
| ArgoL1 | -0.187779606 | 54 |
| BRCA-2_helical | -0.187105231 | 36 |
| Raffinose_syn | -0.186372172 | 25 |
| DNA_pol_B_exo1 | -0.186064223 | 56 |
| COX6A | -0.185754834 | 56 |
| Ribosomal_S5_C | -0.185429269 | 56 |
| Sugar_tr | -0.183519964 | 56 |
| BRO1 | -0.183141219 | 56 |
| CPT | -0.182970462 | 42 |
| NAD_binding_3 | -0.182286991 | 55 |
| LisH | -0.181583852 | 55 |
| 4HBT_3 | -0.181449741 | 46 |
| UCH_C | -0.181105389 | 51 |

| | | |
|---|---|---|
| zf-MIZ | -0.180645955 | 54 |
| TTL | -0.179627158 | 46 |
| Cir_N | -0.178863105 | 56 |
| PEX11 | -0.178638356 | 56 |
| MFS_1 | -0.177469751 | 56 |
| CLTH | -0.175420436 | 56 |
| Dor1 | -0.174927924 | 52 |
| Glycos_transf_1 | -0.172631991 | 56 |
| Glycos_transf_2 | -0.17111389 | 56 |
| Carboxyl_trans | -0.171055455 | 56 |
| DNA_mis_repair | -0.170526518 | 55 |
| ENTH | -0.169433554 | 56 |
| MBF1 | -0.169361679 | 54 |
| SURF4 | -0.169345612 | 56 |
| DUF1998 | -0.168934846 | 24 |
| Noc2 | -0.168472756 | 54 |
| RAC_head | -0.168345002 | 46 |
| Pex14_N | -0.167721434 | 56 |
| THF_DHG_CYH | -0.167516228 | 56 |
| 2-Hacid_dh_C | -0.167008311 | 56 |
| UreF | -0.166544836 | 48 |
| DUF21 | -0.165110785 | 56 |
| FYVE | -0.164459864 | 56 |
| ADH_zinc_N_2 | -0.163934779 | 56 |
| ADH_zinc_N | -0.162507241 | 56 |
| Glyco_hydro_20b | -0.162472648 | 45 |
| AAA_11 | -0.16223761 | 55 |
| Yae1_N | -0.161820979 | 51 |
| zf-CCHC_6 | -0.161365927 | 51 |
| zf-C6H2 | -0.160503484 | 55 |
| UPF0014 | -0.160369151 | 53 |
| AMP_N | -0.159337772 | 56 |
| Gcn1_N | -0.158813502 | 50 |
| STAS | -0.157325397 | 56 |
| PP2C | -0.156916199 | 56 |
| SSB | -0.156269766 | 48 |
| Haspin_kinase | -0.15569712 | 56 |
| RmlD_sub_bind | -0.155621871 | 56 |
| Peptidase_M49 | -0.155198317 | 46 |
| MMS1_N | -0.154756008 | 56 |
| GMC_oxred_N | -0.154691687 | 46 |
| SWI-SNF_Ssr4 | -0.154601944 | 46 |
| Cse1 | -0.154280325 | 55 |
| RBM39linker | -0.154276026 | 50 |
| HD | -0.154212469 | 56 |
| Ribosomal_S13 | -0.153091223 | 56 |
| ATP-sulfurylase | -0.151242116 | 51 |
| RasGAP | -0.151055764 | 56 |
| Dynamin_M | -0.150226075 | 56 |
| Dymeclin | -0.150055887 | 53 |
| MRG | -0.149536765 | 56 |
| Aldedh | -0.148940643 | 56 |
| NT-C2 | -0.148775616 | 43 |
| BRE1 | -0.148460562 | 45 |
| CDC27 | -0.148302515 | 47 |
| GalP_UDP_tr_C | -0.146984426 | 51 |
| Sigma54_activat | -0.146932507 | 55 |
| SAE2 | -0.145907182 | 29 |
| LNS2 | -0.145817572 | 54 |
| DHHC | -0.144791123 | 56 |
| TMP-TENI | -0.14397394 | 43 |
| UBX | -0.14183163 | 56 |
| Pam17 | -0.141271984 | 54 |
| Rdx | -0.141029816 | 42 |
| Methyltransf_15 | -0.140719563 | 55 |
| RMI1_N | -0.14055987 | 54 |
| Ins145_P3_rec | -0.140338073 | 34 |
| zf-TRM13_CCCH | -0.1403063 | 41 |
| UBA_4 | -0.139407697 | 55 |
| SNARE | -0.139109504 | 56 |
| DHquinase_I | -0.138795784 | 53 |
| Bact_lectin | -0.138496974 | 54 |
| PGP_phosphatase | -0.137143154 | 55 |
| NAD_binding_4 | -0.136229438 | 56 |
| PH_RBD | -0.136214538 | 31 |
| UCH | -0.134539928 | 56 |
| Tom5 | -0.133420329 | 20 |
| ILVD_EDD | -0.133056042 | 56 |
| BTG | -0.132842916 | 51 |
| G-patch | -0.132334689 | 56 |
| Snapin_Pallidin | -0.131830333 | 45 |
| HPP | -0.1304188 | 43 |
| Clathrin | -0.130389688 | 56 |
| TPP_enzyme_N | -0.13016981 | 56 |
| Med12 | -0.129315901 | 50 |
| Ribosomal_L16 | -0.128314474 | 56 |
| EF-hand_6 | -0.126566623 | 56 |
| ATP-synt | -0.126351606 | 54 |
| PKD_channel | -0.125959558 | 27 |
| CoA_binding | -0.125829479 | 56 |
| zf-C4pol | -0.123808324 | 54 |
| YL1 | -0.122875963 | 43 |
| Uso1_p115_C | -0.12270952 | 39 |
| zf-DNA_Pol | -0.12260048 | 52 |
| IMS | -0.121700537 | 55 |
| Cation_efflux | -0.121586785 | 56 |
| Fe_hyd_lg_C | -0.120105202 | 55 |
| ArsB | -0.119757827 | 45 |

| | | |
|---|---|---|
| TPP_enzyme_M | -0.118747624 | 56 |
| Thioredoxin_4 | -0.117966343 | 47 |
| PGAP1 | -0.117592898 | 56 |
| DUF1713 | -0.117581518 | 52 |
| PolyA_pol_RNAbd | -0.116725751 | 50 |
| Sec7 | -0.116502588 | 56 |
| Ribosomal_L14 | -0.114777602 | 56 |
| RVP_2 | -0.113000457 | 31 |
| NOT2_3_5 | -0.112379335 | 56 |
| Ribos_L4_asso_C | -0.112010538 | 56 |
| Coa1 | -0.111811379 | 55 |
| Peptidase_M17_N | -0.111748005 | 51 |
| PP2C_2 | -0.110937026 | 56 |
| RPEL | -0.110889886 | 45 |
| NAD_binding_11 | -0.110727466 | 55 |
| Ribosom_S12_S23 | -0.110236957 | 56 |
| CBP_BcsQ | -0.109841982 | 51 |
| ApoO | -0.109648185 | 55 |
| MTHFR | -0.10931257 | 56 |
| NIBRIN_BRCT_II | -0.109109181 | 30 |
| Pro_dh | -0.108542047 | 55 |
| DUF2439 | -0.108229527 | 32 |
| DUF1394 | -0.107436735 | 56 |
| Metalloenzyme | -0.106790603 | 56 |
| Abhydrolase_4 | -0.105526784 | 27 |
| PUA_2 | -0.105196968 | 51 |
| Vps26 | -0.105147622 | 55 |
| DHQ_synthase | -0.105135607 | 54 |
| DUF815 | -0.104138367 | 49 |
| PhoLip_ATPase_N | -0.103225 | 56 |
| UME | -0.102748874 | 35 |
| DUF4460 | -0.101593761 | 40 |
| COX4 | -0.101532224 | 55 |
| BCDHK_Adom3 | -0.101206209 | 56 |
| PRAI | -0.101178719 | 55 |
| DUF4112 | -0.101147231 | 55 |
| RRM_5 | -0.100318827 | 55 |
| G-patch_2 | -0.099967211 | 56 |
| Cation_ATPase | -0.099821489 | 56 |
| zf-C3HC4_3 | -0.099501694 | 56 |
| Put_Phosphatase | -0.099479047 | 45 |
| 4HBT | -0.099442811 | 56 |
| DUF3245 | -0.099320276 | 27 |
| ABC_ATPase | -0.098251957 | 46 |
| BUD22 | -0.098233728 | 46 |
| HSF_DNA-bind | -0.096588639 | 56 |
| Pyr_redox_2 | -0.09534665 | 56 |
| Acyl-CoA_dh_2 | -0.094951515 | 56 |
| Sulfatase | -0.094662514 | 50 |
| DUF2415 | -0.094278111 | 54 |
| Patatin | -0.094248933 | 56 |
| Ribosomal_S8 | -0.093658741 | 56 |
| Pinin_SDK_memA | -0.093431739 | 48 |
| Rad51 | -0.093183803 | 56 |
| CHCH | -0.09244122 | 55 |
| CybS | -0.091894927 | 56 |
| SNF2_N | -0.091767899 | 56 |
| GED | -0.091267494 | 56 |
| GCV_H | -0.090058716 | 54 |
| RRM_1 | -0.08884009 | 56 |
| SNase | -0.088503918 | 53 |
| GST_C_6 | -0.088350516 | 53 |
| ResIII | -0.086726288 | 56 |
| Acyl-CoA_dh_N | -0.086195015 | 56 |
| ECH_1 | -0.085596935 | 56 |
| SpoIIE | -0.085420342 | 55 |
| YjeF_N | -0.085324382 | 56 |
| Tcf25 | -0.085020381 | 48 |
| Dna2 | -0.08438235 | 46 |
| zf_CCCH_4 | -0.084077505 | 52 |
| Glyco_hydro_3 | -0.083853787 | 49 |
| GSIII_N | -0.083425578 | 55 |
| TFIID_NTD2 | -0.083378319 | 54 |
| Mit_KHE1 | -0.08330527 | 44 |
| RPN13_C | -0.083141546 | 48 |
| TPP_enzyme_C | -0.082768888 | 56 |
| DMAP_binding | -0.082498302 | 32 |
| CENP-T_C | -0.082392214 | 56 |
| Tub | -0.082321974 | 51 |
| Amidohydro_1 | -0.082161504 | 56 |
| GCP_C_terminal | -0.080455765 | 56 |
| PCMT | -0.080200897 | 56 |
| Thiolase_N | -0.079709119 | 56 |
| PhoLip_ATPase_C | -0.079509963 | 56 |
| Serinc | -0.079134317 | 56 |
| Flavokinase | -0.078792459 | 56 |
| IMPDH | -0.078040919 | 56 |
| Cation_ATPase_N | -0.077980611 | 56 |
| PAC4 | -0.077708528 | 42 |
| TH1 | -0.077611778 | 51 |
| PMSR | -0.076561607 | 56 |
| DUF3384 | -0.076450338 | 46 |
| MOR2-PAG1_N | -0.075243374 | 56 |
| Ydc2-catalyt | -0.075231757 | 44 |
| Ribosomal_L29 | -0.07483607 | 56 |
| IBN_N | -0.0737516 | 56 |
| Ribonuc_L-PSP | -0.072686043 | 56 |
| Dala_Dala_lig_C | -0.071279277 | 56 |

| | | |
|---|---|---|
| ATP-grasp_2 | -0.070737758 | 56 |
| Anoctamin | -0.070329505 | 54 |
| baeRF_family10 | -0.069712699 | 53 |
| Cytochrom_C1 | -0.069410443 | 56 |
| Rhodanese | -0.068792985 | 56 |
| Methyltransf_5 | -0.068577318 | 39 |
| Ribosomal_S13_N | -0.068233283 | 54 |
| Thioredoxin_6 | -0.068149329 | 56 |
| Cyclin_C_2 | -0.067938747 | 54 |
| NifU_N | -0.067308623 | 55 |
| Spo7 | -0.066245232 | 41 |
| 7tm_3 | -0.065716244 | 43 |
| R3H | -0.064700306 | 56 |
| Clathrin_H_link | -0.064362439 | 54 |
| IP_trans | -0.064291788 | 49 |
| Hydrolase_like | -0.064147756 | 56 |
| zf-TFIIIC | -0.064136381 | 38 |
| Med25_VWA | -0.064043997 | 42 |
| hSac2 | -0.063613101 | 51 |
| DUF3605 | -0.063564096 | 46 |
| SAM_decarbox | -0.062741073 | 56 |
| FAD_oxidored | -0.060640307 | 56 |
| FIST | -0.060630284 | 43 |
| Foie-gras_1 | -0.060301232 | 50 |
| NADH_u_ox_C | -0.06000496 | 54 |
| ERGIC_N | -0.058415678 | 55 |
| SLIDE | -0.058035965 | 54 |
| VCBS | -0.056739783 | 53 |
| 14-3-3 | -0.056643213 | 56 |
| RRM_2 | -0.056369392 | 43 |
| Transketolase_C | -0.055846172 | 56 |
| ABC1 | -0.055159084 | 56 |
| CHORD | -0.054600343 | 52 |
| ODC_AZ | -0.053058741 | 53 |
| HECT | -0.052909685 | 56 |
| RrnaAD | -0.052860135 | 56 |
| Piwi | -0.052226502 | 54 |
| PAC3 | -0.051780222 | 44 |
| Ribosomal_S10 | -0.050127611 | 56 |
| Nucleoporin_N | -0.049047841 | 56 |
| Fer4_17 | -0.048958439 | 55 |
| MDM10 | -0.048709305 | 56 |
| Acyl-CoA_dh_1 | -0.048648793 | 56 |
| GTP_EFTU_D3 | -0.047715653 | 56 |
| Spt5_N | -0.046706647 | 46 |
| XRN1_D2_D3 | -0.046370661 | 41 |
| Gln-synt_C-ter | -0.046140102 | 55 |
| HA2 | -0.045795171 | 56 |
| Seryl_tRNA_N | -0.04549756 | 55 |
| GTP_CH_N | -0.043906524 | 48 |
| Mito_carr | -0.043467235 | 56 |
| Ligase_CoA | -0.042980043 | 56 |
| Cation_ATPase_C | -0.042830971 | 56 |
| SDH_beta | -0.04179996 | 55 |
| Prok-JAB | -0.041778605 | 56 |
| Translin | -0.041497364 | 41 |
| HAD | -0.040748293 | 56 |
| SMK-1 | -0.040732929 | 54 |
| GATase_7 | -0.040664494 | 56 |
| CwfJ_C_2 | -0.039554581 | 53 |
| BORCS8 | -0.038855579 | 33 |
| COP-gamma_platf | -0.038488641 | 56 |
| Pribosyltran_N | -0.037903549 | 56 |
| PUL | -0.037182087 | 55 |
| GST_N_4 | -0.037042429 | 54 |
| USP8_dimer | -0.035399907 | 48 |
| SH3BGR | -0.034453228 | 53 |
| PrmA | -0.034397989 | 56 |
| PseudoU_synth_1 | -0.034378878 | 56 |
| Gal_mutarotas_2 | -0.034160618 | 55 |
| Carb_kinase | -0.033948152 | 55 |
| UBA | -0.033381501 | 56 |
| Thiolase_C | -0.032117778 | 56 |
| Brr6_like_C_C | -0.032097987 | 51 |
| Ribosomal_S14 | -0.031403755 | 56 |
| ATP-synt_DE_N | -0.031215186 | 54 |
| Allantoicase | -0.031020436 | 54 |
| Sterile | -0.030997146 | 35 |
| ArgoN | -0.030771017 | 54 |
| FF | -0.030458353 | 55 |
| Polyketide_cyc | -0.030452697 | 45 |
| APS_kinase | -0.030387909 | 51 |
| EFG_IV | -0.030112666 | 56 |
| NAC | -0.029768792 | 55 |
| SAPS | -0.029568503 | 55 |
| PIN_4 | -0.029391199 | 56 |
| SRPRB | -0.02938338 | 56 |
| PEMT | -0.028527832 | 56 |
| RAB3GAP2_N | -0.028389884 | 44 |
| Urease_alpha | -0.028385671 | 49 |
| Far-17a_AIG1 | -0.026123501 | 47 |
| Dynactin_p22 | -0.025975755 | 50 |
| SMN | -0.025523514 | 40 |
| FAD_binding_7 | -0.023627151 | 29 |
| DUF3835 | -0.022206311 | 36 |
| MTS | -0.021951932 | 56 |
| RAMP4 | -0.021158409 | 36 |
| MA3 | -0.020989845 | 56 |

| | | |
|---|---|---|
| FKBP_C | -0.019334316 | 56 |
| Hydrolase | -0.018777926 | 56 |
| Porin_3 | -0.018763309 | 56 |
| DPPIV_N | -0.018489828 | 54 |
| Fer2_3 | -0.01742421 | 55 |
| Lipase_GDSL | -0.017396053 | 55 |
| Urease_gamma | -0.016541376 | 49 |
| Urease_beta | -0.016541376 | 49 |
| Hpt | -0.016207777 | 53 |
| HlyIII | -0.015886827 | 56 |
| PEPCK_ATP | -0.014058892 | 55 |
| Ureidogly_lyase | -0.01338247 | 52 |
| Ribosomal_L3 | -0.013113743 | 56 |
| Homeodomain | -0.012884899 | 56 |
| Biotin_carb_C | -0.012627653 | 56 |
| DRMBL | -0.012424782 | 54 |
| Nro1 | -0.011479928 | 45 |
| S4 | -0.01136385 | 56 |
| Seipin | -0.011307884 | 55 |
| Ribonuc_red_sm | -0.010892984 | 56 |
| GCP_N_terminal | -0.010279001 | 56 |
| acVLRF1 | -0.009649104 | 54 |
| BLM10_mid | -0.009641247 | 49 |
| ClpB_D2-small | -0.009547028 | 56 |
| FANCI_S4 | -0.006806109 | 50 |
| FTHFS | -0.005588093 | 55 |
| Ribosomal_S4 | -0.005510775 | 55 |
| CGI-121 | -0.005487154 | 49 |
| eIF2A | -0.005352931 | 56 |
| GATase_2 | -0.004922821 | 54 |
| Exo_endo_phos | -0.004880915 | 56 |
| AIF_C | -0.00450165 | 41 |
| Fer4 | -0.004210181 | 55 |
| NopRA1 | -0.004118008 | 49 |
| RFC1 | -0.003701929 | 53 |
| DLH | -0.003389092 | 56 |
| Solute_trans_a | -0.002999107 | 56 |
| mRNA_cap_C | -0.002712227 | 54 |
| GATase_6 | -0.002597457 | 56 |
| SCO1-SenC | -0.002474957 | 56 |
| Cut8 | -0.002148646 | 55 |
| Lipin_N | -0.001805626 | 49 |
| Peptidase_M24 | -0.00055992 | 56 |
| Rrp44_S1 | -0.000288083 | 51 |
| Metallophos | -0.000285393 | 56 |
| Asn_synthase | 0.000447918 | 55 |
| zf-C2H2_2 | 0.000563682 | 56 |
| NAD_binding_1 | 0.002056002 | 56 |
| DUF2013 | 0.002287386 | 54 |
| Robl_LC7 | 0.002938863 | 54 |
| DOT1 | 0.003160249 | 56 |
| TAL_FSA | 0.003694647 | 56 |
| V-SNARE | 0.00655016 | 53 |
| PfkB | 0.007563637 | 56 |
| H2TH | 0.00758808 | 37 |
| Alba | 0.007835569 | 50 |
| HEAT_PBS | 0.008319467 | 56 |
| Sulfate_transp | 0.010845784 | 56 |
| Biotin_carb_N | 0.010919118 | 56 |
| RCC1_2 | 0.011445984 | 56 |
| Pol_alpha_B_N | 0.011847689 | 44 |
| SH2 | 0.012470236 | 39 |
| zf-AN1 | 0.012916263 | 56 |
| Creatinase_N_2 | 0.013712625 | 55 |
| DcpS_C | 0.014461675 | 56 |
| Glyco_transf_15 | 0.015065579 | 53 |
| CDC24_OB3 | 0.016271644 | 44 |
| SLC35F | 0.01628434 | 56 |
| ABC2_membrane | 0.016376036 | 56 |
| PYC_OADA | 0.016834383 | 53 |
| COPIIcoated_ERV | 0.017124297 | 56 |
| Epimerase | 0.01754244 | 56 |
| Iso_dh | 0.018394901 | 56 |
| SDH_alpha | 0.018555428 | 56 |
| KilA-N | 0.019164511 | 41 |
| COQ9 | 0.019270525 | 50 |
| Band_7 | 0.019635398 | 56 |
| MipZ | 0.019899641 | 55 |
| Coatomer_g_Cpla | 0.020114441 | 56 |
| Ctr | 0.020536832 | 56 |
| Ribosomal_S7 | 0.021217567 | 56 |
| Sec5 | 0.022575277 | 55 |
| DnaJ_C | 0.023170331 | 56 |
| Peptidase_M42 | 0.023740508 | 54 |
| FMN_dh | 0.023773344 | 56 |
| Tudor-knot | 0.024005913 | 56 |
| IPK | 0.024836076 | 55 |
| SNAP | 0.025885749 | 56 |
| Metallophos_2 | 0.025950536 | 54 |
| Peptidase_M24_C | 0.026681297 | 56 |
| Toprim | 0.028470493 | 56 |
| BCNT | 0.028567486 | 45 |
| Striatin | 0.028697132 | 50 |
| Init_tRNA_PT | 0.02956701 | 47 |
| RNase_P_pop3 | 0.029831196 | 43 |
| BRCT | 0.031000132 | 56 |
| PC_rep | 0.031277515 | 56 |
| AIM24 | 0.031378906 | 49 |

| | | |
|---|---|---|
| COX5A | 0.031825456 | 55 |
| AAA_5 | 0.032471565 | 56 |
| HAND | 0.0325151 | 54 |
| Atg8 | 0.033148577 | 55 |
| HIRA_B | 0.033813302 | 34 |
| MreB_Mbl | 0.034255426 | 56 |
| Efg1 | 0.034367611 | 52 |
| P_proprotein | 0.0346264 | 55 |
| FAD_binding_6 | 0.035122057 | 56 |
| SF3a60_bindingd | 0.03557245 | 47 |
| CAF1A | 0.035741523 | 49 |
| Med17 | 0.036576858 | 52 |
| Ski2_N | 0.037130572 | 51 |
| Fer4_10 | 0.038188429 | 55 |
| Nucleoporin_C | 0.038243459 | 54 |
| Lung_7-TM_R | 0.038422544 | 55 |
| Aldose_epim | 0.038431447 | 56 |
| PAPS_reduct | 0.039241501 | 56 |
| Urb2 | 0.039686847 | 47 |
| GATase | 0.039693249 | 56 |
| RRP14 | 0.039975402 | 50 |
| NOGCT | 0.040324244 | 55 |
| XPG_N | 0.041344264 | 56 |
| Questin_oxidase | 0.042568923 | 50 |
| CRAL_TRIO | 0.042738364 | 56 |
| Ribosomal_L6 | 0.042786596 | 56 |
| OB_NTP_bind | 0.043333203 | 56 |
| NDUFB10 | 0.044380271 | 40 |
| PRK | 0.044463754 | 55 |
| TBP | 0.045722264 | 56 |
| CDP-OH_P_transf | 0.046161513 | 56 |
| Ribonuclease_3 | 0.047696385 | 56 |
| Ribosomal_L22e | 0.047789674 | 55 |
| dCMP_cyt_deam_2 | 0.049030478 | 51 |
| PUA | 0.049738164 | 56 |
| Nic96 | 0.049857448 | 55 |
| SANT_DAMP1_like | 0.049990159 | 53 |
| eIF3_subunit | 0.051142696 | 56 |
| ThrE | 0.052081146 | 53 |
| Succ_DH_flav_C | 0.052621414 | 56 |
| ICMT | 0.053334236 | 55 |
| Sad1_UNC | 0.053909446 | 56 |
| Cullin | 0.054885239 | 56 |
| zf-MYST | 0.055304969 | 56 |
| Pyr_redox_dim | 0.055726942 | 56 |
| DUF2340 | 0.056593988 | 47 |
| HBS1_N | 0.056742362 | 43 |
| HIG_1_N | 0.057191328 | 56 |
| PAC2 | 0.058305032 | 49 |
| NRDE-2 | 0.058859742 | 49 |
| Atg14 | 0.059528141 | 48 |
| TP_methylase | 0.05990607 | 56 |
| KOW | 0.060481217 | 56 |
| Phosphodiest | 0.060922296 | 56 |
| PRELI | 0.061469975 | 56 |
| Clat_adaptor_s | 0.062177911 | 56 |
| MnmE_helical | 0.063047722 | 52 |
| Clathrin_lg_ch | 0.063338595 | 54 |
| DUF4451 | 0.064408811 | 56 |
| DFP | 0.064812224 | 54 |
| KR | 0.065792527 | 56 |
| DegT_DnrJ_EryC1 | 0.065882193 | 56 |
| GUCT | 0.066087095 | 49 |
| SAP30_Sin3_bdg | 0.066225619 | 44 |
| Mur_ligase_M | 0.066350132 | 53 |
| Biotin_lipoyl | 0.066376398 | 56 |
| APG17 | 0.066452777 | 54 |
| Nup192 | 0.066620705 | 56 |
| tRNA_bind | 0.066726224 | 56 |
| Apt1 | 0.067283453 | 44 |
| A_deaminase | 0.067444491 | 56 |
| RIO1 | 0.067489198 | 56 |
| IPP-2 | 0.067691926 | 56 |
| GIDA | 0.067951861 | 56 |
| Pribosyl_synth | 0.068441048 | 56 |
| Elong_Iki1 | 0.069548466 | 55 |
| DHH | 0.069669396 | 43 |
| zf-CCCH | 0.069719378 | 56 |
| Pyr_redox | 0.069907048 | 56 |
| RQC | 0.069947861 | 53 |
| BRAP2 | 0.07041172 | 53 |
| RGS-like | 0.070441803 | 24 |
| DinB_2 | 0.071632677 | 39 |
| zf-Tim10_DDP | 0.071797482 | 56 |
| SE | 0.07257647 | 54 |
| bZIP_Maf | 0.072666645 | 54 |
| MOZ_SAS | 0.07284143 | 56 |
| UFD1 | 0.073336144 | 56 |
| DASH_Dad4 | 0.074155027 | 39 |
| AdoMet_MTase | 0.074585175 | 54 |
| Transthyretin | 0.075003146 | 50 |
| Torus | 0.075191164 | 56 |
| Vac_ImportDeg | 0.075346232 | 56 |
| Med9 | 0.075592576 | 40 |
| HIT | 0.076380996 | 56 |
| PEP_mutase | 0.078162546 | 55 |
| Transket_pyr | 0.080272021 | 56 |
| ADSL_C | 0.08034161 | 55 |

| | | |
|---|---|---|
| Tcp11 | 0.080690504 | 51 |
| ATPase | 0.081094655 | 41 |
| TAF | 0.081810216 | 55 |
| DUF5599 | 0.081923438 | 42 |
| eIF-5_eIF-2B | 0.082630928 | 56 |
| SCAI | 0.083165234 | 55 |
| GCD14 | 0.083503427 | 56 |
| Rft-1 | 0.084604032 | 56 |
| COMPASS-Shg1 | 0.084756939 | 41 |
| Ribosomal_S11 | 0.084877346 | 56 |
| Helicase_C | 0.084930902 | 56 |
| Lactamase_B_2 | 0.085129939 | 56 |
| CRAL_TRIO_2 | 0.085749799 | 56 |
| EFG_II | 0.086781185 | 56 |
| TFIIF_alpha | 0.087466809 | 47 |
| FAD_binding_4 | 0.088600565 | 56 |
| PI3_PI4_kinase | 0.088796018 | 56 |
| Dicer_dimer | 0.089231234 | 50 |
| CMAS | 0.089948869 | 56 |
| Porphobil_deam | 0.090388745 | 54 |
| PPR_1 | 0.091274974 | 55 |
| SAC3_GANP | 0.091292742 | 56 |
| UvrD_C | 0.091391869 | 39 |
| FAA_hydrolase | 0.091460649 | 56 |
| Peptidase_M76 | 0.09157571 | 54 |
| 2OG-FeII_Oxy_4 | 0.091859154 | 56 |
| Fer2_4 | 0.092271421 | 56 |
| NADH-G_4Fe-4S_3 | 0.092271421 | 56 |
| ArgoMid | 0.09304372 | 54 |
| ANAPC5 | 0.093324277 | 42 |
| DNA_pol_A | 0.093411551 | 52 |
| zinc_ribbon_10 | 0.09341462 | 56 |
| CSTF_C | 0.094698236 | 46 |
| p450 | 0.095171001 | 56 |
| ADK | 0.095533545 | 56 |
| DUF1279 | 0.095643842 | 55 |
| AMPK1_CBM | 0.095977572 | 55 |
| BAR_2 | 0.096133182 | 56 |
| Cupin_4 | 0.096365622 | 47 |
| Cyanate_lyase | 0.096402764 | 37 |
| GDI | 0.096479122 | 56 |
| LRR_9 | 0.096512662 | 56 |
| Methyltransf_33 | 0.096694022 | 50 |
| MBOAT | 0.097574779 | 56 |
| M16C_assoc | 0.09768973 | 53 |
| PQ-loop | 0.097784347 | 56 |
| UPF1_Zn_bind | 0.097975483 | 44 |
| ICL | 0.098751637 | 56 |
| zf-C5HC2 | 0.09940837 | 49 |
| FtsJ | 0.099792949 | 56 |
| E1-E2_ATPase | 0.099866943 | 56 |
| Guanylate_cyc_2 | 0.100138464 | 42 |
| CAMSAP_CH | 0.100499291 | 54 |
| Yip1 | 0.100767736 | 56 |
| CDC73_N | 0.100990129 | 48 |
| Mon1 | 0.101284194 | 52 |
| FA_desaturase | 0.101572255 | 56 |
| HABP4_PAI-RBP1 | 0.10160477 | 44 |
| tRNA_U5-meth_tr | 0.10187763 | 54 |
| MEA1 | 0.102006472 | 31 |
| Mrr_cat | 0.102412377 | 33 |
| Bystin | 0.102556262 | 56 |
| Ribosomal_L4 | 0.10263062 | 56 |
| MKT1_C | 0.103207691 | 40 |
| Cohesin_HEAT | 0.103406324 | 49 |
| Protoglobin | 0.103465217 | 41 |
| I_LWEQ | 0.103857104 | 54 |
| CDC24_OB1 | 0.10441974 | 36 |
| UPRTase | 0.104991776 | 56 |
| PI3Ka | 0.105030533 | 56 |
| YccF | 0.105104257 | 53 |
| COX7C | 0.105276226 | 53 |
| Lectin_leg-like | 0.105282043 | 56 |
| SNF5 | 0.105413668 | 56 |
| POPLD | 0.105467562 | 53 |
| ATG101 | 0.105651397 | 48 |
| Ssu72 | 0.106499815 | 50 |
| GST_N_2 | 0.106832051 | 56 |
| DKCLD | 0.107761708 | 54 |
| COQ7 | 0.107769692 | 51 |
| GTP1_OBG | 0.108003459 | 52 |
| SUZ | 0.108079344 | 45 |
| RNA_pol | 0.108123797 | 56 |
| Cullin_Nedd8 | 0.108228294 | 56 |
| DEAD | 0.108330712 | 56 |
| PhyH | 0.109533534 | 53 |
| TCTP | 0.110371358 | 48 |
| RNase_P_Rpp14 | 0.110497298 | 53 |
| DUF3510 | 0.111267996 | 40 |
| PseudoU_synth_2 | 0.111320967 | 56 |
| Ferric_reduct | 0.111708255 | 56 |
| NIPSNAP | 0.112060715 | 54 |
| Aldolase_II | 0.112104656 | 56 |
| Tom37 | 0.112454176 | 55 |
| TrmE_N | 0.113089315 | 50 |
| Ala_racemase_N | 0.11322434 | 55 |
| DnaJ | 0.11410852 | 56 |
| NARG2_C | 0.114221925 | 23 |

| | | |
|---|---|---|
| Methyltransf_28 | 0.114486952 | 56 |
| SQHop_cyclase_C | 0.114603129 | 55 |
| CSTF2_hinge | 0.114673939 | 51 |
| TRAM1 | 0.115008981 | 54 |
| 2-oxogl_dehyd_N | 0.115436174 | 56 |
| MoCF_biosynth | 0.115873208 | 55 |
| tRNA-synt_His | 0.116341422 | 56 |
| Histone | 0.117449242 | 56 |
| DUF3337 | 0.118758511 | 51 |
| PIN_9 | 0.118787579 | 52 |
| eIF-5a | 0.119672684 | 56 |
| Hist_deacetyl | 0.11975262 | 56 |
| NADH-u_ox-rdase | 0.122134021 | 56 |
| CBM_48 | 0.122423634 | 53 |
| Importin_rep_6 | 0.122543106 | 52 |
| adh_short_C2 | 0.122579053 | 56 |
| zf-NF-X1 | 0.122951473 | 48 |
| DUF1712 | 0.123458265 | 53 |
| DUF3437 | 0.123501768 | 50 |
| Succ_CoA_lig | 0.123986477 | 54 |
| SCA7 | 0.124009777 | 53 |
| CN_hydrolase | 0.124082115 | 56 |
| CDC48_N | 0.124221325 | 54 |
| KAP | 0.1252385 | 54 |
| PWWP | 0.125319289 | 55 |
| mRNA_cap_enzyme | 0.125765608 | 56 |
| CPSF100_C | 0.126778864 | 50 |
| ATP_sub_h | 0.12684698 | 54 |
| Catalase-rel | 0.127200775 | 52 |
| VWA_2 | 0.128679531 | 55 |
| SRP_TPR_like | 0.128687553 | 46 |
| Fer4_7 | 0.129026331 | 54 |
| Fer4_9 | 0.129026331 | 54 |
| MIF4G | 0.129029871 | 56 |
| AdoHcyase | 0.129359511 | 55 |
| TPPII | 0.129940739 | 48 |
| CoA_binding_2 | 0.12998602 | 53 |
| ERCC4 | 0.130000721 | 56 |
| Coq4 | 0.131341367 | 53 |
| Ndc1_Nup | 0.131671574 | 50 |
| PPP4R2 | 0.131680988 | 45 |
| HbrB | 0.132012612 | 52 |
| Zn_ribbon_17 | 0.132159745 | 52 |
| Tubulin | 0.132684851 | 56 |
| Muskelin_N | 0.133037813 | 55 |
| adh_short | 0.133154789 | 56 |
| GTP_EFTU | 0.133534632 | 56 |
| NIF | 0.134089239 | 56 |
| Rpp20 | 0.134501381 | 43 |
| Ribosomal_L31 | 0.135850241 | 46 |
| eIF-6 | 0.136169916 | 56 |
| AAA_2 | 0.136417697 | 56 |
| HhH-GPD | 0.136501221 | 56 |
| DNA_photolyase | 0.136903754 | 31 |
| WHEP-TRS | 0.137346135 | 45 |
| Acyl_CoA_thio | 0.137727295 | 46 |
| HAD_SAK_1 | 0.137914425 | 54 |
| Adaptin_binding | 0.137966359 | 39 |
| Ribosomal_S15 | 0.137979178 | 56 |
| EMC1_C | 0.138846658 | 54 |
| GARS_A | 0.139214253 | 56 |
| COBRA1 | 0.140112835 | 53 |
| YL1_C | 0.141481372 | 54 |
| Tyr-DNA_phospho | 0.142394001 | 55 |
| CRAL_TRIO_N | 0.142547409 | 56 |
| Pep3_Vps18 | 0.142561685 | 56 |
| CPSase_sm_chain | 0.142730312 | 56 |
| GTP_cyclohydro2 | 0.143226378 | 55 |
| DUF3453 | 0.143753804 | 49 |
| Coprogen_oxidas | 0.143841239 | 56 |
| NTF2 | 0.144170216 | 56 |
| Pre-PUA | 0.144365102 | 56 |
| MT-A70 | 0.145265381 | 56 |
| DUF1690 | 0.145882182 | 54 |
| PSS | 0.146101864 | 49 |
| Cnd1 | 0.146136852 | 56 |
| COG2 | 0.147661353 | 49 |
| 3Beta_HSD | 0.147712128 | 56 |
| Acatn | 0.148050484 | 52 |
| Mid1 | 0.148160347 | 43 |
| Lactamase_B_4 | 0.149104269 | 53 |
| tRNA-synt_2d | 0.149290029 | 56 |
| FSH1 | 0.150770699 | 56 |
| APG6_N | 0.151010349 | 51 |
| DnaB_C | 0.152367375 | 55 |
| DFRP_C | 0.152379039 | 56 |
| Aconitase_C | 0.152696942 | 56 |
| GrpE | 0.152796618 | 56 |
| Sec61_beta | 0.152856915 | 54 |
| zf-CCHC_2 | 0.153072516 | 55 |
| Cyclin_N | 0.153550057 | 56 |
| zf-CCHC_3 | 0.153971319 | 47 |
| Med11 | 0.154125511 | 43 |
| zf-BED | 0.154126163 | 27 |
| Coatomer_E | 0.154926617 | 55 |
| NOG1 | 0.155199283 | 56 |
| Neurochondrin | 0.155378846 | 54 |
| Lyase_1 | 0.155435514 | 56 |

| | | |
|---|---|---|
| DUF1769 | 0.156567336 | 37 |
| MFS_4 | 0.157273523 | 45 |
| ATP-synt_G | 0.157291008 | 56 |
| OTCace | 0.158089586 | 56 |
| CbiA | 0.158678698 | 56 |
| Helicase_RecD | 0.159506144 | 55 |
| JAB | 0.160219387 | 56 |
| Choline_kinase | 0.160509632 | 56 |
| tRNA_m1G_MT | 0.160743921 | 56 |
| Sec66 | 0.16080232 | 54 |
| AAA_14 | 0.161294404 | 34 |
| PIG-F | 0.161367094 | 48 |
| TAF8_C | 0.161853358 | 53 |
| RGS | 0.162199901 | 56 |
| Porphobil_deamC | 0.162595607 | 53 |
| Glutaredoxin | 0.162778699 | 56 |
| AAA | 0.162851426 | 56 |
| Alpha-amylase | 0.162953866 | 55 |
| ATG_C | 0.163220888 | 56 |
| Rtt106 | 0.164119596 | 56 |
| OTCace_N | 0.164510271 | 56 |
| Band_7_C | 0.164906563 | 55 |
| UDG | 0.165590159 | 56 |
| Myosin_TH1 | 0.165616017 | 54 |
| UPF0016 | 0.165630042 | 54 |
| NAP | 0.167148409 | 56 |
| Fapy_DNA_glyco | 0.16729735 | 31 |
| DTW | 0.167392891 | 55 |
| FhuF | 0.168539585 | 49 |
| RNA_pol_Rpb8 | 0.168851026 | 55 |
| 2-oxoacid_dh | 0.16886082 | 56 |
| FmiP_Thoc5 | 0.169627504 | 50 |
| Mlh1_C | 0.169983348 | 55 |
| GTP_EFTU_D2 | 0.170130066 | 56 |
| Beta-lactamase | 0.170482886 | 33 |
| DUF3543 | 0.170916164 | 51 |
| XPA_C | 0.170951447 | 55 |
| HisG | 0.17128944 | 54 |
| UreD | 0.171503023 | 47 |
| vATP-synt_E | 0.171662394 | 56 |
| PolyA_pol | 0.171891832 | 54 |
| ETF_QO | 0.171908745 | 54 |
| HGTP_anticodon | 0.172191444 | 56 |
| ANAPC4_WD40 | 0.173078822 | 56 |
| RNA_pol_Rpc82 | 0.173735529 | 53 |
| EST1_DNA_bind | 0.173832498 | 53 |
| Tfb4 | 0.17404666 | 56 |
| SMC_N | 0.174437551 | 56 |
| DUF3767 | 0.174488128 | 53 |
| CSN8_PSD8_EIF3K | 0.174860871 | 56 |
| B12-binding_2 | 0.174960549 | 43 |
| iPGM_N | 0.17581039 | 56 |
| zinc_ribbon_6 | 0.176651952 | 51 |
| PSP | 0.177363772 | 56 |
| Q_salvage | 0.177576755 | 54 |
| Coatomer_b_Cpla | 0.178798038 | 55 |
| TIP49 | 0.179568504 | 56 |
| TMF_DNA_bd | 0.179865303 | 31 |
| Homoserine_dh | 0.180865706 | 55 |
| BORCS6 | 0.18105858 | 47 |
| Glyoxalase_5 | 0.181957825 | 42 |
| ETF | 0.182361544 | 56 |
| DMT_YdcZ | 0.182411696 | 32 |
| Bestrophin | 0.183528446 | 50 |
| Ribosomal_L9_C | 0.184645112 | 45 |
| XPG_I | 0.184796131 | 56 |
| DHFR_1 | 0.184915085 | 55 |
| Rad17 | 0.185402017 | 56 |
| Adaptin_N | 0.185643316 | 56 |
| Dynein_light | 0.186015964 | 55 |
| Acetyltransf_9 | 0.186115652 | 36 |
| Med26 | 0.1862643 | 56 |
| Dus | 0.18641468 | 56 |
| SRP9-21 | 0.186756878 | 45 |
| ACAS_N | 0.187289953 | 55 |
| BK_channel_a | 0.187361581 | 49 |
| Lipase_GDSL_2 | 0.187469151 | 55 |
| Helicase_C_2 | 0.18747045 | 56 |
| CAP_C | 0.188224927 | 54 |
| AAA_22 | 0.188852639 | 56 |
| E1_DerP2_DerF2 | 0.1893644 | 56 |
| NAD_kinase | 0.189618221 | 56 |
| Spt5-NGN | 0.189634995 | 50 |
| TP6A_N | 0.189796477 | 50 |
| zf-C2HC5 | 0.190154384 | 49 |
| SIN1_PH | 0.190691851 | 53 |
| Med6 | 0.190959681 | 54 |
| DUF667 | 0.191718185 | 47 |
| CENP-W | 0.191866735 | 25 |
| NADH_dhqG_C | 0.191873299 | 56 |
| Beta_elim_lyase | 0.193093216 | 56 |
| Rer1 | 0.194566253 | 55 |
| HGTP_anticodon2 | 0.194574037 | 55 |
| PAT1 | 0.194971974 | 56 |
| CS | 0.194982655 | 56 |
| Ubiq_cyt_C_chap | 0.195053732 | 54 |
| EMP24_GP25L | 0.195328947 | 56 |
| AdenylateSensor | 0.195703597 | 52 |

| | | |
|---|---|---|
| Catalase | 0.195813206 | 53 |
| Rit1_C | 0.195815205 | 49 |
| zf-UBP | 0.195830197 | 56 |
| RRM_9 | 0.195879505 | 38 |
| SUI1 | 0.196607791 | 56 |
| MutL_C | 0.196651004 | 53 |
| Chlorophyllase2 | 0.197376223 | 23 |
| Trp_Tyr_perm | 0.197835932 | 53 |
| PTCB-BRCT | 0.197891309 | 56 |
| Mago_nashi | 0.198196326 | 53 |
| MscL | 0.198214244 | 23 |
| Sacchrp_dh_NADP | 0.198963397 | 56 |
| COX5B | 0.199322713 | 55 |
| NARP1 | 0.200082045 | 54 |
| Prefoldin_3 | 0.20038168 | 44 |
| RNF220 | 0.200888523 | 49 |
| RRM_4 | 0.200983989 | 53 |
| Redoxin | 0.201264845 | 56 |
| XRN1_D1 | 0.201463935 | 41 |
| PRKCSH-like | 0.201588657 | 56 |
| TruB_C_2 | 0.201632763 | 54 |
| Got1 | 0.201646669 | 56 |
| PCO_ADO | 0.202402718 | 39 |
| LMBR1 | 0.202486614 | 54 |
| tRNA_int_end_N2 | 0.20267881 | 43 |
| RPT | 0.203101285 | 41 |
| DnaJ_CXXCXGXG | 0.203235341 | 56 |
| WES_acyltransf | 0.203404358 | 23 |
| Bot1p | 0.203512426 | 54 |
| Scs3p | 0.204158981 | 52 |
| Alk_phosphatase | 0.204296491 | 55 |
| STAG | 0.204417278 | 53 |
| Cytochrome_CBB3 | 0.205538859 | 55 |
| cobW | 0.206071086 | 56 |
| ATP11 | 0.206118986 | 53 |
| Telomere_reg-2 | 0.206559069 | 50 |
| LRR_4 | 0.206898918 | 56 |
| zf-C2H2_jaz | 0.206947977 | 56 |
| DUF4743 | 0.207538924 | 54 |
| APG12 | 0.207568483 | 56 |
| MutS_III | 0.207898213 | 56 |
| Tubulin_C | 0.208288487 | 56 |
| Peptidase_M48 | 0.208560206 | 55 |
| Ribonuc_P_40 | 0.209456596 | 55 |
| Cytochrom_C | 0.209530988 | 56 |
| Ima1_N | 0.20986733 | 45 |
| Gti1_Pac2 | 0.210330135 | 56 |
| FeoB_N | 0.210501689 | 56 |
| LsmAD | 0.210786308 | 54 |
| S1 | 0.210787377 | 56 |
| MutS_IV | 0.210898626 | 56 |
| ATG22 | 0.211277941 | 51 |
| CDC48_2 | 0.211810764 | 54 |
| Fe-ADH | 0.212045935 | 49 |
| UQ_con | 0.212492503 | 56 |
| zf-met | 0.212880216 | 56 |
| AMP-binding_C | 0.213741298 | 56 |
| Med20 | 0.213828916 | 52 |
| Fe-S_biosyn | 0.214379572 | 56 |
| BCAS2 | 0.215966335 | 54 |
| V_ATPase_I | 0.216112381 | 55 |
| SGS | 0.216145439 | 53 |
| NMD3 | 0.216167883 | 56 |
| Methyltr_RsmB-F | 0.216257137 | 56 |
| ABC_membrane_2 | 0.216813212 | 56 |
| eRF1_1 | 0.217396432 | 56 |
| 1-cysPrx_C | 0.21740602 | 56 |
| AAA_lid_5 | 0.217914252 | 52 |
| KH_8 | 0.21814392 | 56 |
| SpoU_sub_bind | 0.218598154 | 43 |
| COPI_C | 0.219101352 | 55 |
| HDA2-3 | 0.219154033 | 50 |
| Med7 | 0.220057504 | 54 |
| S-methyl_trans | 0.220164835 | 49 |
| Esterase | 0.22046896 | 54 |
| DUF726 | 0.220615812 | 55 |
| TAF4 | 0.222275845 | 50 |
| DUF2781 | 0.222852933 | 53 |
| POP1 | 0.223129604 | 55 |
| Gpi1 | 0.223874814 | 52 |
| MutS_II | 0.224267857 | 56 |
| eIF-1a | 0.224341013 | 56 |
| CBFD_NFYB_HMF | 0.224435596 | 56 |
| TFIID_30kDa | 0.225249848 | 54 |
| Citrate_bind | 0.225948324 | 55 |
| F_actin_cap_B | 0.22695808 | 55 |
| Aminotran_1_2 | 0.227307919 | 56 |
| HBB | 0.227330786 | 55 |
| Ribosomal_S17_N | 0.227573911 | 55 |
| FATC | 0.227674666 | 56 |
| PigN | 0.227925938 | 54 |
| Aha1_N | 0.228050842 | 56 |
| CwfJ_C_1 | 0.22806814 | 56 |
| HHH | 0.228293992 | 51 |
| Ribonuclease_P | 0.228841172 | 30 |
| Leuk-A4-hydro_C | 0.228937528 | 54 |
| DSHCT | 0.229413334 | 56 |
| Radical_SAM | 0.229446938 | 56 |

| | | |
|---|---|---|
| RLI | 0.229564804 | 56 |
| SGT1 | 0.229909362 | 53 |
| Alpha-amylase_C | 0.229970431 | 55 |
| Phosphorylase | 0.230034577 | 46 |
| IGPD | 0.231071808 | 54 |
| HisG_C | 0.231527161 | 53 |
| TFIIF_beta | 0.231627973 | 53 |
| IucA_IucC | 0.23202557 | 48 |
| DUF1325 | 0.23243868 | 52 |
| Diphthami_syn_2 | 0.233442774 | 56 |
| MGS | 0.233970673 | 56 |
| Prp18 | 0.234729474 | 52 |
| Cactin_mid | 0.234942702 | 53 |
| AFG1_ATPase | 0.235148193 | 55 |
| NAD_binding_6 | 0.236053825 | 56 |
| DPM3 | 0.236839858 | 50 |
| ARID | 0.237051198 | 56 |
| Glyco_trans_4_4 | 0.237202935 | 55 |
| CPSase_L_D3 | 0.237212584 | 56 |
| AlaDh_PNT_N | 0.237279597 | 56 |
| SEP | 0.239583191 | 55 |
| ATE_N | 0.239686545 | 50 |
| DUF1604 | 0.240289021 | 51 |
| Sas10_Utp3 | 0.240378762 | 55 |
| RNA_pol_Rpb6 | 0.240455958 | 56 |
| UPF0113 | 0.240904846 | 55 |
| NCA2 | 0.241076427 | 55 |
| Vps39_2 | 0.241254822 | 55 |
| Nas2_N | 0.241269026 | 52 |
| TMF_TATA_bd | 0.241341666 | 44 |
| SCAMP | 0.241412799 | 53 |
| eIF3g | 0.241476963 | 54 |
| AAA_31 | 0.242888738 | 55 |
| DUF1775 | 0.243253945 | 43 |
| CENP-S | 0.24337705 | 46 |
| Carn_acyltransf | 0.243592331 | 56 |
| Vps52 | 0.244386459 | 56 |
| BORCS7 | 0.244844912 | 32 |
| Rho_GDI | 0.2450014 | 56 |
| SNAPc_SNAP43 | 0.245010993 | 50 |
| Glyco_H_20C_C | 0.245190671 | 49 |
| LCAT | 0.245229394 | 55 |
| Utp12 | 0.245349203 | 56 |
| DUF2373 | 0.245389718 | 45 |
| FAT | 0.245496312 | 56 |
| HECT_2 | 0.24676225 | 54 |
| Glyco_trans_4_2 | 0.2469292 | 32 |
| Putative_PNPOx | 0.247231604 | 50 |
| U-box | 0.247461732 | 56 |
| eIF_4EBP | 0.248124267 | 47 |
| Oxidored_q6 | 0.248372332 | 55 |
| RNase_PH_C | 0.248555761 | 56 |
| TANGO2 | 0.2492095 | 52 |
| CPSase_L_D2 | 0.249562981 | 56 |
| TMEM70 | 0.249852818 | 39 |
| Sld5 | 0.25011144 | 56 |
| bVLRF1 | 0.250362451 | 54 |
| ATE_C | 0.250648389 | 52 |
| DUF3381 | 0.251580538 | 54 |
| PI31_Prot_C | 0.251975643 | 52 |
| GHMP_kinases_N | 0.252745002 | 56 |
| CWC25 | 0.252847358 | 52 |
| Mss4 | 0.253194297 | 53 |
| POT1 | 0.253367891 | 29 |
| tRNA_edit | 0.253455664 | 54 |
| Coatamer_beta_C | 0.253707791 | 55 |
| DUF1115 | 0.25377669 | 53 |
| ATP-synt_D | 0.254036806 | 56 |
| Dynamitin | 0.254460938 | 54 |
| Molybdopterin | 0.25459043 | 56 |
| Nse4-Nse3_bdg | 0.254645106 | 49 |
| DUF572 | 0.254818749 | 55 |
| ORMDL | 0.255073149 | 55 |
| rRNA_proc-arch | 0.255268039 | 56 |
| SNO | 0.25539169 | 55 |
| U1snRNP70_N | 0.256137633 | 53 |
| V-ATPase_C | 0.25620473 | 56 |
| Svf1 | 0.256938181 | 50 |
| polyprenyl_synt | 0.257798971 | 56 |
| Nup84_Nup100 | 0.257937239 | 56 |
| Ribosomal_L1 | 0.258069889 | 56 |
| KH_1 | 0.258465308 | 56 |
| Vps54 | 0.258758292 | 53 |
| HAD_2 | 0.258861704 | 56 |
| Endosulfine | 0.258937208 | 56 |
| Met_10 | 0.259025529 | 56 |
| DUF3395 | 0.259156684 | 54 |
| ParA | 0.259655474 | 56 |
| tRNA_SAD | 0.259764721 | 56 |
| CNOT11 | 0.259790005 | 49 |
| AAA_lid_7 | 0.260192811 | 54 |
| UPF0113_N | 0.260373542 | 55 |
| Clp_N | 0.260449004 | 53 |
| SKI | 0.260775652 | 54 |
| UN_NPL4 | 0.261579172 | 46 |
| Thymidylate_kin | 0.262281762 | 55 |
| ETF_alpha | 0.263238334 | 56 |
| IF3_N | 0.263368774 | 48 |

| | | |
|---|---|---|
| Tim17 | 0.263404826 | 56 |
| PRMT5_C | 0.263589988 | 53 |
| PRKCSH_1 | 0.263746172 | 51 |
| Spore_permease | 0.263850924 | 22 |
| Choline_transpo | 0.264507194 | 56 |
| DUF1768 | 0.265730526 | 20 |
| Pescadillo_N | 0.265982636 | 55 |
| UPF0047 | 0.266332892 | 52 |
| NOP5NT | 0.266900335 | 56 |
| Cpn60_TCP1 | 0.267921501 | 56 |
| TFIIF_beta_N | 0.268639825 | 54 |
| HMGL-like | 0.268871807 | 56 |
| Dynactin_p62 | 0.269102917 | 55 |
| DUF1751 | 0.269251966 | 51 |
| MIR | 0.269747291 | 56 |
| XPC-binding | 0.269752509 | 55 |
| Ribosomal_S18 | 0.270835331 | 53 |
| Ldh_1_N | 0.271198121 | 56 |
| ARD | 0.27162545 | 54 |
| OHCU_decarbox | 0.271628518 | 50 |
| CTNNBL | 0.271894183 | 49 |
| PCI | 0.272210105 | 56 |
| Pribosyltran | 0.27363705 | 56 |
| Peptidase_C26 | 0.273755916 | 56 |
| VMA21 | 0.274320169 | 43 |
| MutS_I | 0.274826528 | 56 |
| Prot_ATP_ID_OB | 0.27490124 | 56 |
| GSH_synth_ATP | 0.275006625 | 53 |
| Rab3-GTPase_cat | 0.275133263 | 50 |
| TIP49_C | 0.275216075 | 56 |
| TOM13 | 0.276301151 | 51 |
| SPT_ssu-like | 0.277523303 | 43 |
| Glyco_transf_41 | 0.277592163 | 52 |
| MCM_bind | 0.277649769 | 53 |
| RICTOR_N | 0.279563425 | 55 |
| 2Fe-2S_thioredx | 0.279899237 | 56 |
| DAO_C | 0.280065509 | 55 |
| Abhydrolase_1 | 0.280164554 | 56 |
| Per1 | 0.280400998 | 53 |
| Ribosomal_L12 | 0.280781928 | 54 |
| Mak16 | 0.280991337 | 55 |
| Cnd1_N | 0.281156212 | 54 |
| RED_N | 0.281561527 | 53 |
| DUF3591 | 0.282913688 | 56 |
| zf-C3HC4_4 | 0.282934691 | 41 |
| NUFIP1 | 0.283039858 | 54 |
| Glyco_hydro38C2 | 0.2835583 | 54 |
| DUF1754 | 0.28416281 | 23 |
| E1_dh | 0.28434124 | 56 |
| PPTA | 0.284385729 | 56 |
| Ribophorin_I | 0.285944088 | 54 |
| Leo1 | 0.286220867 | 53 |
| Histidinol_dh | 0.287136783 | 55 |
| PWI | 0.287170089 | 55 |
| ANAPC10 | 0.287686962 | 55 |
| eRF1_2 | 0.288111548 | 56 |
| zf-ZPR1 | 0.288502189 | 51 |
| IFRD | 0.288649901 | 54 |
| ABC_tran | 0.288721173 | 56 |
| Peptidase_M17 | 0.289339196 | 56 |
| Fructosamin_kin | 0.289506053 | 30 |
| G10 | 0.289790537 | 54 |
| Ufd2P_core | 0.290333878 | 55 |
| MoaC | 0.290336067 | 51 |
| TMA7 | 0.29045992 | 43 |
| NOG1_N | 0.290650951 | 55 |
| Rrp40_N | 0.291125029 | 51 |
| UMP1 | 0.291161197 | 56 |
| ETC_C1_NDUFA4 | 0.291225597 | 54 |
| Glyco_transf_4 | 0.291335776 | 56 |
| Met_synt_B12 | 0.291478345 | 44 |
| RRM_Rrp7 | 0.291560131 | 48 |
| Mannosyl_trans | 0.292042866 | 52 |
| LUC7 | 0.292079659 | 54 |
| Syntaxin-18_N | 0.292199668 | 47 |
| RuvB_N | 0.292744178 | 56 |
| CX9C | 0.292913383 | 53 |
| Med22 | 0.292954659 | 52 |
| Trehalase | 0.293174882 | 56 |
| NtCtMGAM_N | 0.294274937 | 40 |
| Snf7 | 0.294290673 | 56 |
| Calpain_III | 0.29493851 | 39 |
| AAA_lid_3 | 0.295428078 | 56 |
| UCR_14kD | 0.295469123 | 53 |
| Rep-A_N | 0.295561512 | 52 |
| CorA | 0.295719671 | 56 |
| E3_binding | 0.296039446 | 56 |
| Syntaxin-6_N | 0.296152549 | 52 |
| Glyco_hydro_47 | 0.296266142 | 56 |
| SURF6 | 0.29638323 | 54 |
| DASH_Dam1 | 0.296475256 | 51 |
| RTP1_C1 | 0.296586801 | 54 |
| DUF4598 | 0.296656052 | 51 |
| Aminotran_4 | 0.296935494 | 56 |
| PIG-P | 0.296980654 | 51 |
| DUF5572 | 0.297337544 | 47 |
| Fcf2 | 0.297673627 | 56 |
| DNA_pol_alpha_N | 0.297701192 | 33 |

| | | |
|---|---|---|
| zf-RING_2 | 0.297851892 | 56 |
| GCIP | 0.29920427 | 54 |
| Cpn10 | 0.299598958 | 56 |
| ChAPs | 0.299878781 | 55 |
| BBE | 0.300314158 | 30 |
| EF_TS | 0.300600192 | 53 |
| Dak2 | 0.30069368 | 46 |
| Polysacc_synt_4 | 0.301140502 | 47 |
| CPSF73-100_C | 0.302272744 | 55 |
| RNA_pol_Rpb2_4 | 0.302381852 | 56 |
| RNA_pol_Rpb2_5 | 0.302381852 | 56 |
| DEAD_2 | 0.303123717 | 56 |
| UEV | 0.30350484 | 53 |
| RNA_pol_Rpa2_4 | 0.303640604 | 55 |
| Calsequestrin | 0.303670725 | 39 |
| SRP72 | 0.304237001 | 53 |
| ATP19 | 0.304925878 | 55 |
| SRP40_C | 0.305292927 | 52 |
| tRNA-synt_2b | 0.305454517 | 56 |
| PALP | 0.305986207 | 56 |
| Rad1 | 0.306028589 | 56 |
| Nucleos_tra2_C | 0.306325376 | 53 |
| XAP5 | 0.306335387 | 53 |
| M20_dimer | 0.306527053 | 56 |
| Transcrip_reg | 0.306797556 | 42 |
| Ric8 | 0.307022481 | 55 |
| cwf18 | 0.307938945 | 53 |
| PAPA-1 | 0.308099473 | 53 |
| Fis1_TPR_N | 0.308192265 | 53 |
| Vac14_Fig4_bd | 0.308352727 | 56 |
| Peptidase_C50 | 0.308432852 | 54 |
| Peptidase_C14 | 0.308805577 | 55 |
| GN3L_Grn1 | 0.308877001 | 50 |
| Palm_thioest | 0.309113975 | 55 |
| zf-CHY | 0.309199814 | 56 |
| ATP-grasp_5 | 0.30977671 | 50 |
| SMAP | 0.309876428 | 46 |
| RPN6_N | 0.31003286 | 55 |
| SLD5_C | 0.310375511 | 45 |
| RSN1_7TM | 0.310766227 | 56 |
| TIM | 0.311432581 | 55 |
| WH2 | 0.31186867 | 54 |
| UDP-g_GGTase | 0.312273207 | 51 |
| Importin_rep_4 | 0.312663691 | 56 |
| GARS_C | 0.31306314 | 55 |
| CAF1 | 0.313338824 | 56 |
| PHM7_cyt | 0.313485287 | 56 |
| LMWPc | 0.313724879 | 54 |
| SPT6_acidic | 0.313732372 | 48 |
| QCR10 | 0.31378604 | 55 |
| RasGEF_N_2 | 0.314608057 | 52 |
| MRC1 | 0.315550261 | 41 |
| Ccdc124 | 0.315756257 | 55 |
| HAT | 0.315892748 | 55 |
| CTK3 | 0.316025627 | 49 |
| Mpv17_PMP22 | 0.316149267 | 54 |
| JmjC | 0.316336706 | 56 |
| zf-RING_UBOX | 0.316511498 | 56 |
| ANAPC8 | 0.316597901 | 48 |
| ThiG | 0.316827573 | 54 |
| PRMT5_TIM | 0.316851676 | 52 |
| LCM | 0.318380373 | 54 |
| EF_assoc_1 | 0.318676323 | 56 |
| Complex1_49kDa | 0.318998047 | 55 |
| Tam41_Mmp37 | 0.319132696 | 56 |
| PROCT | 0.319165175 | 55 |
| CBP4 | 0.319225748 | 46 |
| Aconitase | 0.319287295 | 56 |
| Aim19 | 0.319621241 | 46 |
| SQHop_cyclase_N | 0.319711755 | 55 |
| Exo84_C | 0.320722737 | 55 |
| AAA_17 | 0.320855703 | 56 |
| BPL_LplA_LipB | 0.320886822 | 56 |
| UPF0160 | 0.321054023 | 53 |
| GARS_N | 0.321114362 | 55 |
| DNA_gyraseB | 0.321233107 | 56 |
| FANCI_S2 | 0.32174422 | 51 |
| Tom22 | 0.322265797 | 55 |
| GST_N_3 | 0.32243049 | 56 |
| GATase_3 | 0.322442012 | 56 |
| TruB_N | 0.323549753 | 56 |
| Frataxin_Cyay | 0.323997153 | 55 |
| PDH | 0.32427228 | 53 |
| DASH_Ask1 | 0.324740349 | 53 |
| eRF1_3 | 0.325112492 | 56 |
| Cornichon | 0.325302561 | 54 |
| RL10P_insert | 0.325963689 | 56 |
| Nop10p | 0.325983992 | 51 |
| Mob_synth_C | 0.326381425 | 52 |
| Topoisom_I | 0.326483494 | 53 |
| DUF410 | 0.327253589 | 54 |
| TBPIP | 0.327714169 | 51 |
| Dopey_N | 0.3281559 | 52 |
| F1F0-ATPsyn_F | 0.328664853 | 54 |
| ADK_lid | 0.328851827 | 56 |
| Adenylsucc_synt | 0.32885936 | 56 |
| ATP_bind_1 | 0.329302137 | 56 |
| DUF382 | 0.329938888 | 55 |

| | | |
|---|---|---|
| PIG-H | 0.330313259 | 47 |
| Glyco_hydro_38C | 0.330636786 | 55 |
| SRP19 | 0.330652221 | 52 |
| IPPT | 0.331795534 | 55 |
| Sgf11 | 0.331945355 | 48 |
| GHMP_kinases_C | 0.332129143 | 56 |
| AAA_18 | 0.332805027 | 56 |
| Mt_ATP-synt_D | 0.333099861 | 56 |
| BHD_1 | 0.333321632 | 52 |
| TFIIE_beta | 0.333817754 | 51 |
| Nop | 0.334555646 | 56 |
| Myb_Cef | 0.334721196 | 53 |
| Pex19 | 0.334796874 | 54 |
| FRG1 | 0.335086173 | 52 |
| PINIT | 0.335542492 | 49 |
| Peptidase_M50B | 0.335659428 | 54 |
| CDC45 | 0.335732633 | 53 |
| EPSP_synthase | 0.335734493 | 55 |
| Y_phosphatase2 | 0.336009748 | 56 |
| KxDL | 0.336023264 | 50 |
| Endonuclease_NS | 0.336228082 | 56 |
| RNA_pol_Rpb1_5 | 0.336777139 | 56 |
| Meth_synt_1 | 0.336869463 | 54 |
| Beta-Casp | 0.336894688 | 56 |
| Ribosomal_L10 | 0.337000319 | 56 |
| TGT | 0.337925021 | 56 |
| UPF0086 | 0.337963529 | 51 |
| DNApol_Exo | 0.338110515 | 52 |
| RPAP2_Rtr1 | 0.338321356 | 52 |
| EIF_2_alpha | 0.338424214 | 55 |
| Proteasome_A_N | 0.33942925 | 56 |
| tRNA-synt_1g | 0.339655316 | 56 |
| ATP-synt_C | 0.339725519 | 56 |
| MSC | 0.340109565 | 52 |
| Vfa1 | 0.340191625 | 47 |
| AARP2CN | 0.341327181 | 53 |
| Ribosomal_L12_N | 0.341349008 | 55 |
| TB2_DP1_HVA22 | 0.341492151 | 56 |
| OPT | 0.341807579 | 56 |
| UCR_hinge | 0.341808944 | 52 |
| APH | 0.342267492 | 56 |
| ARS2 | 0.342307425 | 55 |
| Peptidase_M18 | 0.342370255 | 56 |
| Hep_59 | 0.342623297 | 53 |
| Sec20 | 0.342960479 | 54 |
| MCM6_C | 0.34307849 | 48 |
| Zn_clus | 0.343390789 | 56 |
| C1_4 | 0.34345031 | 52 |
| HTH_Tnp_Tc5 | 0.343995449 | 22 |
| Rcd1 | 0.34435526 | 55 |
| RNA_pol_Rpb2_1 | 0.3446194 | 56 |
| Mak10 | 0.345391472 | 54 |
| Mito_fiss_Elm1 | 0.345501426 | 49 |
| zinc_ribbon_9 | 0.345513315 | 29 |
| Kin17_mid | 0.345782425 | 56 |
| EFG_C | 0.346163744 | 56 |
| DASH_Hsk3 | 0.346421703 | 25 |
| COG6 | 0.346488447 | 53 |
| Drc1-Sld2 | 0.346755172 | 31 |
| AAA_33 | 0.347165047 | 56 |
| PH_BEACH | 0.347885456 | 55 |
| Exo70 | 0.348771063 | 54 |
| V-ATPase_G | 0.34885324 | 52 |
| Syja_N | 0.348892759 | 56 |
| Maf1 | 0.349019702 | 55 |
| Calcipressin | 0.349300183 | 55 |
| Fip1 | 0.349452558 | 54 |
| PRA-CH | 0.349732132 | 54 |
| Yos1 | 0.349794917 | 49 |
| Guanylate_kin | 0.350018425 | 54 |
| EAF | 0.350188403 | 39 |
| SAGA-Tad1 | 0.351126776 | 55 |
| JTB | 0.351291408 | 40 |
| TIP120 | 0.351489036 | 54 |
| GNAT_acetyltr_2 | 0.351583194 | 55 |
| 5_nucleotid | 0.351967152 | 40 |
| HTH_44 | 0.352004391 | 51 |
| OTU | 0.352019028 | 54 |
| Coatomer_WDAD | 0.352318053 | 56 |
| Macro | 0.352874896 | 44 |
| GST_C | 0.35295907 | 56 |
| SIS | 0.353245024 | 56 |
| OSCP | 0.353312064 | 56 |
| RIBIOP_C | 0.354876339 | 54 |
| ATP_transf | 0.355096382 | 51 |
| Prenyltrans | 0.35509857 | 56 |
| BCIP | 0.355128908 | 56 |
| Proteasom_PSMB | 0.355417215 | 55 |
| APG5 | 0.355649666 | 55 |
| zf-CCCH_4 | 0.356072754 | 56 |
| SMC_Nse1 | 0.356658183 | 55 |
| Tfb5 | 0.356933699 | 52 |
| MPC | 0.357019883 | 56 |
| Rgp1 | 0.357025695 | 55 |
| UPF0203 | 0.357647067 | 49 |
| Eapp_C | 0.357744728 | 47 |
| Ham1p_like | 0.358100197 | 55 |
| Nucleos_tra2_N | 0.358791861 | 52 |

| | | |
|---|---|---|
| CPSF_A | 0.35887541 | 56 |
| ACPS | 0.359260054 | 56 |
| BolA | 0.360242897 | 56 |
| SRP14 | 0.360872349 | 47 |
| RPN6_C_helix | 0.360880254 | 55 |
| Vps8 | 0.361114562 | 55 |
| Sortilin_C | 0.361300425 | 55 |
| MAM33 | 0.361534248 | 56 |
| Ribul_P_3_epim | 0.361596849 | 55 |
| ATG7_N | 0.361931019 | 51 |
| GST_C_2 | 0.36198075 | 56 |
| Mitoc_mL59 | 0.362877241 | 53 |
| DUF3752 | 0.363222017 | 51 |
| Fis1_TPR_C | 0.363230252 | 54 |
| MitMem_reg | 0.363233421 | 56 |
| WLM | 0.363244389 | 56 |
| Maf | 0.363285982 | 54 |
| CPBP | 0.363370033 | 50 |
| Gaa1 | 0.363425817 | 53 |
| Complex1_30kDa | 0.364112261 | 55 |
| CactinC_cactus | 0.364455628 | 55 |
| Pmp3 | 0.366236648 | 56 |
| DAHP_synth_2 | 0.366465121 | 55 |
| ATP-synt_J | 0.367214996 | 49 |
| PIGA | 0.367260175 | 55 |
| QRPTase_C | 0.367439721 | 51 |
| PRO8NT | 0.367443741 | 55 |
| RPN7 | 0.36753199 | 56 |
| GSH_synthase | 0.368040977 | 53 |
| RICTOR_V | 0.368175718 | 53 |
| Hat1_N | 0.368429387 | 53 |
| Mpp10 | 0.369586671 | 53 |
| GST_C_3 | 0.370612992 | 56 |
| ribosomal_L24 | 0.370633416 | 53 |
| Vps16_N | 0.370785457 | 56 |
| AMMECR1 | 0.370890875 | 51 |
| ATP-synt_Eps | 0.370984396 | 52 |
| RNA_pol_N | 0.371102973 | 54 |
| Fer4_12 | 0.37190594 | 48 |
| Ldh_1_C | 0.372128912 | 56 |
| Cellulase | 0.3727103 | 55 |
| ALO | 0.373483634 | 54 |
| NuiA | 0.374034282 | 35 |
| Rpr2 | 0.374165411 | 52 |
| Longin | 0.375272118 | 56 |
| Ipi1_N | 0.375301807 | 52 |
| Elf1 | 0.375785424 | 53 |
| ALS_ss_C | 0.376058137 | 55 |
| Hira | 0.376160553 | 52 |
| Voldacs | 0.376628539 | 54 |
| REPA_OB_2 | 0.37688768 | 54 |
| zf-C3HC4 | 0.377260011 | 56 |
| Paf67 | 0.378433757 | 53 |
| DUF4217 | 0.379100017 | 56 |
| Complex1_LYR | 0.379266595 | 56 |
| Peptidase_C54 | 0.379463431 | 56 |
| AAA_21 | 0.379908168 | 56 |
| TFIID-18kDa | 0.38077541 | 56 |
| RSN1_TM | 0.380809406 | 56 |
| MRP-S33 | 0.381541249 | 53 |
| Cgr1 | 0.381705276 | 52 |
| Barwin | 0.382083461 | 51 |
| BHD_3 | 0.38247799 | 55 |
| DHHA1 | 0.38251217 | 54 |
| Chorismate_bind | 0.382782351 | 56 |
| TOPRIM_C | 0.383476943 | 56 |
| Dabb | 0.384050414 | 45 |
| CDC37_C | 0.384674716 | 54 |
| tRNA_anti-codon | 0.384823613 | 56 |
| Tim54 | 0.385122072 | 56 |
| Cnd3 | 0.385451219 | 53 |
| Mgm101p | 0.385928085 | 54 |
| URO-D | 0.386287161 | 56 |
| 4F5 | 0.38660787 | 56 |
| BRF1 | 0.388191532 | 53 |
| V-SNARE_C | 0.388207788 | 56 |
| Cofilin_ADF | 0.388326677 | 56 |
| Zip | 0.388612125 | 56 |
| Peptidase_M22 | 0.388801719 | 56 |
| Anth_synt_I_N | 0.389265247 | 55 |
| AhpC-TSA | 0.389335316 | 56 |
| RMMBL | 0.389350534 | 56 |
| AATF-Che1 | 0.390257782 | 51 |
| ORC3_N | 0.390591347 | 56 |
| Ist1 | 0.390697813 | 54 |
| PAP_assoc | 0.391033545 | 56 |
| DBR1 | 0.391097135 | 54 |
| Pet100 | 0.391296756 | 53 |
| YEATS | 0.391578024 | 55 |
| DDOST_48kD | 0.391911627 | 55 |
| RNA_pol_Rpb2_2 | 0.391913876 | 56 |
| YqgF | 0.391946792 | 51 |
| Rotamase_3 | 0.392015444 | 55 |
| Rotamase | 0.392015444 | 55 |
| RNA_pol_Rpb1_4 | 0.392190485 | 56 |
| ERO1 | 0.392574438 | 56 |
| MFAP1 | 0.393416375 | 52 |
| NDUFA12 | 0.393431925 | 56 |

| | | |
|---|---|---|
| RNA_pol_Rpc34 | 0.393512327 | 56 |
| Rsm22 | 0.393654491 | 56 |
| Sec7_N | 0.394715938 | 56 |
| Upf2 | 0.394888875 | 53 |
| PRMT5 | 0.395016165 | 53 |
| ChaC | 0.395419907 | 56 |
| RNA_pol_L_2 | 0.395519683 | 56 |
| GFA | 0.396039715 | 43 |
| zf-U1 | 0.397155827 | 53 |
| Methyltransf_PK | 0.397415026 | 55 |
| RNA_pol_Rpb2_3 | 0.397694397 | 56 |
| UPF0029 | 0.398081276 | 53 |
| DHHA2 | 0.398236066 | 53 |
| IGPS | 0.400121768 | 55 |
| MMR_HSR1_Xtn | 0.400270739 | 56 |
| ThiS | 0.400803538 | 45 |
| RNase_H_2 | 0.40085223 | 43 |
| Glyco_hydro_63N | 0.400886152 | 54 |
| PDZ_1 | 0.401261632 | 55 |
| ETC_C1_NDUFA5 | 0.401357267 | 56 |
| HRDC | 0.401800006 | 53 |
| UTP15_C | 0.401820996 | 54 |
| A_deamin | 0.402512447 | 53 |
| PRP38 | 0.402528894 | 56 |
| Tmemb_14 | 0.40344231 | 49 |
| Auto_anti-p27 | 0.404047089 | 55 |
| LepA_C | 0.404172771 | 51 |
| TGS | 0.40438652 | 56 |
| Rbsn | 0.404440805 | 50 |
| Aminotran_5 | 0.40454735 | 56 |
| ATP-grasp | 0.404635704 | 56 |
| Abhydrolase_6 | 0.40467922 | 56 |
| DUF1726 | 0.40475342 | 55 |
| Atx10homo_assoc | 0.405093306 | 54 |
| HTH_9 | 0.405210931 | 44 |
| ARPC4 | 0.405432513 | 55 |
| Tom7 | 0.405507833 | 37 |
| TRAPP | 0.406006899 | 56 |
| DBINO | 0.406234293 | 54 |
| Acyltransf_C | 0.406626866 | 56 |
| BLACT_WH | 0.406962418 | 49 |
| KH_6 | 0.407433312 | 56 |
| ERp29 | 0.408551251 | 51 |
| COX6B | 0.408714207 | 55 |
| Hydrolase_4 | 0.408803617 | 56 |
| FBPase | 0.408823482 | 54 |
| IF-2B | 0.409134927 | 56 |
| zf-met2 | 0.409434753 | 51 |
| Sin_N | 0.411305481 | 53 |
| Sec62 | 0.411744353 | 55 |
| SYS1 | 0.412042179 | 50 |
| ELFV_dehydrog | 0.412151484 | 56 |
| EFTUD2 | 0.412393285 | 56 |
| RsfS | 0.412696721 | 56 |
| Nucleoporin2 | 0.412763462 | 49 |
| Vps53_N | 0.4131141 | 55 |
| PRP21_like_P | 0.413278815 | 55 |
| zf-HIT | 0.413452407 | 53 |
| Pam16 | 0.414725324 | 56 |
| RNA_pol_Rpb1_1 | 0.415024974 | 56 |
| Utp14 | 0.415438394 | 55 |
| RRP7 | 0.415722754 | 54 |
| Lum_binding | 0.416251207 | 56 |
| GDPD | 0.417130747 | 56 |
| Pantoate_transf | 0.417461293 | 56 |
| Ribosomal_S17 | 0.417478649 | 56 |
| ALAD | 0.417706794 | 55 |
| TAXi_C | 0.418235536 | 33 |
| TFIIA_gamma_C | 0.418526295 | 52 |
| ThiF | 0.418551019 | 56 |
| DBP10CT | 0.419731383 | 52 |
| SHS2_Rpb7-N | 0.419858501 | 56 |
| eIF2_C | 0.419920382 | 55 |
| THOC2_N | 0.420849865 | 54 |
| ABC_membrane | 0.421431721 | 56 |
| AAA_23 | 0.422003723 | 56 |
| Diphthamide_syn | 0.422028862 | 56 |
| CBM_19 | 0.422112662 | 51 |
| Spc24 | 0.423201454 | 37 |
| DUF1168 | 0.423363302 | 54 |
| BAH | 0.423395197 | 53 |
| MINDY_DUB | 0.4237132 | 51 |
| FGAR-AT_linker | 0.424463003 | 55 |
| Cdc6_C | 0.42462575 | 43 |
| Proteasome | 0.425010905 | 56 |
| dsrm | 0.425116279 | 55 |
| RNA_pol_Rpb1_3 | 0.425261127 | 56 |
| LSM | 0.425373847 | 56 |
| ALG11_N | 0.425964184 | 56 |
| TRAPPC9-Trs120 | 0.426035849 | 53 |
| PDCD2_C | 0.426561109 | 56 |
| DNA_topoisoIV | 0.426627948 | 56 |
| DUF2012 | 0.427320658 | 54 |
| MAD | 0.42759938 | 42 |
| CDC37_M | 0.427961939 | 55 |
| Cu-oxidase_3 | 0.429931736 | 54 |
| Ribosomal_L9_N | 0.430534424 | 52 |
| SSF | 0.430704889 | 47 |

| | | |
|---|---|---|
| RINT1_TIP1 | 0.430719587 | 53 |
| SLBB | 0.430877315 | 52 |
| MutS_V | 0.431483557 | 56 |
| FLILHELTA | 0.431664721 | 51 |
| DER1 | 0.431953292 | 55 |
| Ribosomal_L36 | 0.43206415 | 47 |
| Sortilin-Vps10 | 0.432389151 | 55 |
| Mad3_BUB1_I | 0.4328025 | 54 |
| NOB1_Zn_bind | 0.432866401 | 56 |
| TFA2_Winged_2 | 0.43319887 | 53 |
| OPA3 | 0.433424385 | 53 |
| Glyco_hydro_38N | 0.433524069 | 55 |
| zf-CSL | 0.433985629 | 55 |
| Synaptobrevin | 0.434070789 | 56 |
| PGM_PMM_IV | 0.434234766 | 56 |
| ArsA_HSP20 | 0.434862407 | 43 |
| GCN5L1 | 0.435444316 | 52 |
| AAA_15 | 0.435574078 | 55 |
| PRCC | 0.435755292 | 51 |
| RPA43_OB | 0.435774505 | 55 |
| Flavoprotein | 0.435866833 | 51 |
| Peptidase_C13 | 0.435915927 | 54 |
| Suf | 0.435990528 | 56 |
| DUF3546 | 0.436143484 | 55 |
| Gar1 | 0.436225615 | 56 |
| BING4CT | 0.43639028 | 54 |
| Suv3_C_1 | 0.43647707 | 53 |
| Hydantoinase_A | 0.436512805 | 54 |
| Lactamase_B_6 | 0.437178423 | 56 |
| SHNi-TPR | 0.437728593 | 51 |
| Utp21 | 0.438113759 | 55 |
| Peptidase_M50 | 0.438984717 | 48 |
| Img2 | 0.439703394 | 53 |
| TatD_DNase | 0.439990095 | 55 |
| ACC_central | 0.440107107 | 55 |
| RWD | 0.440302916 | 54 |
| Sec16_C | 0.440471522 | 52 |
| eIF3_N | 0.440741636 | 55 |
| ACT_5 | 0.440918538 | 53 |
| S-AdoMet_synt_C | 0.440937297 | 55 |
| tRNA-synt_1e | 0.441328651 | 56 |
| RNA_pol_Rbc25 | 0.44134473 | 55 |
| P34-Arc | 0.441640588 | 54 |
| Sec34 | 0.441673741 | 53 |
| FANCI_HD1 | 0.441883065 | 46 |
| CIAPIN1 | 0.441929622 | 55 |
| NUDIX_2 | 0.442390244 | 33 |
| Glyco_transf_22 | 0.443140473 | 56 |
| AAA_lid_9 | 0.443468473 | 54 |
| MoaE | 0.443845218 | 52 |
| Sdh_cyt | 0.4438604 | 55 |
| MCM2_N | 0.443886942 | 50 |
| Spb1_C | 0.444576407 | 56 |
| Rep_fac-A_C | 0.444727191 | 52 |
| NADH_4Fe-4S | 0.445150556 | 52 |
| FACT-Spt16_Nlob | 0.445520111 | 55 |
| SPT16 | 0.445520111 | 55 |
| DUF4203 | 0.445548865 | 56 |
| ORC4_C | 0.445692852 | 55 |
| FANCL_C | 0.445750976 | 51 |
| Thioesterase | 0.447087101 | 51 |
| TIMELESS | 0.447297412 | 54 |
| COX16 | 0.44744211 | 51 |
| PMM | 0.447606119 | 54 |
| Methyltransf_34 | 0.447731645 | 48 |
| DUF4499 | 0.447860581 | 41 |
| RPAP1_C | 0.448437962 | 54 |
| PRA-PH | 0.448886509 | 55 |
| U5_2-snRNA_bdg | 0.448906888 | 54 |
| PGK | 0.44926142 | 55 |
| ThrE_2 | 0.449346665 | 53 |
| DAD | 0.449390473 | 55 |
| Npa1 | 0.449778925 | 52 |
| Kri1 | 0.449954532 | 54 |
| Topo_C_assoc | 0.450093839 | 53 |
| Brix | 0.450775935 | 56 |
| Prenyltransf | 0.451250366 | 55 |
| Sec63 | 0.452219006 | 56 |
| SART-1 | 0.452523501 | 53 |
| CRIM | 0.453342333 | 53 |
| CTU2 | 0.454058401 | 55 |
| Thioredoxin | 0.454131489 | 56 |
| QRPTase_N | 0.454212951 | 51 |
| SMC_hinge | 0.454420908 | 56 |
| OGG_N | 0.455037036 | 48 |
| TPK_catalytic | 0.455148631 | 53 |
| Arginosuc_synth | 0.45517292 | 55 |
| DUF4078 | 0.455612217 | 53 |
| RTC_insert | 0.456072344 | 53 |
| PHF5 | 0.456097464 | 53 |
| DUF543 | 0.456648486 | 53 |
| Actin | 0.457341168 | 56 |
| Isy1 | 0.457775422 | 55 |
| Ribosomal_L19 | 0.458553239 | 56 |
| zf-CHCC | 0.458641426 | 54 |
| Rtf2 | 0.458862733 | 54 |
| COX15-CtaA | 0.458975569 | 55 |
| BIR | 0.459011915 | 43 |

| | | |
|---|---|---|
| Ribosomal_S6 | 0.459038775 | 55 |
| CPL | 0.459328293 | 56 |
| CBF | 0.459525987 | 56 |
| MCM_N | 0.459568447 | 56 |
| PAP2 | 0.459745944 | 56 |
| ABC_tran_Xtn | 0.460015738 | 56 |
| CDC73_C | 0.460170338 | 53 |
| SHQ1 | 0.460488604 | 53 |
| CM_2 | 0.460539334 | 48 |
| D123 | 0.460757284 | 56 |
| Steroid_dh | 0.46106879 | 56 |
| zf-C3HC4_2 | 0.461092158 | 56 |
| CLP1_N | 0.461215269 | 53 |
| Sacchrp_dh_C | 0.461401316 | 56 |
| NDUF_B7 | 0.46199998 | 56 |
| Peptidase_C2 | 0.462404939 | 54 |
| Ribosomal_L17 | 0.462438668 | 56 |
| Csm1 | 0.462586652 | 41 |
| Radical_SAM_C | 0.462665287 | 56 |
| Tma16 | 0.462670016 | 54 |
| RIC1 | 0.462773496 | 52 |
| Tim44 | 0.462816352 | 56 |
| GTP_cyclohydrol | 0.462919862 | 56 |
| TPK_B1_binding | 0.46305645 | 54 |
| LAMTOR5 | 0.463405395 | 47 |
| Chorismate_synt | 0.463446787 | 56 |
| tRNA_bind_2 | 0.463530012 | 55 |
| AAA_lid_10 | 0.464043194 | 54 |
| DcpS | 0.464048834 | 53 |
| ORC5_C | 0.464544142 | 54 |
| Hexapep | 0.466218314 | 56 |
| S-AdoMet_synt_N | 0.466294951 | 56 |
| S-AdoMet_synt_M | 0.466294951 | 56 |
| OxoGdeHyase_C | 0.466344784 | 56 |
| DUF1764 | 0.466348806 | 47 |
| dCMP_cyt_deam_1 | 0.466540191 | 56 |
| ProRS-C_1 | 0.46671071 | 55 |
| AMP-binding | 0.466930518 | 56 |
| FAS_N | 0.466931019 | 46 |
| HAUS6_N | 0.467046832 | 43 |
| Rogdi_lz | 0.467226165 | 44 |
| SF3b1 | 0.467641867 | 51 |
| F-actin_cap_A | 0.46822504 | 53 |
| TRAPPC-Trs85 | 0.468495 | 53 |
| Mhr1 | 0.468564773 | 48 |
| Perilipin | 0.468676415 | 25 |
| Afi1 | 0.468919788 | 55 |
| GATase_4 | 0.468958052 | 56 |
| 2OG-FeII_Oxy_2 | 0.469648372 | 56 |
| DHO_dh | 0.470276213 | 56 |
| DUF2428 | 0.470813391 | 53 |
| MRP-L20 | 0.470838139 | 54 |
| COG4 | 0.471053291 | 55 |
| CLU_N | 0.471101453 | 56 |
| Prok-RING_4 | 0.471941642 | 53 |
| Ssl1 | 0.472707289 | 55 |
| Rad21_Rec8 | 0.473138106 | 49 |
| TFCD_C | 0.473512883 | 54 |
| CLU | 0.473999333 | 56 |
| L31 | 0.474063131 | 56 |
| Complex1_51K | 0.474138187 | 53 |
| RICTOR_M | 0.474296264 | 54 |
| Pro_isomerase | 0.474680377 | 56 |
| WD-3 | 0.475750113 | 51 |
| RNA_pol_Rpb1_7 | 0.475916879 | 52 |
| Rad10 | 0.475930088 | 54 |
| Sdh5 | 0.476183505 | 53 |
| tRNA-synt_1_2 | 0.476325948 | 56 |
| DUF866 | 0.476494073 | 53 |
| RPN2_C | 0.476804061 | 56 |
| Utp13 | 0.476833466 | 53 |
| AHSA1 | 0.477269613 | 56 |
| XRN_N | 0.477270267 | 53 |
| SF3A3 | 0.477449316 | 55 |
| MAGE | 0.477903407 | 53 |
| Autophagy_N | 0.478098042 | 56 |
| PIG-Y | 0.478321709 | 48 |
| SF3A2 | 0.478540295 | 53 |
| RPN1_RPN2_N | 0.478689401 | 56 |
| ATP-synt_10 | 0.479042665 | 51 |
| tRNA-synt_2 | 0.479292176 | 56 |
| RNA_pol_L | 0.479820705 | 56 |
| Pr_beta_C | 0.480214105 | 54 |
| DUF2039 | 0.480588561 | 55 |
| U3_assoc_6 | 0.480648976 | 51 |
| RPN1_C | 0.48090105 | 56 |
| Myb_DNA-bind_7 | 0.481067037 | 53 |
| dsDNA_bind | 0.481324953 | 56 |
| Sde2_N_Ubi | 0.481395813 | 53 |
| 60KD_IMP | 0.481461263 | 55 |
| Kri1_C | 0.481828719 | 55 |
| AAA_29 | 0.482045205 | 39 |
| AAA_9 | 0.482220026 | 56 |
| Wbp11 | 0.482531115 | 52 |
| Nse4_C | 0.482890052 | 54 |
| PRP8_domainIV | 0.483265734 | 55 |
| 3-HAO | 0.483463824 | 54 |
| GLTSCR1 | 0.483647431 | 52 |

| | | |
|---|---|---|
| PET117 | 0.48376092 | 51 |
| Nipped-B_C | 0.484143312 | 55 |
| TRM | 0.484443442 | 56 |
| Hydantoinase_B | 0.484725992 | 54 |
| GDA1_CD39 | 0.484745458 | 56 |
| IF-2 | 0.484952427 | 56 |
| DUF1298 | 0.486126181 | 23 |
| EFP | 0.486179657 | 52 |
| Prefoldin_2 | 0.4862878 | 56 |
| SIR2 | 0.48633833 | 56 |
| Rpn3_C | 0.486502095 | 54 |
| eIF-3c_N | 0.487854514 | 55 |
| MRP-L46 | 0.48796337 | 55 |
| Phosducin | 0.487970161 | 56 |
| Hydant_A_N | 0.488270495 | 54 |
| NuA4 | 0.488301746 | 53 |
| FGAR-AT_N | 0.488630961 | 55 |
| Elong-fact-P_C | 0.488767404 | 52 |
| Evr1_Alr | 0.489238992 | 56 |
| MRP_L53 | 0.489277916 | 49 |
| WGG | 0.490195794 | 56 |
| fn3_2 | 0.49067552 | 55 |
| NUC130_3NT | 0.490782213 | 55 |
| TRAUB | 0.4908301 | 52 |
| Glyco_hydro_63 | 0.491310935 | 56 |
| ASL_C2 | 0.491435969 | 55 |
| G_glu_transpept | 0.491772415 | 55 |
| SKIP_SNW | 0.491962563 | 56 |
| FAA_hydrolase_N | 0.492195622 | 55 |
| FAM72 | 0.493334628 | 54 |
| UCR_UQCRX_QCR9 | 0.493566163 | 53 |
| Ribosomal_L21p | 0.493611052 | 55 |
| Med21 | 0.493878323 | 55 |
| Citrate_synt | 0.493905761 | 56 |
| RNA_pol_Rpb4 | 0.49398222 | 56 |
| DUF1674 | 0.494060482 | 52 |
| zn-ribbon_14 | 0.49440878 | 54 |
| RNase_PH | 0.494652318 | 56 |
| tRNA-synt_1b | 0.495206333 | 56 |
| DRIM | 0.495291968 | 54 |
| SH3_12 | 0.49543189 | 42 |
| UBA_e1_thiolCys | 0.495704815 | 56 |
| CDO_I | 0.495933598 | 52 |
| TFIIA_gamma_N | 0.495988612 | 53 |
| Fcf1 | 0.496111916 | 55 |
| DNA_pol3_delta2 | 0.496260517 | 56 |
| SRP_SPB | 0.496362947 | 54 |
| PRKCSH | 0.496605538 | 56 |
| Shikimate_dh_N | 0.497060288 | 54 |
| Polyketide_cyc2 | 0.497067055 | 32 |
| JmjN | 0.497409027 | 56 |
| Thioredoxin_2 | 0.497639007 | 48 |
| zf-C3HC | 0.498993996 | 47 |
| GRIM-19 | 0.499366497 | 56 |
| SAICAR_synt | 0.499783853 | 55 |
| DUF498 | 0.499964003 | 55 |
| RPA_C | 0.5003033 | 48 |
| Urm1 | 0.500535652 | 55 |
| Dynein_heavy | 0.501005646 | 55 |
| ArgJ | 0.501529275 | 54 |
| zf-RING_11 | 0.501720097 | 56 |
| YIF1 | 0.501731013 | 53 |
| PAXNEB | 0.501995354 | 56 |
| Alg6_Alg8 | 0.502176626 | 56 |
| Dak1 | 0.503237674 | 47 |
| tRNA_synt_1c_R2 | 0.503574763 | 55 |
| RPOL_N | 0.503679362 | 55 |
| Nop25 | 0.503699293 | 49 |
| DUF4477 | 0.503981303 | 40 |
| PEX-1N | 0.504007636 | 55 |
| EXOSC1 | 0.505825592 | 55 |
| Ribo_biogen_C | 0.50594065 | 56 |
| TIP41 | 0.505986447 | 53 |
| PMC2NT | 0.506175157 | 51 |
| ECR1_N | 0.506280711 | 56 |
| Homeobox_KN | 0.506401523 | 56 |
| DHBP_synthase | 0.506717998 | 55 |
| THOC7 | 0.507005724 | 50 |
| LETM1 | 0.507203625 | 56 |
| MRP-L47 | 0.507278843 | 53 |
| DUF4211 | 0.507495638 | 34 |
| PCNA_C | 0.508028094 | 56 |
| Ten1_2 | 0.508113434 | 38 |
| CRCB | 0.508116004 | 53 |
| DUF3449 | 0.508310476 | 55 |
| HORMA | 0.508714548 | 54 |
| Erf4 | 0.508798342 | 51 |
| DUF383 | 0.508952445 | 55 |
| DSS1_SEM1 | 0.50942129 | 54 |
| NOC3p | 0.511381935 | 55 |
| UPF0139 | 0.511682858 | 52 |
| BOP1NT | 0.512271378 | 54 |
| Rad9 | 0.512357992 | 56 |
| SYF2 | 0.512685529 | 54 |
| IGR | 0.512804388 | 47 |
| MMM1 | 0.512885963 | 54 |
| UbiA | 0.513350618 | 56 |
| Methyltr_RsmF_N | 0.513548347 | 49 |

| | | |
|---|---|---|
| DASH_Dad3 | 0.514247831 | 42 |
| Pantoate_ligase | 0.514520285 | 55 |
| Thioredoxin_12 | 0.5155546 | 53 |
| Thioredoxin_13 | 0.5155546 | 53 |
| PDDEXK_1 | 0.515974115 | 22 |
| SAD_SRA | 0.516351493 | 34 |
| SPC22 | 0.516702077 | 52 |
| GATase_5 | 0.516942604 | 55 |
| Lysine_decarbox | 0.517114102 | 51 |
| DUF846 | 0.517116304 | 55 |
| NFACT-R_1 | 0.517195208 | 55 |
| Fe-ADH_2 | 0.517198083 | 53 |
| Cnd2 | 0.517962636 | 55 |
| ATP-synt_ab_Xtn | 0.517975203 | 56 |
| CAP_N | 0.518176016 | 54 |
| GMP_synt_C | 0.518590055 | 54 |
| HCNGP | 0.519142101 | 51 |
| Pex2_Pex12 | 0.519629829 | 56 |
| RNA_pol_Rpb1_6 | 0.519683714 | 54 |
| DUF3535 | 0.52047675 | 56 |
| Glyoxalase | 0.521273492 | 56 |
| Mem_trans | 0.521391027 | 54 |
| hGDE_central | 0.521928249 | 54 |
| Prp31_C | 0.522086242 | 56 |
| HEM4 | 0.522239548 | 50 |
| Peptidase_M1 | 0.522253315 | 56 |
| FRB_dom | 0.522538526 | 56 |
| PheRS_DBD3 | 0.52254175 | 56 |
| ATP12 | 0.52254175 | 56 |
| ORC2 | 0.522855298 | 54 |
| PGI | 0.52341751 | 56 |
| NAT | 0.523855963 | 56 |
| RibD_C | 0.524423388 | 52 |
| RNA_pol_3_Rpc31 | 0.524659068 | 53 |
| PRP4 | 0.524664936 | 54 |
| Thoc2 | 0.524840996 | 54 |
| Fumble | 0.525028235 | 56 |
| Thr_dehydrat_C | 0.525176118 | 55 |
| Bud13 | 0.525209207 | 55 |
| AD | 0.525405659 | 50 |
| NIF3 | 0.525587298 | 54 |
| Inositol_P | 0.525767191 | 56 |
| tRNA_Me_trans | 0.525834081 | 55 |
| 2OG-FeII_Oxy_3 | 0.525907396 | 56 |
| DUF1682 | 0.526009311 | 55 |
| Methyltransf_10 | 0.526212273 | 51 |
| BRCA-2_OB1 | 0.526324978 | 51 |
| FANCI_S1 | 0.526360073 | 51 |
| MIS13 | 0.526691771 | 53 |
| MoeA_N | 0.527292998 | 53 |
| Tubulin_3 | 0.528352295 | 55 |
| Sybindin | 0.528653227 | 56 |
| DUF747 | 0.529288019 | 56 |
| EAP30 | 0.529971539 | 56 |
| zf-C2H2_4 | 0.530081008 | 55 |
| MAT1 | 0.531145932 | 56 |
| Inos-1-P_synth | 0.531394249 | 55 |
| SPC25 | 0.532617157 | 53 |
| Las1 | 0.532735476 | 53 |
| Topoisom_I_N | 0.532803986 | 53 |
| GDE_C | 0.533171874 | 54 |
| DCP2 | 0.533186443 | 48 |
| Sec10 | 0.533209755 | 56 |
| Med10 | 0.533287847 | 55 |
| RNA_pol_Rpb1_2 | 0.533331445 | 56 |
| AICARFT_IMPCHas | 0.533363228 | 55 |
| SDF | 0.533441297 | 46 |
| Adap_comp_sub | 0.533452947 | 56 |
| DUF1776 | 0.533611083 | 32 |
| Prefoldin | 0.534767678 | 56 |
| Uso1_p115_head | 0.535137976 | 52 |
| PhetRS_B1 | 0.535346066 | 55 |
| TehB | 0.535447563 | 30 |
| Rio2_N | 0.535455652 | 55 |
| PPP5 | 0.535478766 | 53 |
| PIN_6 | 0.535555548 | 56 |
| DASH_Spc19 | 0.536163215 | 43 |
| efThoc1 | 0.536310307 | 56 |
| Autophagy_C | 0.53647199 | 53 |
| PGM_PMM_II | 0.536509161 | 55 |
| Thg1 | 0.537460582 | 55 |
| HAUS4 | 0.537987025 | 38 |
| MT | 0.538433562 | 56 |
| NUC153 | 0.538439227 | 56 |
| RNA_pol_Rpb2_6 | 0.538957483 | 56 |
| GDC-P | 0.539276122 | 55 |
| RNA_pol_Rpb2_7 | 0.540035246 | 56 |
| Peptidase_S9 | 0.540379804 | 56 |
| PRP1_N | 0.540401984 | 54 |
| NPR3 | 0.540458304 | 54 |
| Smg4_UPF3 | 0.540861965 | 56 |
| HSCB_C | 0.541590054 | 55 |
| Mre11_DNA_bind | 0.542014201 | 55 |
| Peptidase_M1_N | 0.542211788 | 56 |
| Nsp1_C | 0.542392408 | 52 |
| PNK3P | 0.542910349 | 54 |
| CtaG_Cox11 | 0.543490853 | 56 |
| RRP36 | 0.544162861 | 53 |

| | | |
|---|---|---|
| MMADHC | 0.544462876 | 42 |
| MIF4G_like_2 | 0.544520563 | 56 |
| ATP-grasp_3 | 0.545811086 | 47 |
| Semialdhyde_dh | 0.545881421 | 56 |
| Med18 | 0.545923505 | 47 |
| ASF1_hist_chap | 0.546054231 | 55 |
| MMgT | 0.54609033 | 49 |
| Ferrochelatase | 0.546543529 | 55 |
| Vac14_Fab1_bd | 0.54658003 | 56 |
| PI3K_C2 | 0.546609145 | 48 |
| AMPKBI | 0.54770396 | 54 |
| Ribosomal_S21 | 0.547799661 | 54 |
| Thioredoxin_14 | 0.547843436 | 53 |
| TMEM223 | 0.548142004 | 47 |
| zinc_ribbon_16 | 0.548356553 | 50 |
| DUF3128 | 0.548382361 | 52 |
| Cytidylate_kin | 0.548572322 | 54 |
| RPN5_C | 0.548677485 | 54 |
| PMT | 0.548860201 | 56 |
| Nuf2 | 0.549194958 | 45 |
| tRNA-synt_1 | 0.549197581 | 56 |
| DUF775 | 0.549348205 | 55 |
| Trm112p | 0.549382325 | 55 |
| ATP_synt_H | 0.549495396 | 47 |
| FAD_binding_8 | 0.550096054 | 55 |
| MIF4G_like | 0.550100441 | 56 |
| Tuberin | 0.550255449 | 52 |
| ApbA | 0.550429893 | 55 |
| Asparaginase | 0.550650476 | 56 |
| TIM21 | 0.550675646 | 52 |
| WBS_methylT | 0.550701456 | 53 |
| Med8 | 0.550810082 | 39 |
| Med4 | 0.551116884 | 49 |
| GWT1 | 0.551432048 | 52 |
| Ribosomal_L27 | 0.551707957 | 55 |
| MDD_C | 0.552140871 | 56 |
| Ins_P5_2-kin | 0.552150163 | 51 |
| TRI12 | 0.552363473 | 56 |
| TUDOR | 0.552706591 | 56 |
| Pkr1 | 0.553001449 | 46 |
| tRNA-synt_1c | 0.553178329 | 56 |
| Gcd10p | 0.557323224 | 55 |
| Mog1 | 0.557740695 | 55 |
| Rep_fac-A_3 | 0.557914171 | 55 |
| ApbA_C | 0.558269038 | 56 |
| BPL_N | 0.558563503 | 51 |
| PGM_PMM_III | 0.55901275 | 55 |
| tRNA_synt_1c_R1 | 0.55965691 | 55 |
| Pyridox_oxase_2 | 0.562242466 | 50 |
| NPR2 | 0.562429825 | 56 |
| DUF2423 | 0.562534167 | 53 |
| U6-snRNA_bdg | 0.562860437 | 54 |
| DUF974 | 0.563058978 | 54 |
| NDK | 0.563059684 | 56 |
| DUF92 | 0.563661179 | 53 |
| Sec1 | 0.564402481 | 56 |
| RNase_HII | 0.564451403 | 56 |
| Vta1_C | 0.564500886 | 51 |
| APG6 | 0.564651027 | 55 |
| She9_MDM33 | 0.564859477 | 56 |
| Erv26 | 0.564961016 | 49 |
| MRP-L28 | 0.565019748 | 55 |
| AIRS_C | 0.565566863 | 56 |
| DUF2462 | 0.565989974 | 54 |
| SF3b10 | 0.566021104 | 52 |
| DUF1242 | 0.566144581 | 46 |
| DNA_primase_lrg | 0.566367295 | 55 |
| hGDE_N | 0.567173781 | 50 |
| ACT | 0.567390995 | 55 |
| SBDS_C | 0.56773414 | 53 |
| DUF2315 | 0.567901897 | 49 |
| Glyco_transf_24 | 0.568184148 | 53 |
| PGM_PMM_I | 0.568845192 | 56 |
| ANAPC4 | 0.569042775 | 48 |
| DUF2431 | 0.569442215 | 55 |
| PMI_typeI | 0.569958123 | 54 |
| Peptidase_S24 | 0.570216872 | 56 |
| Rav1p_C | 0.570310483 | 54 |
| Semialdhyde_dhC | 0.570649677 | 56 |
| Es2 | 0.570682378 | 55 |
| Thioredoxin_8 | 0.571207708 | 35 |
| SRR1 | 0.571531034 | 49 |
| tRNA-synt_1c_C | 0.571682353 | 56 |
| STT3 | 0.572043106 | 55 |
| Tti2 | 0.572586208 | 46 |
| Biopterin_H | 0.572688017 | 55 |
| HD_2 | 0.572771359 | 49 |
| Clp1 | 0.572804954 | 56 |
| Rrp15p | 0.57288291 | 53 |
| Alpha_adaptin_C | 0.572894905 | 54 |
| DEK_C | 0.572903254 | 56 |
| P16-Arc | 0.574220013 | 55 |
| NAD_binding_9 | 0.574569289 | 32 |
| Vma12 | 0.574755287 | 51 |
| Glycos_transf_4 | 0.574782538 | 53 |
| NatB_MDM20 | 0.574845859 | 56 |
| GPI2 | 0.575499549 | 53 |
| UPF0172 | 0.575738052 | 51 |

| | | |
|---|---|---|
| Tau95_N | 0.575845394 | 55 |
| DHC_N1 | 0.576085343 | 56 |
| IML1 | 0.57658766 | 56 |
| DNA_RNApol_7kD | 0.576649807 | 52 |
| RTC | 0.577128872 | 53 |
| GSHPx | 0.577159182 | 53 |
| GYF | 0.577618275 | 50 |
| B3_4 | 0.578229998 | 55 |
| CMS1 | 0.578463438 | 52 |
| Thg1C | 0.578543063 | 54 |
| Alpha-mann_mid | 0.579547616 | 55 |
| TAP42 | 0.579683901 | 55 |
| Pterin_4a | 0.579739458 | 53 |
| 5-FTHF_cyc-lig | 0.579809423 | 53 |
| AstE_AspA | 0.579907192 | 49 |
| RNA_POL_M_15KD | 0.580224271 | 52 |
| Glyco_hydro_17 | 0.580291236 | 50 |
| Josephin | 0.580906355 | 43 |
| IDO | 0.581534655 | 56 |
| Ribosomal_L34 | 0.582062828 | 51 |
| ATP-synt_F | 0.58291835 | 55 |
| Spermine_synt_N | 0.58419091 | 56 |
| SHMT | 0.584902631 | 56 |
| ORC_WH_C | 0.585637428 | 55 |
| SRP54_N | 0.585749777 | 56 |
| TFIID-31kDa | 0.586088064 | 56 |
| Ribosomal_L37 | 0.58667209 | 55 |
| UcrQ | 0.587274031 | 56 |
| Nfu_N | 0.587460323 | 55 |
| VPS28 | 0.587657534 | 54 |
| NLE | 0.587878733 | 55 |
| SPT2 | 0.588528058 | 53 |
| Ndc80_HEC | 0.588724815 | 51 |
| Nkap_C | 0.589876192 | 53 |
| Sas10 | 0.590264427 | 55 |
| TBCA | 0.590325132 | 54 |
| Mis12 | 0.590836589 | 50 |
| PTPLA | 0.591547209 | 56 |
| Vta1 | 0.59160695 | 56 |
| COPI_assoc | 0.592766381 | 50 |
| DNA_pol_B_exo2 | 0.592906738 | 52 |
| Prenylcys_lyase | 0.593401336 | 51 |
| LIAS_N | 0.593497574 | 54 |
| DHC_N2 | 0.593615969 | 56 |
| SRP54 | 0.593746581 | 56 |
| ATP-synt_E | 0.593769721 | 52 |
| NUC173 | 0.594510754 | 55 |
| Calreticulin | 0.596885429 | 55 |
| Indigoidine_A | 0.597140264 | 51 |
| PRP3 | 0.597301276 | 56 |
| Rep_fac_C | 0.597684531 | 56 |
| Nop53 | 0.59773156 | 54 |
| PTPS | 0.597876246 | 54 |
| PIG-L | 0.598479619 | 53 |
| PTPA | 0.598502316 | 56 |
| Chs7 | 0.598567532 | 56 |
| AAA_3 | 0.599681323 | 56 |
| MgsA_C | 0.59992815 | 53 |
| SIP1 | 0.601610469 | 50 |
| RNA_pol_I_A49 | 0.601729022 | 54 |
| CTK3_C | 0.602546403 | 51 |
| NifU | 0.603057375 | 55 |
| NDUF_B12 | 0.604908251 | 53 |
| PITH | 0.605469046 | 56 |
| MoeA_C | 0.605975593 | 49 |
| Nrap_D3 | 0.606443488 | 55 |
| Nrap_D5 | 0.606443488 | 55 |
| Nrap_D4 | 0.606443488 | 55 |
| FeS_assembly_P | 0.606706554 | 55 |
| Thr_synth_N | 0.606943523 | 54 |
| TFIIA | 0.607070569 | 54 |
| zf-SNAP50_C | 0.607260303 | 46 |
| B5 | 0.607585958 | 55 |
| 3HCDH_N | 0.607692349 | 56 |
| RNA_pol_A_bac | 0.608217176 | 56 |
| NAD_binding_5 | 0.609361265 | 55 |
| HHH_7 | 0.609663773 | 53 |
| Pyridoxal_deC | 0.610265724 | 56 |
| Ribosomal_S16 | 0.610425108 | 53 |
| Elongin_A | 0.612232383 | 44 |
| Myotub-related | 0.612384158 | 54 |
| zf-DNL | 0.612878611 | 54 |
| Endonuclease_5 | 0.615025552 | 50 |
| AAA_assoc_2 | 0.615312359 | 53 |
| zf-Nse | 0.615535196 | 55 |
| INCENP_ARK-bind | 0.615998343 | 43 |
| PTS_2-RNA | 0.616146914 | 54 |
| L51_S25_CI-B8 | 0.616755856 | 55 |
| TruD | 0.617088501 | 56 |
| zf-rbx1 | 0.617265229 | 56 |
| Erg28 | 0.61948756 | 54 |
| cwf21 | 0.619877918 | 53 |
| AAA_7 | 0.620309461 | 56 |
| MWFE | 0.620450745 | 53 |
| MRP-S28 | 0.623027593 | 51 |
| Cys_Met_Meta_PP | 0.623065302 | 56 |
| RRS1 | 0.623718803 | 56 |
| Asparaginase_C | 0.624409304 | 55 |

| | | |
|---|---|---|
| NMT | 0.62468592 | 55 |
| Sedlin_N | 0.625058839 | 53 |
| Iwr1 | 0.625302944 | 47 |
| YABBY | 0.625576176 | 43 |
| EnY2 | 0.62647924 | 41 |
| zf-C3HC4_5 | 0.626679685 | 52 |
| PMT_4TMC | 0.626835817 | 56 |
| RAI1 | 0.626936528 | 49 |
| CDC37_N | 0.627425824 | 56 |
| Prp19 | 0.62751111 | 55 |
| DNA_binding_1 | 0.628540936 | 53 |
| zf-RING_5 | 0.62855498 | 56 |
| Memo | 0.628717427 | 54 |
| SNRNP27 | 0.628815794 | 53 |
| RRN3 | 0.629242486 | 55 |
| OMPdecase | 0.62929228 | 55 |
| DUF1077 | 0.629550715 | 52 |
| Sof1 | 0.630383372 | 54 |
| GST_N | 0.630408616 | 56 |
| DUF3385 | 0.630664914 | 55 |
| AAA_6 | 0.631096014 | 55 |
| Spt4 | 0.631218917 | 48 |
| MFS_MOT1 | 0.631506687 | 51 |
| SGTA_dimer | 0.632671211 | 48 |
| TFIIIC_sub6 | 0.634389322 | 47 |
| AIRC | 0.634486321 | 55 |
| PurK_C | 0.634486321 | 55 |
| Thymidylat_synt | 0.635428644 | 55 |
| Nrap_D6 | 0.63603374 | 55 |
| NAD_synthase | 0.636614206 | 56 |
| Sod_Cu | 0.636901363 | 56 |
| Paf1 | 0.637100147 | 48 |
| E1_4HB | 0.637160209 | 55 |
| Glycohydro_20b2 | 0.637245796 | 56 |
| Tho2 | 0.637769547 | 54 |
| Scramblase | 0.637910149 | 49 |
| Slu7 | 0.638371926 | 54 |
| Dfp1_Him1_M | 0.638703354 | 49 |
| AIRS | 0.638797889 | 56 |
| Cmc1 | 0.641377657 | 55 |
| DUF962 | 0.643388712 | 56 |
| GatB_Yqey | 0.643429728 | 56 |
| MafB19-deam | 0.643644314 | 56 |
| FDX-ACB | 0.643906498 | 50 |
| Ribophorin_II | 0.64405608 | 53 |
| ATG16 | 0.64491755 | 48 |
| HMG_CoA_synt_N | 0.644951686 | 54 |
| Helicase_C_3 | 0.645077421 | 54 |
| CLP1_P | 0.645743694 | 56 |
| RIX1 | 0.646083256 | 54 |
| TAFII55_N | 0.646122433 | 53 |
| rRNA_processing | 0.647000713 | 49 |
| Rad52_Rad22 | 0.647041477 | 53 |
| Mod_r | 0.647079486 | 50 |
| Peptidase_M28 | 0.647097013 | 56 |
| LIDHydrolase | 0.647997325 | 54 |
| UPF0004 | 0.648024878 | 48 |
| Nop16 | 0.648660922 | 55 |
| Tyr_Deacylase | 0.649608347 | 52 |
| VRR_NUC | 0.65046962 | 47 |
| DUF2034 | 0.650553581 | 44 |
| DNA_methylase | 0.651673736 | 56 |
| Tfb2_C | 0.65277707 | 56 |
| Nrap | 0.654930341 | 55 |
| DUF4097 | 0.655451666 | 44 |
| LZ3wCH | 0.655640543 | 53 |
| Nefa_Nip30_N | 0.656069067 | 51 |
| tRNA-synt_2c | 0.656781007 | 56 |
| Svf1_C | 0.657072621 | 53 |
| Alg14 | 0.65856535 | 50 |
| CASP_C | 0.658633388 | 45 |
| Eaf7 | 0.659035373 | 44 |
| RF-1 | 0.659725233 | 56 |
| PCNA_N | 0.659796562 | 56 |
| TBCC | 0.659988366 | 51 |
| DNA_pol_E_B | 0.660431565 | 56 |
| OST3_OST6 | 0.661474037 | 56 |
| Romo1 | 0.661943722 | 50 |
| Formyl_trans_C | 0.662019561 | 46 |
| Fer2 | 0.66253747 | 56 |
| AAA_8 | 0.662538045 | 56 |
| Dynein_AAA_lid | 0.662538045 | 56 |
| PCRF | 0.662717525 | 55 |
| RRF | 0.666023701 | 55 |
| E1_UFD | 0.666452148 | 55 |
| PFU | 0.666743665 | 55 |
| SBDS | 0.666779265 | 55 |
| MRP-L27 | 0.66708192 | 54 |
| Anticodon_1 | 0.668179986 | 56 |
| Utp11 | 0.669719709 | 56 |
| Vps23_core | 0.673611368 | 44 |
| MMR_HSR1_C | 0.673723794 | 21 |
| zf-DBF | 0.674258372 | 52 |
| Cu-oxidase | 0.675323559 | 51 |
| DUF2040 | 0.675978498 | 51 |
| DUF1748 | 0.676106978 | 54 |
| GCV_T | 0.677017885 | 56 |
| Hus1 | 0.677342113 | 52 |

| | | |
|---|---|---|
| Autophagy_act_C | 0.677389941 | 56 |
| RNA_pol_Rpb5_C | 0.677614192 | 56 |
| UPF0061 | 0.67823799 | 52 |
| DUF1744 | 0.678673133 | 53 |
| CLP_protease | 0.67872734 | 56 |
| Tfb2 | 0.680408612 | 56 |
| MCM_lid | 0.680409568 | 56 |
| ESCRT-II | 0.680480478 | 55 |
| PIG-X | 0.680801373 | 53 |
| Raptor_N | 0.680835232 | 53 |
| SURF1 | 0.68123368 | 54 |
| Zn_dep_PLPC | 0.681248095 | 43 |
| GatB_N | 0.681367119 | 55 |
| HAT_KAT11 | 0.681734554 | 54 |
| 3HCDH | 0.682734663 | 56 |
| Cu-oxidase_2 | 0.682881182 | 54 |
| Ribosomal_L35p | 0.683832022 | 46 |
| SecE | 0.684669047 | 55 |
| Peptidase_M16_C | 0.685685151 | 56 |
| MAPKK1_Int | 0.685741293 | 51 |
| RNA_pol_A34 | 0.686703639 | 54 |
| RNA_lig_T4_1 | 0.686722388 | 56 |
| Cid2 | 0.687514775 | 48 |
| Nop14 | 0.689905568 | 55 |
| NPL4 | 0.690382651 | 56 |
| BP28CT | 0.690560688 | 51 |
| MPP6 | 0.691470331 | 51 |
| AAR2 | 0.692882519 | 53 |
| HPPK | 0.693071232 | 56 |
| TMEM208_SND2 | 0.693307604 | 52 |
| MRP-S25 | 0.69341042 | 56 |
| Pet191_N | 0.693449101 | 46 |
| UNC-93 | 0.693664765 | 56 |
| N2227 | 0.693799356 | 55 |
| PPI_Ypi1 | 0.694193465 | 50 |
| Nup96 | 0.694998828 | 45 |
| FMN_red | 0.696125697 | 56 |
| MCM_OB | 0.696205162 | 56 |
| Ctf8 | 0.696720784 | 46 |
| Ran_BP1 | 0.697702468 | 56 |
| LDcluster4 | 0.69866442 | 50 |
| NAPRTase | 0.700022252 | 55 |
| DIM1 | 0.701831969 | 54 |
| Peptidase_S10 | 0.702215641 | 56 |
| AOX | 0.703148932 | 50 |
| DCB | 0.703362626 | 56 |
| P21-Arc | 0.704147204 | 54 |
| Arg_tRNA_synt_N | 0.705219006 | 52 |
| SUA5 | 0.70671238 | 56 |
| NGP1NT | 0.706735173 | 55 |
| Profilin | 0.706771151 | 56 |
| ATG27 | 0.707485068 | 55 |
| Med14 | 0.707539207 | 54 |
| NDUF_B8 | 0.708375458 | 47 |
| SelR | 0.709264675 | 55 |
| zf-NPL4 | 0.710210857 | 56 |
| UTP25 | 0.710456197 | 56 |
| TFIIS_C | 0.711219436 | 56 |
| Formyl_trans_N | 0.711952299 | 55 |
| SRP68 | 0.711971352 | 55 |
| Vps55 | 0.712519671 | 52 |
| DNA_pol_D_N | 0.712864778 | 55 |
| Ribosomal_L32p | 0.712900893 | 53 |
| SLX9 | 0.712994884 | 53 |
| Dcc1 | 0.713056625 | 51 |
| SH2_2 | 0.714216391 | 53 |
| C-C_Bond_Lyase | 0.715600606 | 47 |
| AKAP7_NLS | 0.71572424 | 45 |
| Nnf1 | 0.718136594 | 48 |
| TRAM_LAG1_CLN8 | 0.718387316 | 56 |
| Aquarius_N | 0.718582313 | 54 |
| Mnd1 | 0.718945555 | 51 |
| E1_FCCH | 0.720347819 | 55 |
| ESSS | 0.723730285 | 56 |
| TMEM234 | 0.724802408 | 49 |
| eIF3_p135 | 0.72518023 | 56 |
| Peroxin-3 | 0.725828439 | 55 |
| DWNN | 0.726784949 | 52 |
| Nrap_D2 | 0.727373548 | 55 |
| AAA_lid_11 | 0.727428205 | 55 |
| NMT_C | 0.727778181 | 55 |
| PHP | 0.729545549 | 55 |
| Cwf_Cwc_15 | 0.729961897 | 51 |
| DNA_primase_S | 0.730078397 | 53 |
| Mago-bind | 0.731284437 | 36 |
| SSrecog | 0.732876803 | 54 |
| eIF-3_zeta | 0.733262199 | 56 |
| AlaDh_PNT_C | 0.7341324 | 39 |
| ELP6 | 0.735000252 | 55 |
| DUF4208 | 0.73564084 | 50 |
| Trp_syntA | 0.73738955 | 56 |
| zf-4CXXC_R1 | 0.737442526 | 20 |
| GIY-YIG | 0.737909188 | 41 |
| CHD5 | 0.738600067 | 53 |
| zf-UBP_var | 0.740156482 | 54 |
| TF_Zn_Ribbon | 0.742800449 | 55 |
| E2_bind | 0.742985333 | 49 |
| RNA_pol_Rpb5_N | 0.743642569 | 55 |

| | | |
|---|---|---|
| ADIP | 0.743814166 | 46 |
| KIX_2 | 0.746167114 | 24 |
| YqeY | 0.7463722 | 53 |
| FolB | 0.747550241 | 56 |
| DXP_synthase_N | 0.748568232 | 30 |
| Plus-3 | 0.74961743 | 54 |
| Vps36_ESCRT-II | 0.749649398 | 47 |
| MCM | 0.752584932 | 56 |
| Spindle_Spc25 | 0.753236064 | 46 |
| CAS_CSE1 | 0.75333098 | 55 |
| POB3_N | 0.753395431 | 54 |
| Surp | 0.755361009 | 56 |
| Pcc1 | 0.756092188 | 48 |
| HMG_CoA_synt_C | 0.756485236 | 54 |
| DUF5110 | 0.758095551 | 39 |
| Glyco_hydro_20 | 0.760109071 | 56 |
| PIG-U | 0.760914494 | 55 |
| DCP1 | 0.761601541 | 50 |
| COX17 | 0.761650161 | 49 |
| Gpi16 | 0.763129031 | 56 |
| Nucleoside_tran | 0.763246421 | 55 |
| HVSL | 0.765061589 | 48 |
| ABC2_membrane_3 | 0.765214242 | 34 |
| SPC12 | 0.765532969 | 48 |
| zf-C3H2C3 | 0.766740953 | 35 |
| ABA_GPCR | 0.766993208 | 51 |
| Nodulin-like | 0.767518002 | 40 |
| P5-ATPase | 0.767593824 | 50 |
| B12-binding | 0.76833077 | 48 |
| mRNA_triPase | 0.768671302 | 56 |
| Pterin_bind | 0.768791917 | 56 |
| VPS11_C | 0.768798854 | 35 |
| Sec8_exocyst | 0.768861865 | 54 |
| CoaE | 0.769171688 | 56 |
| GalKase_gal_bdg | 0.770518639 | 53 |
| PHO4 | 0.771202895 | 45 |
| Mcl1_mid | 0.772358957 | 55 |
| SDA1 | 0.77296767 | 55 |
| Ribosomal_L28 | 0.773465291 | 53 |
| zf-Mss51 | 0.774698971 | 55 |
| PDT | 0.775977559 | 55 |
| vATP-synt_AC39 | 0.777695408 | 56 |
| PROCN | 0.778777954 | 55 |
| RNA_pol_Rpc4 | 0.77988539 | 56 |
| MDM31_MDM32 | 0.780037792 | 51 |
| CLASP_N | 0.780587399 | 56 |
| KTI12 | 0.78208215 | 53 |
| Val_tRNA-synt_C | 0.783203523 | 20 |
| APG9 | 0.787411656 | 55 |
| Cytochrom_B561 | 0.787612104 | 49 |
| BLOC1_2 | 0.788372449 | 45 |
| DUF423 | 0.789640183 | 52 |
| Med31 | 0.790030298 | 49 |
| zf-NOSIP | 0.790853078 | 51 |
| MOZART1 | 0.791251935 | 43 |
| FbpA | 0.793434101 | 54 |
| GCV_T_C | 0.794887155 | 53 |
| TFIIE_alpha | 0.795977395 | 44 |
| Flavodoxin_2 | 0.798404721 | 53 |
| Peptidase_C48 | 0.800201109 | 56 |
| Nbl1_Borealin_N | 0.800622602 | 37 |
| DPM2 | 0.802430884 | 49 |
| TAF6_C | 0.803238355 | 49 |
| TPMT | 0.805329593 | 53 |
| DUF4602 | 0.806298018 | 24 |
| Sec6 | 0.807947325 | 56 |
| RNase_H2_suC | 0.808998586 | 48 |
| Ribonuc_2-5A | 0.81189488 | 46 |
| Pox_MCEL | 0.812366828 | 56 |
| hDGE_amylase | 0.814446477 | 53 |
| Voltage_CLC | 0.816466036 | 56 |
| TAFII28 | 0.816985561 | 51 |
| NFACT-C | 0.817265843 | 54 |
| tRNA_int_endo | 0.818071775 | 50 |
| TFIIB | 0.81851411 | 56 |
| GIDA_assoc | 0.819286032 | 56 |
| IKI3 | 0.820871485 | 55 |
| Ceramidase | 0.821098266 | 51 |
| EST1 | 0.824551443 | 48 |
| Rsa3 | 0.826248596 | 49 |
| Glyco_hydro_18 | 0.827624104 | 56 |
| Cyclin_C | 0.828342967 | 56 |
| CKS | 0.829060291 | 50 |
| Yippee-Mis18 | 0.830561064 | 53 |
| Sec3_C | 0.832129719 | 56 |
| PTH2 | 0.832170766 | 56 |
| Cyto_heme_lyase | 0.833311496 | 55 |
| Nop52 | 0.836003851 | 53 |
| SF1-HH | 0.836175049 | 50 |
| DASH_Dad1 | 0.837878602 | 44 |
| Pkinase_fungal | 0.83964305 | 56 |
| Det1 | 0.842371047 | 31 |
| LTV | 0.844614756 | 53 |
| Use1 | 0.846557405 | 50 |
| FancD2 | 0.849995861 | 54 |
| Mob1_phocein | 0.853762375 | 55 |
| Inhibitor_I78 | 0.854053554 | 39 |
| Cupin_1 | 0.859472964 | 30 |

| | | |
|---|---|---|
| DS | 0.859997322 | 56 |
| Med1 | 0.865994186 | 54 |
| EFP_N | 0.867232359 | 41 |
| Peptidase_M48_N | 0.869160002 | 51 |
| Vps16_C | 0.87053553 | 56 |
| Mg_chelatase | 0.870796902 | 56 |
| Rad21_Rec8_N | 0.873587857 | 56 |
| His_biosynth | 0.874236582 | 56 |
| zf-LYAR | 0.875080669 | 49 |
| PCC_BT | 0.875969564 | 49 |
| DUF384 | 0.876665865 | 47 |
| Sen15 | 0.878528229 | 37 |
| Ribosomal_L33 | 0.879837027 | 43 |
| Asparaginase_2 | 0.880138209 | 52 |
| PP-binding | 0.880796155 | 56 |
| Arylesterase | 0.882309636 | 51 |
| MRP-S34 | 0.885641079 | 38 |
| Pet127 | 0.888001994 | 50 |
| GLE1 | 0.888543504 | 54 |
| PNP_UDP_1 | 0.891501405 | 56 |
| Skp1_POZ | 0.892251813 | 56 |
| Tau95 | 0.892622532 | 48 |
| ELO | 0.895718402 | 56 |
| Peptidase_M16 | 0.90140572 | 56 |
| XRN_M | 0.90174123 | 44 |
| Bmt2 | 0.905580186 | 50 |
| POLO_box | 0.906517347 | 46 |
| Orn_DAP_Arg_deC | 0.908621159 | 53 |
| Ran-binding | 0.914684367 | 46 |
| MKT1_N | 0.914949226 | 41 |
| Lipid_DES | 0.924324302 | 55 |
| tRNA_synthFbeta | 0.924606565 | 55 |
| DUF1981 | 0.924704997 | 55 |
| PIG-S | 0.931979537 | 54 |
| EF_assoc_2 | 0.932096825 | 56 |
| Reprolysin | 0.934907523 | 31 |
| Vps4_C | 0.936077559 | 56 |
| Spermine_synth | 0.937112972 | 56 |
| PH_13 | 0.939730577 | 51 |
| zf-ANAPC11 | 0.944794778 | 56 |
| HgmA | 0.945550869 | 56 |
| IU_nuc_hydro | 0.945758042 | 40 |
| DUF5310 | 0.947065301 | 21 |
| INSIG | 0.948150315 | 20 |
| ERAP1_C | 0.949721477 | 56 |
| RNase_H | 0.952069362 | 44 |
| Dynein_C | 0.955363166 | 47 |
| Glyco_hydro_16 | 0.955618453 | 54 |
| MaoC_dehydratas | 0.956940418 | 56 |
| Mo25 | 0.958986678 | 55 |
| Ofd1_CTDD | 0.961874956 | 56 |
| DIE2_ALG10 | 0.966708008 | 52 |
| ELFV_dehydrog_N | 0.967861947 | 55 |
| Ion_trans_2 | 0.969558375 | 56 |
| PDZ_6 | 0.970421643 | 52 |
| DASH_Duo1 | 0.971945312 | 46 |
| ketoacyl-synt | 0.975334096 | 56 |
| MaoC_dehydrat_N | 0.975702264 | 53 |
| DOPA_dioxygen | 0.978868259 | 25 |
| Stn1 | 0.979352625 | 20 |
| UBD | 0.981295777 | 37 |
| DUF953 | 0.982361122 | 54 |
| Sua5_yciO_yrdC | 0.987158762 | 56 |
| Peptidase_S8 | 0.988647802 | 56 |
| UDPGT | 0.993542234 | 47 |
| Bax1-I | 0.997235163 | 50 |
| CHS5_N | 0.998520329 | 42 |
| YCII | 0.999576158 | 45 |
| Peptidase_S26 | 1.000641785 | 21 |
| Tctex-1 | 1.008615448 | 51 |
| SRP-alpha_N | 1.010617317 | 54 |
| DUF89 | 1.014027498 | 53 |
| DUF2015 | 1.020780252 | 54 |
| CIA30 | 1.022365022 | 54 |
| Ydr279_N | 1.023359204 | 36 |
| RTT107_BRCT_5 | 1.023465694 | 37 |
| Mannosyl_trans2 | 1.026917796 | 52 |
| PDZ_2 | 1.027582369 | 38 |
| Peptidase_M20 | 1.031189677 | 56 |
| Cob_adeno_trans | 1.037302743 | 47 |
| zf-CDGSH | 1.037964018 | 21 |
| Neugrin | 1.03901377 | 44 |
| DUF836 | 1.041851993 | 48 |
| EMC3_TMCO1 | 1.042928621 | 55 |
| DNA_pol_phi | 1.043453551 | 56 |
| Zw10 | 1.053653769 | 49 |
| Orn_Arg_deC_N | 1.054616471 | 54 |
| MOSC | 1.063960718 | 53 |
| MCD | 1.064956232 | 42 |
| MOSC_N | 1.069132286 | 52 |
| Glycos_transf_3 | 1.072657723 | 56 |
| HD_3 | 1.073440099 | 52 |
| MAPEG | 1.073762654 | 56 |
| Ints3 | 1.076122486 | 51 |
| TPR_MLP1_2 | 1.07813465 | 20 |
| STI1 | 1.10202334 | 45 |
| PDZ | 1.102785477 | 33 |
| Metallopep | 1.107895139 | 39 |

| | | |
|---|---|---|
| Glyco_hydro81C | 1.109525202 | 52 |
| Fes1 | 1.112938457 | 46 |
| Peptidase_M3 | 1.11324781 | 56 |
| Sod_Fe_N | 1.113725436 | 55 |
| His_Phos_2 | 1.131922505 | 56 |
| VID27 | 1.138615257 | 40 |
| MTCP1 | 1.139321826 | 26 |
| TPX2 | 1.146984559 | 21 |
| DAGAT | 1.149232243 | 52 |
| Methyltransf_3 | 1.151928896 | 56 |
| GT87 | 1.152473392 | 24 |
| LeuA_dimer | 1.154778298 | 44 |
| ArsA_ATPase | 1.156589015 | 56 |
| Inhibitor_I9 | 1.157759257 | 56 |
| AXE1 | 1.16277101 | 20 |
| MCD_N | 1.166538906 | 32 |
| GPHR_N | 1.17150611 | 41 |
| VID27_N | 1.175446647 | 38 |
| SUV3_C | 1.176142015 | 31 |
| Arv1 | 1.18106208 | 49 |
| VID27_PH | 1.183042297 | 38 |
| HpcH_HpaI | 1.19107294 | 55 |
| Glyoxalase_4 | 1.193947068 | 51 |
| Rsm1 | 1.194573112 | 33 |
| GCS | 1.204730607 | 54 |
| DJ-1_PfpI | 1.205564575 | 55 |
| Methyltransf_24 | 1.206916636 | 54 |
| Glyco_tran_28_C | 1.206927631 | 55 |
| SOR_SNZ | 1.207542542 | 54 |
| Pro_CA | 1.211315156 | 56 |
| Skp1 | 1.214499797 | 55 |
| Peptidase_S41 | 1.21461227 | 41 |
| PTR2 | 1.21799302 | 56 |
| EBP | 1.218431726 | 51 |
| DMRL_synthase | 1.222094473 | 55 |
| Dicty_CAR | 1.231924239 | 26 |
| Glycos_trans_3N | 1.232417477 | 53 |
| Amidase | 1.246203193 | 56 |
| Med27 | 1.2488255 | 38 |
| Uricase | 1.262374234 | 53 |
| HSP20 | 1.264445183 | 47 |
| Glyco_hydro_81 | 1.292797151 | 52 |
| Inv-AAD | 1.315951621 | 54 |
| Rab5ip | 1.318519554 | 48 |
| Man-6-P_recep | 1.327864734 | 36 |
| Misat_Tub_SegII | 1.328366658 | 51 |
| Oxidored-like | 1.346324993 | 46 |
| BAAT_C | 1.348994142 | 21 |
| ATP_bind_3 | 1.371091624 | 55 |
| AA_permease_C | 1.381569992 | 34 |
| CsbD | 1.395939207 | 41 |
| Ald_Xan_dh_C2 | 1.416559095 | 25 |
| Sod_Fe_C | 1.419236716 | 55 |
| MIP | 1.420224416 | 54 |
| DSBA | 1.426412859 | 47 |
| Collagen | 1.449428386 | 21 |
| Lipase_3 | 1.511733888 | 56 |
| Acyl_transf_1 | 1.521242017 | 56 |
| RRM_3 | 1.523352195 | 28 |
| CLPTM1 | 1.541368428 | 55 |
| tRNA_lig_CPD | 1.54389478 | 24 |
| zinc_ribbon_15 | 1.547812793 | 20 |
| Fer4_14 | 1.552376115 | 23 |
| tRNA_lig_kinase | 1.559692616 | 38 |
| Trypsin_2 | 1.570845718 | 55 |
| Cohesin_load | 1.575003441 | 21 |
| Ketoacyl-synt_C | 1.578250927 | 56 |
| Ald_Xan_dh_C | 1.582928378 | 25 |
| CO_deh_flav_C | 1.584487477 | 25 |
| Mis14 | 1.5872813 | 26 |
| MFS_3 | 1.596009147 | 36 |
| CytochromB561_N | 1.607010832 | 29 |
| MeaB | 1.631354121 | 39 |
| zf-LITAF-like | 1.639378316 | 49 |
| MatE | 1.640484779 | 56 |
| GILT | 1.642619988 | 35 |
| zf-B_box | 1.647011447 | 27 |
| Ribonuclease_T2 | 1.665446876 | 52 |
| Symplekin_C | 1.673612117 | 36 |
| Peptidase_M16_M | 1.674942994 | 50 |
| FAD_binding_5 | 1.68704022 | 26 |
| Hom_end_hint | 1.701470509 | 20 |
| TUG-UBL1 | 1.715809038 | 24 |
| Coiled-coil_56 | 1.718560237 | 25 |
| Fer2_2 | 1.74791132 | 26 |
| Nucleoporin_FG | 1.748243217 | 28 |
| Pre-SET | 1.75109645 | 44 |
| TAP_C | 1.758684745 | 24 |
| DAP3 | 1.778585472 | 29 |
| Hexapep_2 | 1.786784173 | 38 |
| Glyco_hydro_46 | 1.80352842 | 38 |
| X8 | 1.835399902 | 51 |
| Fas_alpha_ACP | 1.842783421 | 50 |
| MM_CoA_mutase | 1.843722837 | 25 |
| Peptidase_S28 | 1.845830927 | 48 |
| CIMR | 1.872481578 | 24 |
| CAP59_mtransfer | 1.892227147 | 31 |
| SLC12 | 1.893294507 | 23 |

| | | |
|---|---|---|
| Mac | 1.897567607 | 28 |
| Cauli_VI | 1.898396912 | 34 |
| CENP-C_C | 1.906644151 | 23 |
| Peptidase_S9_N | 1.918813084 | 54 |
| zf-C2H2_3 | 1.931184951 | 23 |
| Swi5 | 1.949324324 | 24 |
| Acetyltransf_13 | 1.956118497 | 26 |
| Thiamine_BP | 1.984206452 | 22 |
| zf-CCCH_3 | 2.004746423 | 21 |
| Scm3 | 2.021107833 | 21 |
| EVE | 2.025364338 | 26 |
| PAP2_C | 2.04629385 | 46 |
| DUF4436 | 2.057785411 | 33 |
| Glyco_hydro_72 | 2.06942365 | 51 |
| PAF-AH_p_II | 2.0800353 | 21 |
| Acetyltransf_5 | 2.088693715 | 21 |
| DUF2418 | 2.226299048 | 28 |
| UPF0054 | 2.228120703 | 20 |
| Myb_DNA-bind_4 | 2.250020709 | 31 |
| 5_nucleotid_C | 2.251564784 | 46 |
| FPL | 2.278441378 | 20 |
| Condensation | 2.278744441 | 50 |
| dUTPase | 2.291668455 | 45 |
| CoA_trans | 2.294342603 | 21 |
| FAS_I_H | 2.299792206 | 49 |
| FAS_meander | 2.308898017 | 49 |
| ELYS | 2.330724858 | 28 |
| SAT | 2.3343447 | 49 |
| SLAC1 | 2.338162081 | 24 |
| DUF1729 | 2.342686373 | 50 |
| ISN1 | 2.407114231 | 27 |
| TRIC | 2.438105757 | 24 |
| Transferase | 2.449313946 | 49 |
| PI-PLC-Y | 2.572865476 | 28 |
| DUF2045 | 2.585329426 | 26 |
| FTP | 2.655802844 | 52 |
| BTHB | 2.694091701 | 22 |
| DUF4539 | 2.764707819 | 23 |
| DUF5598 | 2.798291097 | 25 |
| CAP | 2.884971048 | 55 |
| PI-PLC-X | 2.895790627 | 29 |
| SKN1 | 2.909746924 | 23 |
| TspO_MBR | 2.952665751 | 20 |
| Peptidase_M36 | 3.000396696 | 54 |
| SSXT | 3.076928935 | 23 |
| LPMO_10 | 3.534099451 | 28 |
| SGL | 3.858134411 | 23 |
| Trypsin | 4.405040113 | 54 |
| Tyrosinase | 5.76557213 | 23 |
| DUF3421 | 6.144143212 | 21 |
